# NitrOFF: An engineered fluorescent biosensor to illuminate nitrate transport in living cells

**DOI:** 10.1101/2025.03.22.644677

**Authors:** Mariah A. Cook, Jonathan D. Smailys, Ke Ji, Shelby M. Phelps, Jasmine N. Tutol, Wantae Kim, Whitney S. Y. Ong, Weicheng Peng, Caden Maydew, Y. Jessie Zhang, Sheel C. Dodani

## Abstract

The duality of nitrate is nowhere best exemplified than in human physiology – a detrimental pollutant but also a protective nutrient and signaling ion – particularly as connected to reactive nitrogen oxides. Aside from limited insights into nitrate uptake and storage, foundational nitrate biology has lagged. Genetically encoded fluorescent biosensors can address this gap with real-time imaging. However, imaging technologies for mammalian cell applications remain rare. Here, we set out to design and engineer a two-domain chimera fusing the split green fluorescent protein EGFP and the nitrate recognition domain NreA from *Staphylococcus carnosus*. Over 7 rounds of directed evolution, 15 mutations were accumulated resulting in the functional biosensor NitrOFF. NitrOFF has a high degree of allosteric communication between the domains reflected in a turn-off intensiometric response (*K*_d_ ≈ 9 µM). This was further reinforced by X-ray crystal structures of apo and nitrate bound NitrOFF, which revealed that the two domains undergo a large-scale conformational rearrangement that changes the relative positioning of the EGFP and NreA domains by 68.4°. Such a dramatic difference was triggered by the formation of a long helix at the engineered linker connecting the two domains, peeling the β7 strand off the EGFP and thus extinguishing the fluorescence upon nitrate binding. Finally, as a proof-of-concept, we highlighted the utility of this first-generation biosensor to monitor exogenous nitrate uptake and modulation in a human embryonic kidney (HEK) 293 cell line.

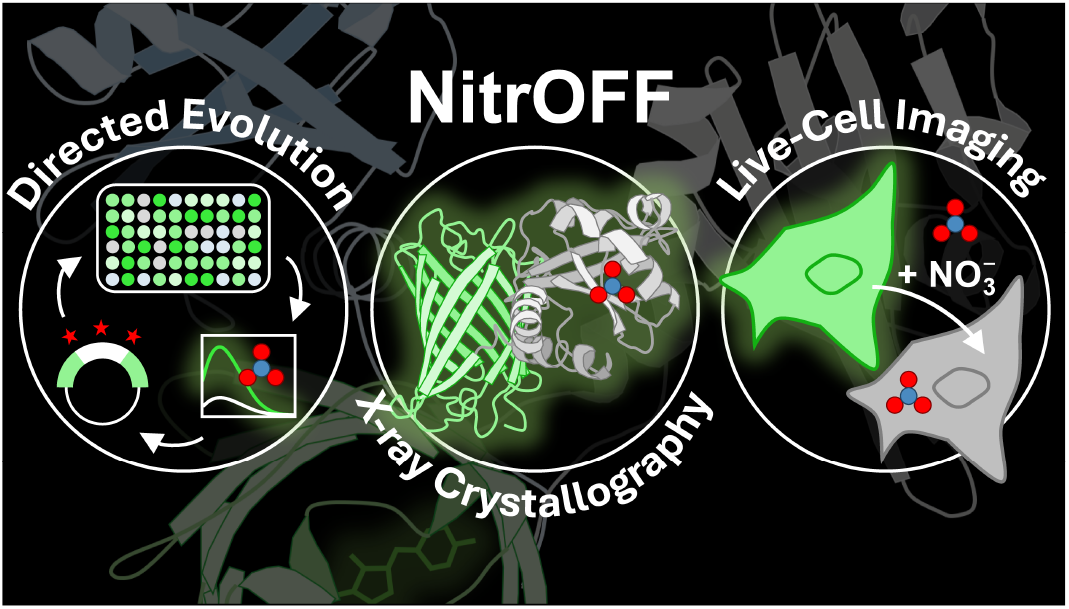

## Introduction

The role of the nitrate anion (NO_3_^-^) in human health is that of both friend and foe.^1^ As a ubiquitous environmental pollutant, nitrate was once considered to be detrimental to human health.^1^ This was due to its potential to form N-nitroso compounds, and its link to diseases such as methemoglobinemia.^1^ However over the last twenty years, nitrate has gained attention through its positive impact in cardiovascular health and exercise performance.^2^ These effects are brought about by its role as a stable reservoir for the signaling molecule nitric oxide.^3^ This reservoir is maintained by two sources: ingested nitrate from dietary sources and metabolized nitrate from the nitrate-nitrite-nitric oxide pathway.^4^ Current evidence points to the skeletal muscle as the largest and primary reservoir of nitrate in the human body.^5–7^ The nitrate-nitrite-nitric oxide pathway and these nitrate reservoirs are promising therapeutic targets for cardiovascular disease, endothelial dysfunction, ischemia-reperfusion injuries, and gastric ulcers.^8,9^ While these effects are known on the organismal level, it remains largely unknown how these effects cascade from changes in nitrate dynamics on the cellular and sub-organellar levels.

Motivated by the unknowns of nitrate biology, we seek to illuminate the dynamic mobilization, regulation, and biomolecular targets of nitrate in mammalian cells. Towards this, we are engineering fluorescent imaging technologies as part of our larger program aimed at unraveling the roles of anions in biology for the betterment of human health.^10–14^ To date, traditional biochemical assays – including electrophysiology, radiolabeling, chromatography, and enzymatic/chemical assays (e.g., the Griess reaction) – have provided insights into the cellular uptake and storage of nitrate.^15,16^ In fact, known intracellular nitrate concentrations can vary from 5 µM up to 10 mM depending on the cell type and transporter profile.^17,18^ Along these lines, known transporters with nitrate permeability include aquaporin-6 (AQP-6), chloride channels (CLCs), the cystic fibrosis transmembrane conductance regulator (CFTR), the calcium-activated chloride channel (TMEM16), and various members of the solute carrier (SLC) transporter family.^19–22^ Chief among the SLC transporters are the sodium/iodide symporter (SLC5A5), sialic acid transporter (SLC17A5), and chloride/bicarbonate exchanger (SLC26A7).^23–25^

As a complement, researchers have also turned to genetically encoded biosensors to probe nitrate in a spatiotemporal manner in living cells using fluorescence microscopy. To date, several fluorescent protein (FP)-based biosensors for nitrate have been reported in the literature. These include (i) the intensiometric sCiNiS and YFP-H148Q/I152L, (ii) the ratiometric ClopHensor, and (iii) the Forester resonance energy transfer (FRET)-based FLIP-NT, NiMet3.0, NiTrac1, and sNOOOpy (Table S1).^26–32^ The majority of these biosensors have had a strong foothold in illuminating plant biology and, with the singular exception of sNOOOpy, they have yet to demonstrate potential applicability in mammalian systems. This is due to a variety of reasons including but not limited to weak binding affinity, narrow dynamic range, bulky size, and sensitivity to other biologically relevant anions such as nitrite (NO_2_^-^) and chloride (Cl^-^).^26–32^

As such, we worked to develop a user-friendly fluorescent biosensor tailored to study nitrate uptake in mammalian cells with a compact design and potential for evolvability – traits that have not been explored in currently available tools. Such features are exemplified by the single fluorescent protein biosensor design that affords relatively large signal-to-noise ratios at a single emission wavelength. In this design, such as that of the GCaMP biosensor for calcium, an analyte recognition domain is fused to a split FP.^33^ Typically, the recognition domain is inserted at a disordered region of the FP spatially near the chromophore known as the β-bulge (i.e., residues 145 – 148 in the enhanced green fluorescent protein, EGFP).^34,35^ Given this, conformational changes in the recognition domain upon analyte binding are required to allosterically modulate the chromophore environment, inducing changes in the fluorescence intensity output. Broad applicability and tailoring of the properties of these two-domain chimeras can be achieved through directed evolution as demonstrated for analytes ranging from calcium, citrate, glutamate, and lactate, among others.^36–40^ It is important to note that such a design has yet to be explored for nitrate. To this end, the challenge lies in identifying a recognition domain that undergoes a conformational change upon nitrate binding, of which there are a limited number of examples.^31,41–44^

Most recently, biophysical and *in silico* investigations conducted by our lab have identified the soluble nitrate sensory protein NreA from *Staphylococcus carnosus* as a viable target for biosensor design.^45^ NreA is a small recognition domain (~17 kDa) compared to those used in current biosensors (~49 – 106 kDa). ^26–32,46^ The nitrate ion is recognized by an oxyanion hole, coordinated by the backbone amides of residues L67, A68, and I97 as well as the indole amide of W45 (Figure S1). Nearby hydrophobic residues further shield the anion from solvent. As measured by isothermal titration calorimetry (ITC), nitrate binding to NreA is enthalpically driven, resulting in a suitable micromolar binding affinity and a negative heat capacity change. The latter is key for our design as it indicates that NreA can undergo a conformational change.^45,47^ This is further supported by our molecular dynamics (MD) simulations that capture coordinated motions across the protein, allowing for nitrate recognition and release (Figure S1).^45^ Here, we have leveraged these collective features to engineer a single fluorescent protein biosensor named NitrOFF.

### Design and *in vitro* characterization of a prototype biosensor – NreA-EGFP

To generate the prototype biosensor, NreA from *Staphylococcus carnosus* was inserted into the β-bulge of the enhanced green fluorescent protein (EGFP). Specifically, we drew inspiration from the design of ncpGCaMP6, a non-circularly permuted biosensor for calcium.^48^ The residues L2 to P155 of NreA were fused to N144 and N149 of EGFP, respectively. We refer to this construct as NreA-EGFP, and all residue identifiers and position numbers correspond to the full construct shown in Figure S2. While short sequences can be added to serve as flexible linkers between the two domains, we opted not to add additional residues due to the conformationally dynamic nature of the NreA termini.^45^ Here, we define the N-terminal linker as the last four residues of NreA (145 – 148) and the C-terminal linker as the last two residues of NreA and the following two residues of EGFP (297 – 300).

We first expressed and purified NreA-EGFP with a C-terminal polyhistidine tag from *Escherichia coli* to evaluate its response to nitrate (Figure S3). At pH 7, apo NreA-EGFP has two absorption maxima centered at 396 and 504 nm (Figure S4, S5). In line with split EGFP biosensors, these correspond to the phenol and phenolate forms of the chromophore.^49,50^ Excitation of the latter results in an emission maximum at 514 nm (Figure S4, S5). Titration with up to 800 mM sodium nitrate, but not chloride or gluconate, generates a concentration-dependent absorption change that shifts the chromophore equilibrium towards the phenol form, translating to fluorescence quenching (Figure S4 – S7).

Further fitting of the emission response to a single-site binding model reveals that the affinity (apparent dissociation constant, *K*_d_) for nitrate is 144.5 ± 29.1 mM (Figure S5). However, we have previously determined the *K*_d_ of NreA to be approximately (*ca*.) 4 µM at 25 °C using ITC, indicating the potential for higher affinity.^45^ Barring any technical differences between fluorescence spectroscopy and ITC, this difference could be correlated with a greater degree of conformational restraint in NreA upon anchoring the termini to the β-barrel of EGFP.^51^ While the nitrate binding affinity falls out of a physiologically relevant detection range, we were encouraged by this prototype design because nitrate binding could be registered distally to the chromophore. Thus, in turn, allosterically tuning the EGFP fluorescence. At first glance, one would not know where to begin to tailor the suitability of NreA-EGFP for in-cell applications by enhancing features such as sensitivity and brightness. However, such an endeavor is not insurmountable as exemplified by many split FP biosensors.^38,40,52^ For this, we harnessed the power of directed evolution (Figure 1A).^53,54^

**Figure 1.**
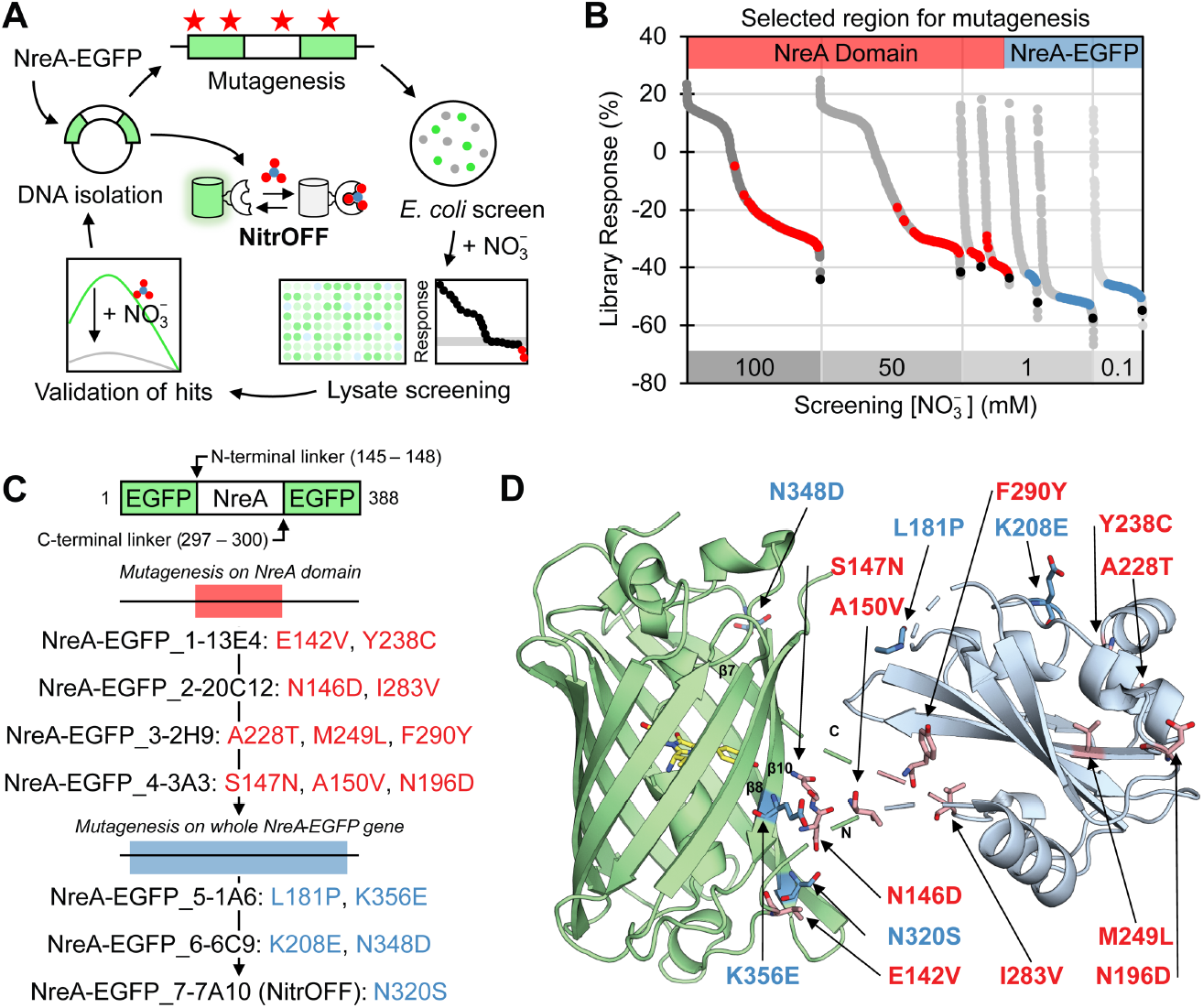
Directed evolution of NreA-EGFP to NitrOFF. (A) Protein engineering workflow. (B) Summary of library screening. For each round, variants shown as gray dots, corresponding to the screening concentration of sodium nitrate, are ranked from lowest to highest fluorescence quenching response defined as ΔF = ((F_f_-F_i_)/F_i_ × 100%) at pH 7.4 with sodium nitrate from left to right. The selected hit (black) (λ_ex_ = 485 nm, λ_em_ = 515 nm) was used as the new parent for the next round of mutagenesis. The red dots correspond to the libraries for the NreA domain only, and the blue dots correspond to the libraries for the entire gene. (C) Summary of mutations accumulated in each round of directed evolution on the NreA domain (red) or the whole gene (blue). (D) AlphaFold model of NitrOFF where the EGFP is in green, and the NreA domain is in light blue. The flexible regions connecting the domains with low confidence are shown as dashed lines. All mutations are labeled with the single letter amino acid code and position number from panel C. The N- and C-terminal regions of the NreA domain are labeled as N and C, respectively. The chromophore is shown in yellow.

### Directed evolution of NreA-EGFP

When considering both domains of NreA-EGFP, the β-barrel structure of EGFP is largely conformationally restricted, whereas the NreA domain is not. With this logic, we sought to enhance the coupling of the conformational change of NreA upon nitrate binding to the change in the EGFP fluorescence. We opted to introduce random mutations via error-prone polymerase chain reactions in two stages. First, we targeted the region spanning from D133 in the flexible loop region on the N-terminus through NreA to G298 on the C-terminus (Table S2, S3). To further fine-tune the interactions between the two domains, mutagenesis was extended to the entire NreA-EGFP gene (V2 – K390) (Table S2, S3).

We initially selected transformed *E. coli* colonies randomly, followed by those exhibiting green fluorescence (Figure S8 – S22). Eliminating dim variants not only reduced library screening size (~300 – 800 selected out of ~2,000 – 4,000 colonies per round) but also biased the selection towards variants with potentially improved expression and/or brightness. The selected variants were then expressed and screened in *E. coli* lysate in the absence (F_i_) and presence (F_f_) of sodium nitrate in 50 mM sodium phosphate buffer at pH 7.4 to determine response to nitrate (ΔF = (F_f_-F_i_)/F_i_ × 100%) (Figure 1B). During the rescreen process, the top variant was selected based on its improved response to nitrate as well as its baseline fluorescence in lysate, which could be due to improved expression or folding of the protein (Table S4, S5). As the quenching response improved (>10% difference between parent and variant), we increased the selection pressure by lowering the screening concentration of nitrate.

In the first stage of directed evolution across the NreA domain, the screening concentration of nitrate was shifted from 100 mM in the first round to 50 mM in the second round (Figure 1B). The top variant from the first round, with mutations E142V and Y238C, exhibited a ~13% improvement in the quenching response to 50 mM (ΔF = -36.7% ± 0.3%) compared to the NreA-EGFP parent (ΔF = -23.0% ± 0.2%) in the rescreen process (Figure 1C, Table S4). In the second round, the top variant with mutations at N146D and I283V exhibited a ~15% improvement in response to 1 mM (ΔF = -35.4% ± 0.1%) compared to the first round hit (ΔF = -20.0% ± 0.1%) (Figure 1C, Table S4). Hence, the screening concentration was lowered from 50 mM to 1 mM nitrate. With nominal gains in biosensor response to 1 mM nitrate in the third (ΔF = -38.3% ± 5.0%) and fourth (ΔF = -43.7% ± 5.2%) rounds, the mutagenesis region was expanded to include EGFP.

In the second stage of directed evolution, the screening concentration of nitrate was kept at 1 mM for two rounds (Figure 1B). In the sixth overall round, the second round over the whole gene, the top variant with mutations at K208E and N348D exhibited a ~13% improvement in the quenching response to 0.1 mM nitrate (ΔF = -52.8% ± 0.5%) compared to the fifth round hit (ΔF = -39.8% ± 2.0%) (Figure 1C, Table S5). With the screening concentration of nitrate lowered to 0.1 mM in the seventh round, we identified a variant that responded with a ~60% fluorescence quenching response in *E. coli* lysate (Table S5). This final variant was selected for further characterization and designated as NitrOFF (Figure 1C, 1D).

### Mutational Landscape of NitrOFF

Of the fifteen mutations that separate NitrOFF from NreA-EGFP, four were found in EGFP with the remaining eleven in the NreA domain. Most of the mutations accumulated either at the interface between the two domains or near the binding pocket (Figure 1D).^55,56^ The four mutations in EGFP near the interface include E142V in the flexible loop between β-strands 6 and 7, N320S on β-strand 8, and N348D and K356E on β-strand 10. Mutations in this region of EGFP are known to stabilize interactions with the fused recognition domain and even gate water entry from bulk solvent to chromophore environment.^35,50,57^

As seen with other split GFP biosensors, the composition of the linker regions connecting GFP and the recognition domain are critical for functional sensing due to chromophore proximity.^35^ Structural characterizations of these biosensors reveal that residues within the linker regions can interact with and stabilize the chromophore.^36,58–60^ In the case of NitrOFF, the N-terminal linker region accumulated several mutations while the second linker region at the C-terminus of NreA did not (Figure 1D). The N-terminal linker mutations, N146D and S147N, were found during the first stage of directed evolution on the NreA domain.

The substitution of N146 to the negatively charged aspartate appeared in the top three variants from the second round (Table S4). This critical mutation was responsible for increasing the dynamic range to allow for the screening concentration of nitrate to be reduced from 50 mM (~6% improvement from first-round hit) to 1 mM (~15% improvement) in the following rounds (Figure 1B; Table S4). Additionally, the substitution of S147 to the polar asparagine found in the fourth round may provide additional stabilization to the chromophore environment. Similar mutations of a linker residue – whether on the N- or C-terminus – to an asparagine residue have been observed to interact with the chromophore in the Citron biosensor for citrate and the K-GECO1 biosensor for calcium.^36,61^

Beyond the linkers, we looked to the mutations in the NreA domain itself. One of the first mutations was Y238C. Originally defined as the lid to the binding pocket, the Y238 residue (corresponding to Y95 in NreA) has been shown to affect binding affinity although it does not actively coordinate with the anion.^46^ Introduction of this mutation in combination with E142V led to the identification of the first-round hit with an ~13% improvement in the quenching response to 50 mM nitrate (Table S4). Like the N146D mutation, other mutations that appeared multiple times include L181P located in a flexible loop between two β-sheets in NreA and K208E located near the binding pocket (Table S5). From our MD simulations of NreA, we identified the R205 and K208 (R62 and K65 in NreA) as anchors to attract and guide anions into the binding pocket (Figure S1).^45^ Mutation of K208 to a negatively charged glutamate and Y238 to a smaller cysteine residue, in combination with other accumulated mutations, could alter local and global protein motions. Thereby, enhancing the coupling between the two domains as reflected in the observed sensitivity to nitrate in lysate.

### *In vitro* characterization of NitrOFF

To evaluate the effects of the accumulated mutations on the performance of NitrOFF, the biosensor was expressed and purified from *E. coli* with a C-terminal polyhistidine tag (Figure S23, S24). At pH 7, apo NitrOFF retains two absorption maxima centered at 396 nm (ε = 17,716 ± 120 M^-1^ cm^-1^) and 492 nm (ε = 31,542 ± 278 M^-1^ cm^-1^) with an isosbestic point at 430 nm, corresponding to the phenol and phenolate forms of the chromophore, respectively (Figure 2A, S25 – S28).^49,50^ Excitation of both maxima yields an emission consistent with the phenolate form at 512 nm (Figure S25). However, given the relative intensity differences, only the phenolate excitation was pursued (Φ = 0.60 ± 0.02; Figure S29, S30). Titration with up to 0.5 mM sodium nitrate generates a concentration-dependent response, tuning the chromophore equilibrium. This is seen as a shift in the absorption spectrum from the phenolate (ε = 6,187 ± 442 M^-1^ cm^-1^) to the phenol (ε = 29,970 ± 168 M^-1^ cm^-1^) form (Figure 2A, S27, S28). This translates to a *ca*. 77% fluorescence quenching (Φ = 0.73 ± 0.07) with no shift in the emission maximum (Figure 2B, Figure S29, S30). Indeed, the overall brightness (ε × Φ) decreases from 18.9 ± 0.7 to 4.5 ± 0.6 with the addition of sodium nitrate. While this tracks with the molar extinction coefficient, it is counter to the unexpected increase in the quantum yield – a rare, if not, unprecedented property for turn-off biosensors.

**Figure 2.**
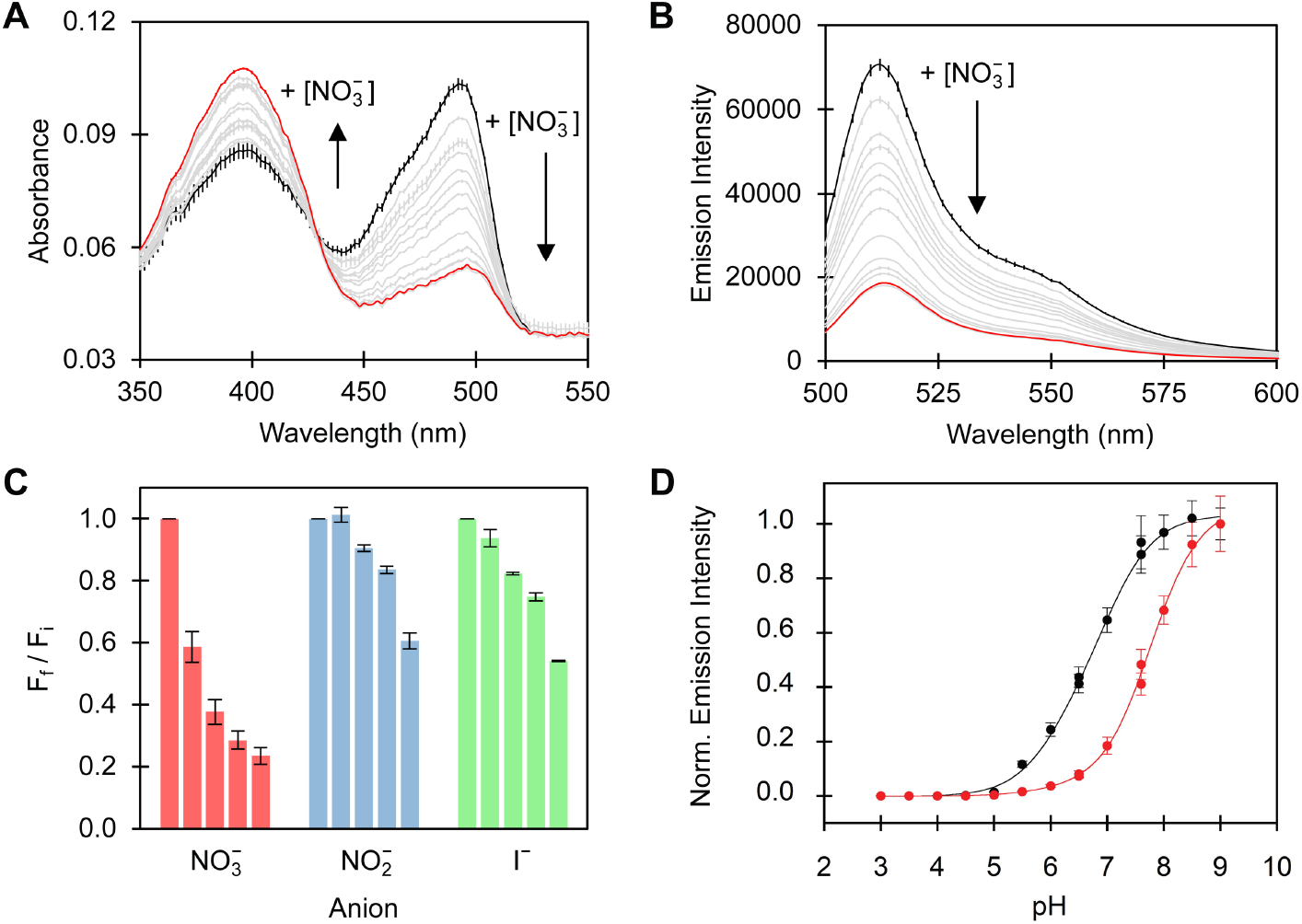
Spectroscopic characterization of NitrOFF. (A) Absorption and (B) emission spectra of 4 µM NitrOFF in the presence of 0 (black, bold), 2, 4, 6, 8, 10, 15, 25, 50, 75, 100, 200, 250 (gray), and 500 µM (red, bold) sodium nitrate at pH 7 (λ_ex_ = 480 nm, λ_em_ = 500 – 600 nm). For clarity, the average with standard error of the mean for three technical replicates with one protein batch is shown. All data for a duplicate protein batch is shown in the *Supporting Information*. (C) The fluorescence response of 4 µM NitrOFF in the presence of 0 (F_i_), 10, 50, 100, and 500 µM (F_f_) sodium nitrate (red), nitrite (blue), and iodide (green) at pH 7. (D) The normalized emission response of 4 µM NitrOFF at 512 nm in the absence (black circles) and presence of 500 µM (red circles) sodium nitrate from pH 3 to pH 9. The average with standard error of the mean for three technical triplicates across two protein batches is shown with the Henderson-Hasselbalch curve fittings.

Relative to the NreA-EGFP parent, NitrOFF boasts a 16,000-fold enhanced affinity for nitrate (*K*_d_ = 8.9 ± 0.6 µM) (Figure S26). Like the NreA domain, NitrOFF maintains a preference for nitrate over nitrite (*K*_d_ = 309.2 ± 19.2 µM; *ca*. 70% quench with 10 mM) and iodide (*K*_d_ = 136.1 ± 11.7 µM; *ca*. 61% with 10 mM) (Figure 2C, Figure S31, S32) by 35-fold and 15-fold, respectively.^45^ To account for ionic strength effects, the fluorescence response of NitrOFF was evaluated to high concentrations (10 – 100 mM) of sodium chloride or sodium gluconate (Figure S33, 34). While fluorescence quenching (*ca*. 10 – 30%) is observed, such effects are not unfounded with *in vitro* characterization of fluorescent proteins.^62,63^ Furthermore, in line with this design, the baseline emission and response to nitrate is pH-dependent.^35,50,51,60^ As can be seen in Figure 2D, NitrOFF maintains sensitivity, albeit to a varying extent, across a physiological range from pH 6 – 8 (Figure S35, S36). Fitting of these data links the sensing mechanism to a shift in the chromophore p*K*_a_ from 6.71 ± 0.04 to 7.61 ± 0.05 in the presence of nitrate.

### X-ray structures of NitrOFF

To probe if structural changes drive the observed sensing mechanism, we crystallized NitrOFF in the presence and absence of nitrate. NitrOFF was purified via a gel filtration column with 20 mM Tris buffer at pH 7.5 containing 150 mM sodium nitrate to generate the bound or OFF state (Figure S37). After screening and optimization, diffraction quality crystals were obtained at room temperature (RT) with 1.15 M ammonium sulfate at pH 5.4. Crystals diffracted to 3 Å resolution in the H3 space group. For molecular replacement, the structures of EGFP (PDB ID: 7VCM) and NreA (PDB ID: 6IZJ) were used as individual structural components to search for solutions.^51,64^ The final structure was determined with 18 NitrOFF molecules per asymmetric unit (ASU) with both EGFP and NreA domains well resolved (Figure 3A).

**Figure 3.**
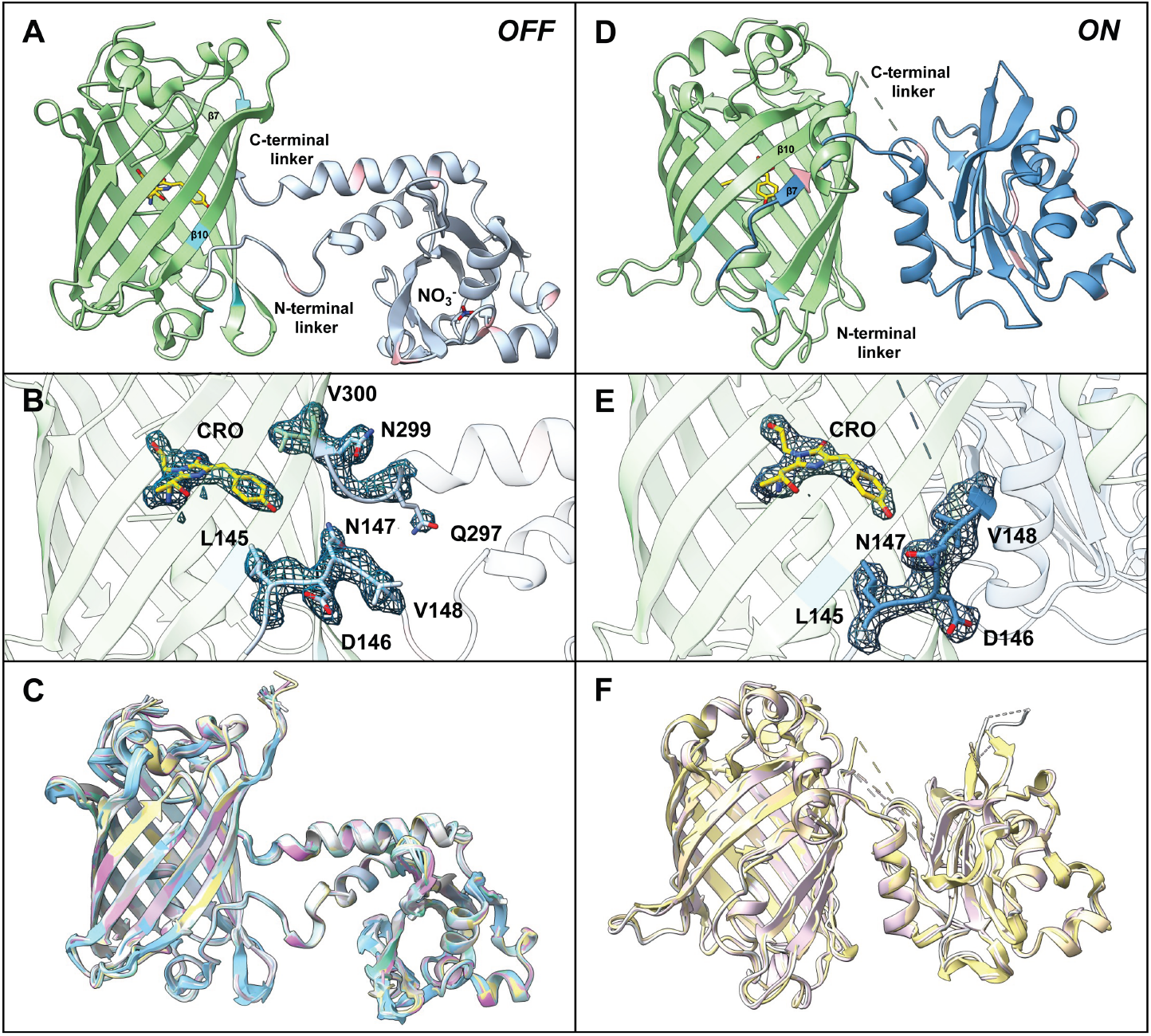
X-ray crystal structures of NitrOFF in the OFF and ON states. (A) General overview of the OFF state, with EGFP colored green, the NreA domain colored light blue and chromophore in yellow. Mutated positions in EGFP are colored in aquamarine and the NreA domain are in pink. (B) 2Fo-Fc electron density map overlay of the N- and C-terminal linkers for the OFF state shown in sticks, depicted at 2 σ. (C) Superimposition of the 18 molecules contained within the ASU in OFF state. (D) General overview of the ON state, with EGFP colored in green, and the NreA domain in dark blue. Mutated positions in the EGFP domain are colored in aquamarine and in the NreA domain are in pink. (E) 2Fo-Fc density map of the N- and C-terminal linkers for the ON state depicted at 2 σ. (F) Superimposition of the 4 molecules contained within the ASU of the ON state. Abbreviation: ASU, asymmetric unit.

With both domains identified, we observed strong positive and continuous electron density indicative of the linker residues connecting the two domains (N-terminal linker residues 145 – 148 and C-terminal linker residues 297 – 300; Figure 3B). We considered the possibility that the linker configurations might vary among the 18 molecules of the ASU. Thus, we manually built linker residues for each molecule independently. We then superimposed all molecules in the ASU to compare the relative positioning of the linkers connecting the split EGFP to NreA. Intriguingly, the 18 molecules are almost completely superimposable with an average root mean square deviation (RMSD) of 0.30 Å (Figure 3C). Considering the various local crystal packing environments for the molecules in the ASU, the high conservation of the relative orientation and the configuration of the linkers indicates that the linker regions of the OFF state assume the observed conformation without interference from crystal packing.

In parallel, we prepared the apo or ON state of NitrOFF using a gel filtration column buffer with 20 mM Tris buffer at pH 7.5 containing 150 mM sodium chloride (Figure S37). At RT, the protein crystallized with 1.7 M ammonium phosphate at pH 5.0 with crystals diffracting up to 3.0 Å in the space group of C2 2 2_1_, containing four molecules in the ASU (Figure 3D). Just like our procedure with the bound biosensor, we built the linkers between EGFP and NreA individually for each molecule of the ASU (Figure 3E). While the N-terminal linker for the sensor (residues 145–148) was well-resolved with a strong connected positive density (Figure 3E), we only observed scattered density close to the C-terminal linker (residues 292 – 309), indicative of high disorder (Figure 3F). Additionally, residues 179 – 183 in the NreA domain also exhibited high flexibility in all four copies of the molecules in the ASU. Nevertheless, the N-terminal linker (residues 145 – 148) where NreA was inserted into the split EGFP retained the same configuration in all 4 molecules of the ASU with clear electron density (Figure 3F).

### Nitrate binding site in the ON and OFF states of NitrOFF

With the structures of the two different states available, we examined the putative recognition site in the NreA domain to identify nitrate binding. In the OFF state structure, we observed strong positive density at the putative nitrate binding site (Figure 4A).^46^ The density is consistent with the incorporation of the nitrate ion (Figure 4A). Nitrate is placed in the oxyanion hole formed by the two short helices (residues 209 – 218 and 240 – 245). An extensive hydrogen bonding network forms between nitrate and surrounding residues, including backbone amide groups of L210, P239, and I240 (Figure 4B). Additional favorable interactions form between the indole nitrogen of W188 with the oxygen atom of nitrate (3.2 and 3.5 Å) (Figure 4B).

**Figure 4.**
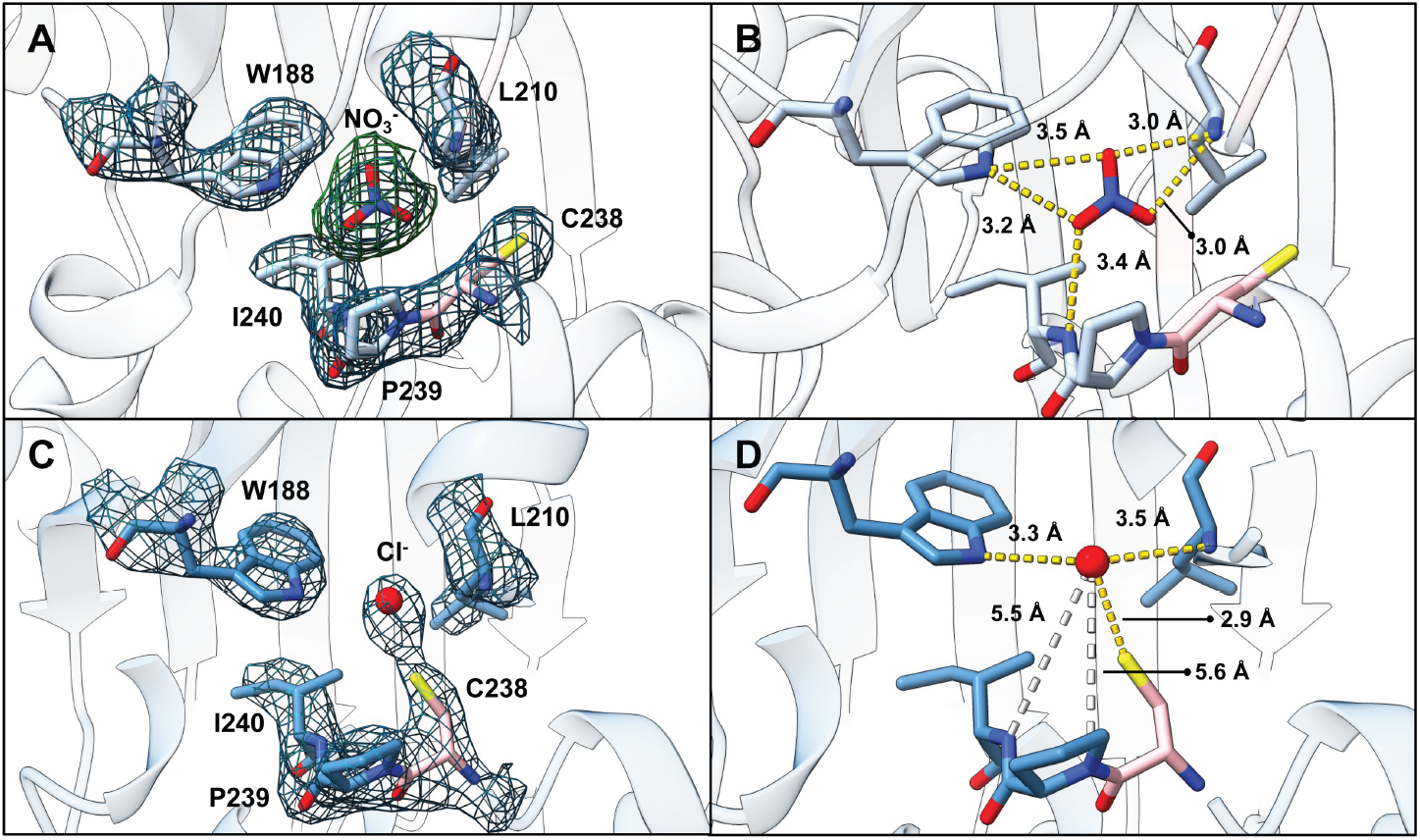
The interaction network at the NreA binding pocket in the OFF and ON states of NitrOFF. (A) 2Fo-Fc electron density map (contoured at 2 σ) of the residues (dark blue) pertinent to the formation of the hydrogen bond network coordinating nitrate in the OFF state. Omit Fo-Fc density map (contoured at 3 σ) at the nitrate binding site when nitrate was not included in the map calculation (green). The residues involved in nitrate binding are shown as sticks with carbon in sky blue except for cysteine in pink. (B) Hydrogen bonding interactions to nitrate in the OFF state are shown as yellow dashed lines with their corresponding distances. (C) ON state 2Fo-Fc electron density map (contoured at 2 σ) with the chloride ion shown in red. The analogous residues from panel A are shown. (D) Hydrophilic interactions are shown as yellow dashed lines, with white dashed lines representing bonds showing different distances of atoms involved in hydrogen bonding in panel B.

At first glance, the ON state NreA revealed little difference from the OFF state NreA at the nitrate binding site (RMSD of 0.8 Å) (Figure 4C). However, careful inspection reveals the lack of nitrate leads to a reduction of volume in the binding pocket at the oxyanion hole, allowing binding of smaller molecules such as a chloride ion (Figure 4D). Without the stabilization of the nitrate ion to the amide backbones within the nitrate binding pocket, the NreA domain in the ON state is more flexible compared to the OFF state (Figure 3D). The reduction of order in the absence of nitrate in this region increases the disorder in the proximal loop (residues 203 – 209), which has a connecting helix (residues 209 – 218) that forms part of the oxyanion hole (Figure 3D). The double helical knot that forms on the other side of the oxyanion hole also undergoes a shift in conformation in the absence of nitrate binding. Specifically, the engineered cysteine residue at position 238 has now rotated to partially occupy the binding pocket (Figure 4D). This twist of the cysteine side chain causes the unwinding of the short helix (residues 231 – 237) (Movie S1).

### Relative orientation of NreA and EGFP in the ON and OFF states

The most striking difference between the ON and OFF states is the relative orientation of the NreA domain to the EGFP domain, exhibiting a large conformational change (Figure 5A, Movie S2). To quantify this change, the EGFP domain from the two states was superimposed to measure the relative position of the domains. The planes defining the measurement were generated using the axis of the EGFP β-barrel and ion binding positions in the NreA domain, the resulting planes intersected at an angle of 68.4 ° (Figure 5B). This dramatic change is induced by a newly formed α-helix in the OFF state near the C-terminal linker (residues 279 – 296) (Figure 5B). This long helix of 22 Å acts as a spring and rigidifies the relative orientation of EGFP and NreA. In contrast, the same region in the ON state is much more flexible, adopting a short β-zipper (residues 279 – 291) and then becoming disordered from residues 292 to 307 (Figure 5B). The high flexibility of this region with the C-terminal linker provides no anchoring force to limit the orientation of EGFP and NreA (Movie S2). Notably, this rationalizes the unexpected higher quantum yield observed with NitrOFF bound to nitrate.

**Figure 5.**
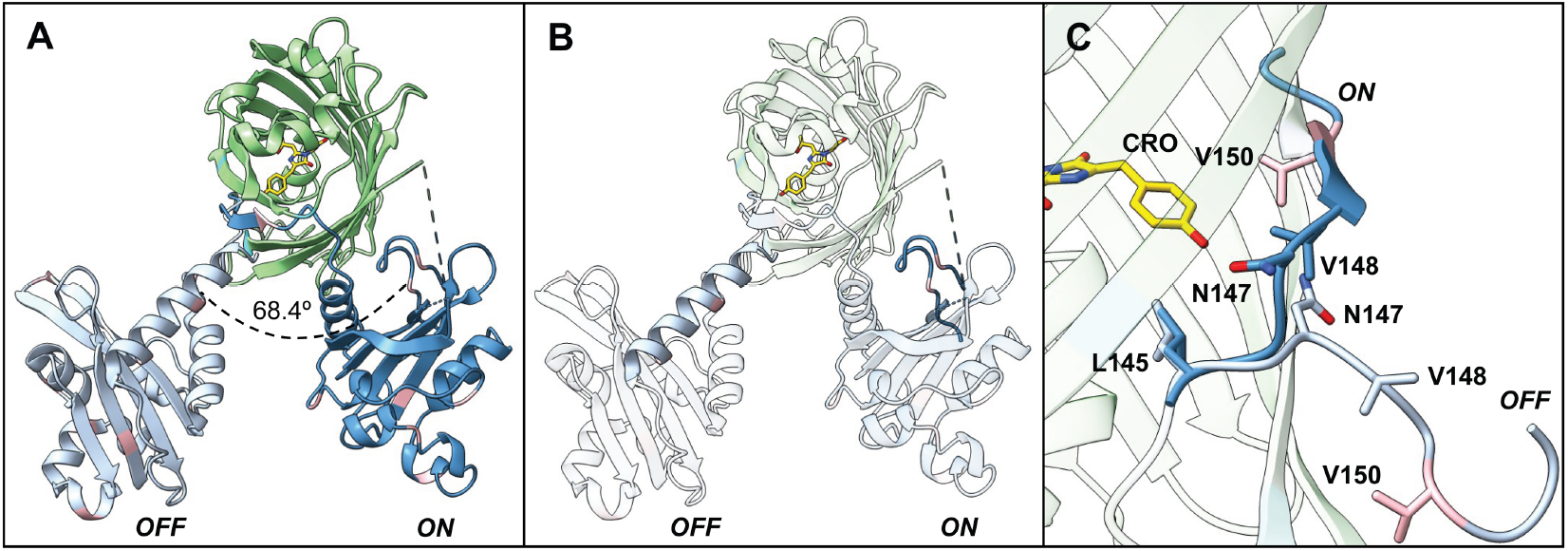
Conformational rearrangement between the OFF and ON states of NitrOFF. (A) Skyview of the two states, superimposed by the EGFP domains. The NreA domain undergoes a 68.4° change. (B) Skyview of the two states where the chromophore (yellow) and residues 279 – 291 (light or dark blue for the OFF and ON states, respectively) are opaque. (C) Zoomed in view of the superimposition for the N-terminal linker between the OFF (light blue) and ON (dark blue) states.

Intriguingly, the large conformational changes of the linker connecting EGFP and NreA upon nitrate binding changes the local environment of the chromophore (Figure 5C). The formation of the long helix in the OFF state generates a pulling force that peels part of the β7 strand from the core β-barrel of EGFP, turning the backbone 90° away and creating a gap in the β-barrel of the 7^th^ EGFP strand (Figure 5C). The breakage of the β7 strand exposes the typically shielded chromophore to the surrounding solvent, diminishing fluorescence. Several residues in the EGFP β-barrel for the OFF state adopt alternate conformations (Figure 5C). Specifically, engineered residue N147 flips almost 180°, altering the direction of the linker backbone by ~90° in the OFF state (Figure 5C and Movie S2). This rotation results in a gap formed by insertion of NreA, with extensive hydrophilic interactions involving N147, Q296, K315 and R317. The redirected linker further reorients the gate residue V148 and flips its hydrophobic side chain outwards, losing the hydrophobic lid covering the chromophore in the OFF state (Figure 5C). This is dramatically different from the ON state in which the EGFP β-barrel structure remains intact (Figure 5C).

Collectively, the structural differences observed between the OFF and ON states indicate how the turn-off response is produced. Without nitrate bound in the ON state, the NreA domain is flexible, and the linker regions do not pose any steric restriction. This allows for the adoption of the correct fold for the EGFP β-barrel while also maintaining the integrity of the hydrophobic chromophore environment. In contrast, the binding of nitrate in the OFF state rigidifies the NreA structure and promotes the formation of a long helical linker that exposes the chromophore to the bulk solvent, thus quenching the fluorescence emission (Figure 5C).

### In-cell validation of NitrOFF

Finally, we evaluated the utility of NitrOFF to monitor nitrate transport in mammalian cells with time-lapse fluorescence microscopy. For this proof-of-concept, we selected human embryonic kidney (HEK) 293 cells because of their propensity to express recombinant proteins.^65,66^ Moreover, specific to our goal, this cell line endogenously expresses anion-sensitive transporters including CLC-2, TMEM16A, and TMEM206 that are permeable to nitrate.^67–72^ While nitrate uptake has not been directly monitored via fluorescence imaging in HEK 293 cells, intracellular acidification upon treatment with millimolar concentrations of nitrate has served as an indirect proxy.^73^ Since the sensing mechanism of NitrOFF is connected to pH, we first determined if exogenously supplemented nitrate would induce a pH change, if at all, in our real-time anion exchange assay. For this, we turned to the excitation ratiometric fluorescent protein indicator for pH called RpHlourin2 (Table S7, S8, Figure S38).^74^ Briefly, transiently transfected cells were initially incubated and imaged in a phosphate-based buffer containing 137 mM sodium chloride. Following this, the cells were perfused with buffers containing increasing concentrations of sodium nitrate while maintaining the total ionic strength. The emission ratio change shows intracellular acidification with 0.5 mM sodium nitrate (1.18-fold ± 0.06) but not with 0.25 mM (1.03-fold ± 0.03) (Figure S39, S40, Movie S3 – S7). With this information in hand, we pursued an anion exchange assay with 0.25 mM sodium nitrate for NitrOFF. The gene encoding NitrOFF was codon optimized and expressed under a CMV promoter with no targeting sequences, resulting in a bright fluorescence signal in the cytosol and nucleus (Figure 6A; Figure S41). To test the response of the biosensor to chloride, the most biologically abundant anion, the cells were incubated and imaged in buffer containing 137 mM sodium chloride and then perfused with the same buffer (Figure 6B, S42, Movie S8 – S10).^75^ A similar trend in fluorescence response was observed when the cells were incubated in buffer containing 137 mM sodium gluconate to deplete the cells of labile chloride and then perfused with buffer containing 137 mM sodium chloride (Figure 6B, S43, S44, Movie S11 – S16). The large variability in fluorescence response (17.5% ± 0.1 quench) could be attributed to a combination of factors such as ionic strength and/or changes in cell shape and movement. Finally, following incubation and imaging in buffer containing 137 mM sodium chloride, the cells were perfused with buffer containing 0.25 mM sodium nitrate and 136.75 mM sodium chloride. As nitrate uptake occurred, the fluorescence signal quenched by 42.1% ± 0.1, indicating that NitrOFF can distinguish nitrate in the presence of chloride, despite their similarities in size and charge (Figure 6B, S45, Movie S17 – S19).^76^

**Figure 6.**
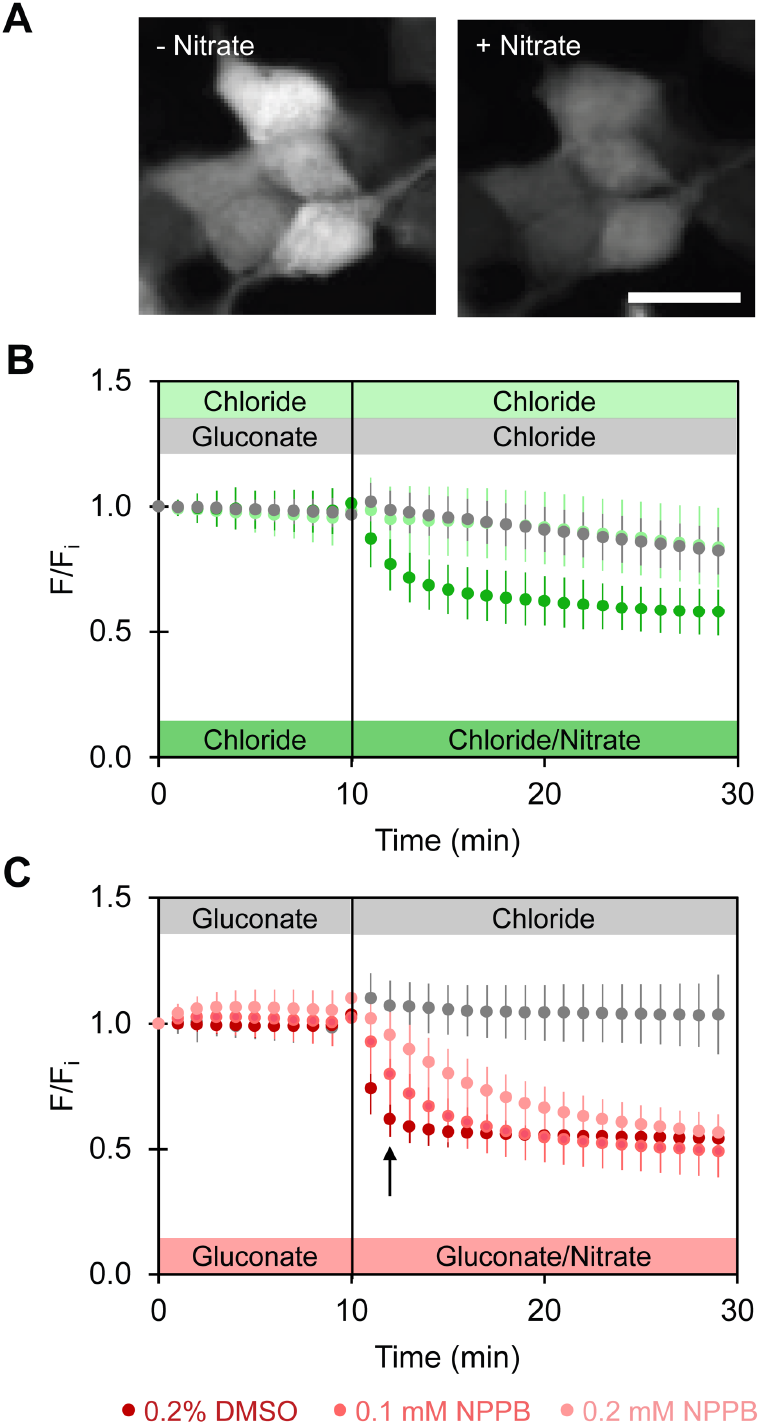
Fluorescence microscopy reveals that NitrOFF can monitor nitrate transport in HEK293 cells. (A) Representative snapshot of NitrOFF expressing cell before (left panel) and after (right panel) perfusion with 0.25 mM nitrate Scale bar: 20 µm. (B) The average median fluorescence response (F/F_i_) with standard deviation of NitrOFF from 3 biological replicates with *n* regions of interest for the exchange of 137 mM chloride with 137 mM chloride (light green dots, *n* = 624), 137 mM gluconate with 137 mM chloride (gray dots, *n* = 2384), and 137 mM chloride with 136.75 mM chloride/0.25 mM nitrate (dark green dots, *n* = 664). The start time of the perfusion to exchange the buffer is indicated by the vertical dashed line. (C) The average median fluorescence response (F/F_i_) with standard deviation of NitrOFF from 3 biological replicates with *n* regions of interest for the exchange of 137 mM gluconate with 136.75 mM gluconate/0.25 mM chloride (gray dots, *n* = 2129) and 137 mM gluconate with 136.75 gluconate/0.25 mM nitrate in the presence of 0.2% DMSO (dark red dots, *n* = 1414), 0.1 µM NPPB (red dots, *n* = 1823), or 0.2 µM NPPB (pink dots, *n* = 1483). The start time of the perfusion to exchange the buffer is indicated by the vertical dashed line. The largest change with NPPB is denoted by the black arrow. All experiments were carried out at pH 7.4, 37 °C. Abbreviation: DMSO, dimethyl sulfoxide; NPPB, 5-nitro-2-(3-phenylpropylamino)benzoic acid.

Motivated by these results, we finally probed if nitrate uptake could be affected by the modulation of endogenous transporters. In lieu of a dedicated nitrate transporting protein or small molecule modulator, we selected 5-nitro-2-(3-phenylpropylamino)-benzoic acid (NPPB), a broad-spectrum chloride channel inhibitor that targets CLC, TMEM16A, and TMEM206 mentioned above.^72,77–81^ In connection to nitrate, electrophysiology experiments in human submandibular gland cells and atrial myocytes show that nitrate uptake can be inhibited by NPPB.^24,82^ To test this, HEK 293 cells were first incubated in buffer containing 137 mM sodium gluconate to lower the competition by chloride for transporters, and then perfused with buffer containing 0.25 mM sodium chloride or nitrate and 136.75 mM sodium gluconate. Under this exchange assay, NitrOFF retained its response to nitrate (45.7% ± 0.1 quench) but not chloride (1.04-fold ± 0.02) (Figure 6C, S46 S47, Movie S20 – S25). The replacement of the DMSO vehicle control with NPPB decreased the nitrate uptake with a dose-dependent response to NPPB (Figure 6C, S48, S49, Movie S26 – S31). With respect to the control, the greatest difference in quenching was observed two minutes after the exchange (DMSO = 38.0% ± 0.1%; 0.1 mM NPPB = 20.0% ± 0.1%; 0.2 mM NPPB = 4.6 ± 0.1%) (Figure 6C). Moreover, intracellular pH remained unchanged in these assays (Figure S50 – S53, Movie S32 – S43). While the rate of nitrate uptake was affected, overall nitrate transport was not inhibited. This perhaps is not surprising given the functionally redundant pathways for nitrate entry, hinting at the essentiality of nitrate for biological function.^19–25^

## Conclusion

In summary, we have designed and developed NitrOFF, a new genetically encoded biosensor for nitrate based on the fusion of split EGFP and the nitrate recognition domain NreA. While the initial NreA-EGFP prototype is responsive to nitrate, its sensitivity precludes applicability in mammalian cells. Harnessing the power of directed evolution, over 7 rounds, 15 accumulated mutations dramatically tuned the allosteric communication between the two domains. In turn, translating to a turn-off intensiometric response with a measurable 16,000-fold enhancement in binding affinity. Furthermore, in a rare instance, X-ray crystal structures of both the apo and nitrate-bound forms of NitrOFF were resolved, illustrating the allosteric mechanism by which nitrate binding can be communicated to the chromophore.^83–86^ In the absence of nitrate, the C-terminal linker region is conformationally flexible, while both EGFP and NreA are well folded. Particularly, the key β7 strand of EGFP shields the chromophore from solvent, creating a microenvironment to stabilize the fluorescent state (ON state). However, the binding of nitrate induces rigidification of the C-terminal linker connecting NreA and EGFP, resulting in the formation of a long α-helix. Coupled with this, a 68.4° conformational shift invokes peeling of the β7 strand off the EGFP barrel, exposing the chromophore to bulk solvent and extinguishing the fluorescence (OFF state). Finally, as a proof-of-concept, NitrOFF was transiently encoded in the HEK 293 cell model and visualized using fluorescence microscopy. Our real-time anion exchange assays reveal that NitrOFF provides a direct readout of nitrate uptake and accumulation without any confounding artifacts from pH or ionic strength. Moreover, NitrOFF can be used to link nitrate transport through NPPB-sensitive chloride channels. To our knowledge, such a demonstration is unprecedented by fluorescence imaging with a biosensor, providing a rapid and scalable complement to traditional electrophysiology measurements.

As the toolmakers and end users of NitrOFF, we recognize its potential to illuminate nitrate across a spectrum of cellular processes and motivate end users to do the same. Building from our nitrate uptake assay, cryptic nitrate transporters and activity can be uncovered. From our perspective, such a pursuit has exciting and untapped potential to be interfaced with the discovery of small molecule modulators, of which there are not many.^18,24,73^ We acknowledge that progress to this end and beyond can be limited by this first-generation biosensor itself. As such, we will draw on the modular design, evolvability, and structural insights of NitrOFF to engineer an expanded portfolio with broad utility in illuminating nitrate across time and space dimensions.

## Data Availability Statement

The data in this study is available in the Main Text and Supporting Information. The corresponding authors can be contacted for additional requests.

## Supporting Information

Experimental methods and data (PDF). X-ray crystallography movies (MP4). Fluorescence imaging movies (MP4).

## Accession Codes

The structure factors and coordinates reported in this paper have been deposited in Protein Data Bank. Crystal Structure of NitrOFF in the bound and unbound states are deposited as 9NCU and 9NCX, respectively.

## Author Contributions

S.C.D. designed and supervised the research project with contribution from Y.J.Z. for X-ray crystallography. M.A.C. carried out the directed evolution process. M.A.C. and K.J. carried out protein purification and spectroscopy. J.D.S. carried out the X-ray crystallography. M.A.C. and S.M.P. carried out the in-cell assays. J.N.T. contributed technical expertise to the in-cell assays.

W.K. contributed technical expertise to the X-ray crystallography. K.J., W.S.Y.O., W.P., and C.M. contributed to early-stage design and engineering, purification, and spectroscopy. M.A.C., J.D.S., K.J., Y.J.Z., and S.C.D. prepared the manuscript with input from all co-authors.

## Acknowledgements

S.C.D. acknowledges support from the University of Texas at Dallas, Welch Foundation (AT-1918-20170325, AT-2060-20210327, AT-2060-20240404), National Institute of General Medical Sciences of the National Institutes of Health (R35GM128923), and National Science Foundation (2240095). J.N.T. acknowledges support from the American Chemical Society for the Irving S. Sigal Postdoctoral Fellowship. Y.J.Z. acknowledges support from the National Institute of General Medical Sciences of the National Institutes of Health (R35GM148356) and Campbell Professorship from the University of Texas at Austin. The X-ray diffraction data collection was supported by an agreement between the Advanced Photon Source, a U.S. Department of Energy (DOE) Office of Science user facility operated for the DOE Office of Science by Argonne National Laboratory under Contract No. DE-AC02-06CH11357, and the Diamond Light Source, the United Kingdom’s national synchrotron science facility, located at the Harwell Science and Innovation Campus in Oxfordshire, where the work was performed under proposal AU36008 to Y.J.Z. The findings reported in this manuscript are the sole responsibility of the authors and do not represent the views of the funding sources.

