## Supporting Information for "NitrOFF: An engineered fluorescent biosensor to illuminate nitrate transport in living cells"

### Table of Contents

Methods……………………………………………………………………………………………S3

Figures and Tables…………………………………………………………………………………S13

Summary of fluorescent protein-based biosensors for nitrate…………………………....S13

Structural and molecular dynamics characterization of NreA……………………………..S14

Plasmid design of NreA-EGFP for expression in *E. coli*…………………………………..S15

SEC and SDS-PAGE gel of purified NreA-EGFP…………………………………………..S16

Spectroscopic characterization of NreA-EGFP…………………………………………….S17

Error prone PCR and library screening results……………………………………………..S21

Plasmid design of NitrOFF for expression in *E. coli*………………………………………..S40

SEC and SDS-PAGE gel of purified NitrOFF for spectroscopic characterization....…...S41

Spectroscopic characterization of NitrOFF…………………………………………………S42

SEC and SDS-PAGE gel of purified NitrOFF for protein crystallization………………….S54

Data and statistics for crystallization of NitrOFF……………………………………………S55

RpHluorin2 cloning conditions……………………………………………………………….S56

Plasmid design of RpHluorin2 for expression in mammalian cells………………………S58

Plasmid design of NitrOFF for expression in mammalian cells…………………………...S61

Fluorescence microscopy data in HEK 293 cells…………………………………………..S62

References……………………………………………………………………………………..S74

### Methods

**General.** Unless specified, all chemicals, reagents, and supplies used in this study were acquired from Research Products International, Sigma-Aldrich, Thermo Fisher Scientific, Thomas Scientific, USA Scientific, or VWR.

**Bacterial plasmid design, construction, and preparation.** The amino acid sequence of NreA from *Staphylococcus carnosus* (NreA, UniProt ID: B9DL91) was inserted into the β-bulge region of mEGFP (FPbase ID: QKFJN) between residues N144 and N149 to generate the NreA-EGFP construct. The initial methionine residue of NreA was omitted, and the final P155 residue of NreA was mutated to a glycine residue. The gene encoding NreA-EGFP was commercially codon optimized for expression in *Escherichia coli* K12, synthesized, and cloned into the pET-21a(+) expression vector between the NdeI and NotI restriction enzyme sites with a C-terminal polyhistidine tag (Genscript, Figure S2).

From a 200 ng/µL stock solution of the commercially prepared plasmid (4 µg), one microliter of 5 ng/µL diluted plasmid was used to transform *E. cloni* 10G ELITE Competent Cells (Lucigen) via electroporation (Bio-Rad Laboratories). The transformation mixture was then plated on a Miller's Luria Broth (LB, 10 g/L sodium chloride) agar plate containing 100 µg/mL ampicillin and incubated for ~16 h at 37 °C (New Brunswick Innova 42R, Eppendorf). A single colony was used to inoculate 5 mL of LB containing 100 µg/mL ampicillin and incubated for ~16 h at 37 °C with shaking at 250 rpm. Following this, cells were harvested via centrifugation at 2,500*g* (5810 R, Eppendorf) for 15 min, and the plasmid was isolated using the QIAprep Spin Miniprep Kit (Qiagen).

**Large-scale protein expression and purification.** Briefly, the NreA-EGFP and NitrOFF1 plasmid was prepared and transformed into *E. cloni* EXPRESS BL21 (DE3) cells via electroporation as described above. Ten mL of Terrific Broth (TB) media containing 100 µg/mL ampicillin was inoculated with a single colony and incubated overnight at 30 °C for 13 h with shaking at 250 rpm. The next day, 750 µL of the overnight culture was used to inoculate 600 mL of TB media containing 100 µg/mL ampicillin and incubated at 37 °C for 4.5 h with shaking at 250 rpm until the OD600 of ~4 was reached. The culture was gradually cooled to 18 °C for 1 h with shaking prior to induction with 0.5 mM isopropyl β-d-1-thiogalactopyranoside (IPTG). The expression culture was incubated at 18 °C for 20 h with shaking at 250 rpm. The cells were harvested via centrifugation at 3,000*g* for 30 min at 4 °C. Following a wash and resuspension step with pre-chilled 20 mM Tris buffer at pH 7.5 with 200 mM sodium chloride, the cell pellet was centrifuged again and stored dry at -20 °C. Purification was carried out as previously described.[S1] Briefly, all protein samples were purified using affinity, desalt, and size exclusion chromatography to isolate the monomeric form in a chromatography refrigerator at 10 ˚C (Figures S3, S24).

For crystallization, the protein samples were purified following the same protocol with the following modifications. After elution from the affinity column, half of the fractions with an absorbance of 280 nm were pooled and loaded on a desalting column with 20 mM Tris buffer at pH 7.5 with 150 mM sodium chloride for the apo or ON state of the protein. The remaining half was loaded onto the column with 20 mM Tris buffer at pH 7.5 with 150 mM sodium nitrate for the nitrate bound or OFF state. All fractions with an absorbance at 280 were re-pooled and re-concentrated using an EMD Millipore Amicon Ultra-15 Centrifugal Filter Unit with a 30 kDa molecular weight cut-off. The concentrated fractions were loaded onto a pre-equilibrated size-exclusion column and eluted with 20 mM Tris buffer at pH 7.5 with either 150 mM sodium chloride for the ON state or 150 mM sodium nitrate for the OFF state. Fractions of both the ON and OFF states were collected and re-concentrated using an EMD Millipore Amicon Ultra-15 Centrifugal Filter Unit with a 30 kDa molecular weight cut-off (Figure S38).

The purity of each protein batch was evaluated using SDS-PAGE as previously described (Figures S3, S24, S37).[S2]

**Protein concentration determination.** The protein concentrations were determined using UV-vis spectroscopy following our previously reported studies.[S2,S3] The predicted extinction coefficient for the full-length protein was determined to be 43,320 $M^{-1}cm^{-1}$ for the parent NreA-EGFP and NitrOFF using the ProtParam program in ExPASy.[S4] The purified protein was concentrated to ~200 µM for spectroscopic measurements and to 28 and 20 mg/mL for the apo (ON) and bound (OFF) state crystallizations. Aliquots were flash frozen on dry ice and stored at -80 °C for future use.

**General spectroscopy methods.** All spectroscopy methods and analysis were adapted from our previously reported studies.[S1–S3] Measurements were acquired on a Tecan Spark plate reader at room temperature (24–26 °C). Absorbance spectra were recorded from 350 – 550 nm (2 nm step size, 3.5 nm bandwidth). For emission spectra, when excitation was provided at 400 nm (5 nm bandwidth), the emission signal was recorded from 470 – 620 nm (2 nm step size, 5 nm bandwidth, 90 gain); when excitation was provided at 480 nm (5 nm bandwidth), the emission signal was recorded from 500 – 620 nm (2 nm step size, 5 nm bandwidth, 90 gain). For each protein batch (*n* = 2), three technical replicates were carried out for each measurement described below.

**Anion titrations and apparent dissociation constants.** Frozen aliquots of purified protein samples were thawed on ice and diluted 50-fold (8 µL of 200 µM protein in 392 µL anion solutions) to a final protein concentration of ~4 µM in 25 mM sodium phosphate buffer, 1 mM sodium chloride at pH 7 with various concentrations of sodium nitrate, nitrite, and iodide. For titrations of NreA-EGFP with sodium nitrate, the buffer contained 0, 25.5, 51, 102, 153, 204, 255, 306, 357, 408, 510, 612, 714, and 816 mM sodium nitrate for a final concentration of 0, 25, 50, 100, 150, 200, 250, 300, 350, 400, 500, 600, 700, and 800 mM, respectively. For titrations of NitrOFF with sodium nitrate, the buffer contained 0, 2, 4.1, 6.1, 8.2, 10.2, 15.3, 25.5, 51, 76.5, 102, 204, 255, and 510 µM sodium nitrate for a final concentration of 0, 2, 4, 6, 8, 10, 15, 25, 50, 75, 100, 200, 250, 500 µM, respectively. For titrations of NitrOFF with sodium nitrite, the buffer contained 0, 0.01, 0.05, 0.10, 0.20, 0.31, 0.41, 0.51, 0.76, 1.02, 2.55, 5.10, and 10.2 mM sodium nitrite for a final concentration of 0, 0.01, 0.05, 0.1, 0.2, 0.3, 0.4, 0.5, 0.75, 1, 2.5 5, 7.5, 10 mM, respectively. For titrations of NitrOFF with sodium iodide, the buffer contained 0, 0.01, 0.25, 0.05, 0.076, 1.0, 1.53, 2.55, 5.10, and 10.2 mM sodium iodide for a final concentration of 0, 0.01, 0.025, 0.05, 0.075, 0.1, 0.25, 0.5, 0.75, 1, 1.5, 2.5, 5, 10 mM, respectively. Each sample was prepared at a volume of 400 µL and a 100 µL portion was added to three wells of a 96-well half-area UV-star microtiter plate (Greiner Bio-One). The samples were scanned as described in the *General spectroscopy methods* section above. The absorbance and emission spectra ($\lambda_{ex}$ = 400 nm and $\lambda_{ex}$ = 480 nm) from each protein batch are plotted as the average with standard deviation of three measurements (Figures S4, S25, S31, S32).

The following equation was used to calculate the apparent dissociation constant ($K_d$) for each anion:

$$F_{\text{obs}} = \frac{[X^-] \times (F_{\text{max}} - F_{\text{min}})}{K_d + [X^-]} + F_{\text{min}}$$

Where $F_{obs}$ is the average fluorescence intensity at 514 nm ($\lambda_{ex}$ = 480 nm) for each concentration tested, $F_{min}$ and $F_{max}$ are the lowest and highest fluorescence intensities for all the concentration tested, respectively. The $K_d$ values from two protein batches are reported as the average with propagated standard deviation (Figures S5, S26, S31, S32).

To test for ionic strength effects, frozen aliquots of purified protein samples were thawed on ice and diluted 50-fold to a final protein concentration of ~4 μM in 25 mM sodium phosphate buffer, 1 mM sodium chloride at pH 7 containing 0, 10.2, and 102 mM sodium chloride or gluconate for a final concentration of 0, 10, and 100 mM, respectively. Samples were prepared and scanned as described above (see *General spectroscopy methods*). The fold change ($F_f/F_i$) were calculated as $F_i$ being the emission intensity at 514 nm ($\lambda_{ex}$ = 480 nm) in the absence of any anions, and $F_i$ being the emission intensity at 514 nm ($\lambda_{ex}$ = 480 nm) in the presence of 10 or 100 mM anion. Data from the two protein batches are reported as the average with propagated standard deviation (Figures S6, S7, S33, S34).

**Protein extinction coefficients and quantum yields.** For extinction coefficient (ε) measurements, frozen aliquots of purified protein samples were thawed on ice and diluted 50-fold (54 μL of 200 μM protein in 2646 μL of 25 mM sodium phosphate buffers at pH 7 containing 0 and 0.51 mM sodium nitrate) to a final protein concentration of 4 μM in 25 mM sodium phosphate buffer at pH 7 with a final concentration of 1 mM sodium chloride and 0 or 0.5 mM sodium nitrate. The resulting solutions were diluted 1.25-fold into the same buffers to a final protein concentration of 3.2 μM. These working stock solutions were serially diluted six more times into the same buffers. Absorbance spectra were recorded from 250 – 550 nm (2 nm step size, 3.5 nm bandwidth).

The absorbance intensities at 280, 396 and 492 nm were plotted versus the sample dilution factors to obtain the respective slopes: $A_{280}$, $A_{396}$, and $A_{492}$. Data points that fell outside of the linear fit of the curve were not included in the analysis. The following reduction of the Beer-Lambert law was used as the optical pathlengths ($l$) and protein concentrations ($c$) are constant:

$$\Delta A = \varepsilon \times l \times \Delta c$$

$$\varepsilon_{396} = \varepsilon_{280} \times \frac{A_{396}}{A_{280}}$$

$$\varepsilon_{492} = \varepsilon_{280} \times \frac{A_{492}}{A_{280}}$$

where $\varepsilon_{280}$, $\varepsilon_{396}$, and $\varepsilon_{492}$ are the extinction coefficients at 280, 396, and 492 nm, respectively. Data from the two protein batches are reported as the average with propagated standard deviation (Figures S27, S28).

For quantum yield (Φ) determination, the same procedure as the extinction coefficient was used to prepare the protein samples. The absorbance and emission ($\lambda_{ex}$ = 480 nm) spectra were collected as described in the *General spectroscopy methods* section above. For each spectrum, the area under the curve was integrated. The ratio of the integrated areas from 500 – 700 nm ($\lambda_{ex}$ = 480 nm) and from 470 – 700 nm ($\lambda_{ex}$ = 400 nm) was used to extrapolate the integrated area from 470 – 500 nm for the emission spectra of $\lambda_{ex}$ = 480 nm. The integrated area from 470 –700 nm ($\lambda_{ex}$ = 480 nm) was plotted versus the absorbance intensities at 480 nm. This was fitted to a linear trendline.

Data outside of the linear fit of the curve was not included in the analysis. The following equation was used to calculate the quantum yields:

$$\Phi_{FP} = \Phi_{\mathrm{ref}} \times \left(\frac{F_{\mathrm{FP}}}{F_{\mathrm{ref}}}\right) \times \left(\frac{\eta_{\mathrm{FP}}}{\eta_{\mathrm{ref}}}\right)^2$$

Where $\Phi_{ref}$ is the quantum yield of fluorescein ($\Phi_{ref}$ = 0.92), $F_{FP}$ and $F_{ref}$ are the slopes of the linear plots for the protein tested and fluorescein respectively, and $\eta_{FP}$ and $\eta_{ref}$ are defined to be equivalent to the refractive index of water ($\eta$ = 1.33).[S5,S6]

The values of $F_{ref}$ for fluorescein were obtained by plotting the integrated areas of fluorescent intensities from a linear range of the serially diluted samples of fluorescein and Coumarin-153 using the above equation (Figure S29).[S5,S7] Data from the two protein batches are reported as the average with propagated standard deviation (Figure S30).

**Chromophore p*K*a.** Frozen aliquots of purified protein samples were thawed in the refrigerator and diluted 50-fold (8 μL of 200 μM protein in 396 μL of 0 or 0.51 mM sodium nitrate solution at each pH) to a final protein concentration of 4 μM in 50 mM sodium acetate buffer, 1 mM sodium chloride from pH 3 – 5.5, 50 mM MES buffer, 1 mM sodium chloride from pH 5.5 – 6.5, 50 mM MOPS buffer, 1 mM sodium chloride from pH 6.5 – 7.6, and 50 mM bicine buffer, and 1 mM sodium chloride from pH 7.6 – 9.0 in the absence and presence of 0.5 mM sodium nitrate. The absorbance and emission ($\lambda_{ex}$ = 480 nm) spectra were collected as described in the *General spectroscopy methods* section above.

The p$K_a$ values of apo and nitrate bound NitrOFF for each protein preparation were determined using the Henderson-Hasselbach equation as follows:

$$F_{\mathrm{obs}} = \frac{F_{\mathrm{min}} + \left(F_{\mathrm{max}} \times 10^{(\mathrm{pH}-\mathrm{p}K_{\mathrm{a}})}\right)}{1 + 10^{(\mathrm{pH}-\mathrm{p}K_{\mathrm{a}})}}$$

where $F_{obs}$ is the normalized fluorescence intensity observed at 514 nm in all the pH ranges tested, $F_{min}$ and $F_{max}$ are the lowest and highest fluorescence intensities observed in all the pH ranges tested, respectively.

The fitting of the Henderson-Hasselbach equation was minimized using a weighted differential evolution method, where the initial values for $F_{max}$ or $F_{min}$ were set to 1 and 0, respectively. Data from the two protein batches are reported as the average with propagated standard deviation (Figures S35, S36).

**Library cloning.** The random mutagenesis library was generated using the prepared NreA-EGFP plasmid as a starting template. Error-prone polymerase chain reaction (EP-PCR) and PCR were carried out in parallel using two sets of complementary primers (Sigma-Aldrich) based on our previous study with modifications.[S8] The forward and reverse primer sets used depended on the region that was being mutated (Table S2). The low fidelity Taq DNA polymerase (New England Biolabs) with the addition of manganese chloride was used to amplify the insert with mutations. The insert consisted of the NreA domain or the entire NreA-EGFP construct for libraries 1 – 4 and 5 – 7, respectively. Final concentrations of 50, 100, 200, 300, 400, or 500 mM manganese chloride were used to generate libraries with varying degrees of error rates. The optimal manganese chloride concentration was determined for each library through retention of function (ROF) plots. The details regarding the ROF plots are described in the section below. The high fidelity Phusion Hot Start Flex (New England Biolabs) was used to amplify the backbone without mutations. The backbone consisted of the EGFP domains plus the pET-21a(+) vector or just the pET-21a(+) vector for libraries 1 – 4 and 5 – 7, respectively. The reaction components and PCR conditions used to amplify the insert and backbone are shown in Table S3. The PCR products were incubated with 1 μL of DpnI (New England Biolabs) at 37 °C for 1 h. Following this, the PCR products were purified using 1% agarose (Gold Biotechnology) gel electrophoresis and the Zymoclean Gel DNA Recovery Kit (Zymo Research). Concentrations of the purified PCR products were determined using the NanoDropLite Spectrophotometer (Thermo Fisher Scientific). For assembly, 100 ng of total DNA (3:1 concentration of insert to backbone) was diluted to 10 μL and incubated with 10 μL of 2X HiFi Assembly Master Mix (New England Biolabs) at 50 °C for 1 h. The resulting reaction product was purified with the DNA Clean & Concentrator Kit 5 (Zymo Research) and stored at -20 °C until further use. This procedure was repeated for each round of directed evolution.

**ROF library expression and screening.** To determine the optimal manganese chloride concentration to use for each round, 1 μL of the template and library plasmids as prepared above were used to transform *E. cloni* EXPRESS BL21(DE3) Competent Cells (Lucigen) via electroporation. The transformation mixtures were plated onto LB agar plates containing 100 μg/mL ampicillin and incubated at 37 °C for 14 h. To express the colonies, autoinduction media (AIM) with 100 μg/mL ampicillin was prepared by combining 928 mL autoclaved ZY media (10 g/L bacto tryptone, 5 g/L yeast extract), 20 mL autoclaved 50x 5052 (0.5% glycerol, 0.05% glucose, 0.2% α-lactose), 50 mL sterile filtered 20x NPS (25 mM ammonium sulfate, 50 mM disodium phosphate, 50 mM monopotassium phosphate), and 1 mL sterile filtered 1 M magnesium sulfate.[S9] To a 96-well deep well plate (Griener Bio-One), 1050 μL AIM was added to each well using a 96-channel electronic pipette (MINI96, INTEGRA Biosciences). Six parent colonies and 88 random library colonies were picked into each well with the remaining two wells only containing AIM as a control for contamination. One plate was picked for each manganese chloride concentration used. The deep-well plate was sealed with an EasyApp microporous film (USA Scientific) and incubated at 37 °C for 14 h with shaking at 250 rpm and chilled at 10 °C for 10 h. Following expression, glycerol stocks of each expression plate were prepared by transferring 50 μL of expression culture from each well via an automated liquid handler to a new deep-well plate containing 50% glycerol. The glycerol stock plates were mixed via vortexing and stored at –80 °C. The cells were then harvested via centrifugation at 2,500*g* for 15 min at 4 °C and the supernatant was removed. The harvested cell pellets were stored at -20 °C.

On the day of screening, the cell pellets underwent three freeze/thaw cycles in 10 min increments prior to resuspension in 500 µL of 50 mM phosphate buffer at pH 7.4 with 2 mM magnesium chloride, 10 µg/mL deoxyribonuclease I (DNase I), and 2 mg/mL lysozyme. Plates were incubated at 37 °C for 90 min with shaking at 250 rpm prior to lysate clarification via centrifugation at 2,500*g* (Allegra X-14R, Beckman Coulter) for 30 min. An automated liquid handler (Biomek NXP liquid handler, Beckman Coulter) was used to transfer 175 µL clarified lysate to a clear 96-well microtiter plate (Caplugs). Microtiter plates were scanned on a Spark 10M plate reader (Tecan) at room temperature. Excitation was provided at 485 nm (5 nm bandwidth), and the emission intensity was recorded at 515 nm ($F_i$, 5 nm bandwidth; 30 flashes; 100, 90, 70, or 60 gain for libraries 1, 2 – 5, 6, and 7, respectively) for each well. Following this, 25 µL of 50 mM phosphate buffer at pH 7.4 with 800, 400, 8, or 0.8 mM sodium nitrate (for libraries 1, 2, 3 – 6, and 7, respectively) was added to each well to a final concentration of 100, 50, 1, or 0.1 mM sodium nitrate, respectively, and the emission intensity at 515 nm was recorded again ($F_f$). Using these measurements, the fluorescence response of each well was calculated as a percentage:

$$\Delta F/F = \left(\frac{F_f - F_i}{F_i}\right) \times 100\%$$

In each plate, the average fluorescence response (ΔF/F) of the parent with standard deviation (σ) and coefficient of variance (CV = σ / (ΔF/F)) were determined (Tables S4, S5). The ROF was determined by the percentage of library colonies that retained parent-like function. Colonies that exhibited a fluorescence response (ΔF/F) within three standard deviations of the parent average were considered parent-like and colonies with a response outside of the three standard deviations were considered dead. The manganese chloride concentrations that retained 30 – 40% function were selected for further library expression and screening (Figures S8, S10, S13, S15, S17, S19, S21).

**Library expression, screening, and validation.** Parent and library plasmids were transformed, expressed, and screened following the same protocol as for the ROF library with the following modifications. For libraries 1 and 2, approximately 1,800 library colonies were randomly picked for expression. For libraries 3 – 7, library colonies were selected based on fluorescence using a fluorescence imager (Azure 400, Azure Biosystems). Agar plates with transformed parent colonies were first excited at 472 nm and the emission intensity was captured (*f*1:16) with a 513 nm filter. The average relative fluorescence units (RFUs) of the parent colonies were determined. Agar plates with transformed library colonies were then imaged with the same settings in two different channels. The image captured in the green channel was adjusted so that the lower RFU limit was equal to the average RFUs of the parent colonies. The image captured in the grayscale channel was not adjusted so all colonies were visible in the final composite image shown. Approximately 3,000 – 4,000 colonies in each library were filtered so that only colonies displaying fluorescence greater than or equal to the parent were selected for further expression and screening (250 – 700 colonies, Figure S12). As described above for the ROF libraries, the expression cultures were preserved in glycerol, harvested, stored, and screened. Library variants exhibiting a fluorescence quenching response (ΔF/F) greater than the parent average were restreaked from the glycerol stock plates onto LB agar plates containing 100 µg/mL ampicillin and incubated at 37 °C for 14 – 16 hours for rescreening. For each variant, two colonies were picked into 5 mL of AIM and incubated at 37 °C for 14 h with shaking at 250 rpm followed by chilling at

10 °C for 10 h. Following this, the cells were harvested via centrifugation at 2,500*g* for 15 min at 4 °C and the supernatant was removed. The harvested cell pellets were stored at -20 °C. On the day of the rescreen, the cell pellets underwent three freeze/thaw cycles in 10 min increments prior to resuspension in 2.5 mL of 50 mM phosphate buffer at pH 7.4 with 2 mM magnesium chloride, 10 µg/mL deoxyribonuclease I (DNase I), and 2 mg/mL lysozyme. The pellets were incubated at 37 °C for 90 min with shaking at 250 rpm prior to lysate clarification via centrifugation at 2,500*g* (5810 R, Eppendorf) for 30 min. To a microtiter plate, 175 µL clarified lysate was added for each variant. Each well was excited at 485 nm (5 nm bandwidth) and the emission was recorded from 500 – 600 nm (5 nm step size, 5 nm bandwidth, 30 flashes). Following an initial scan to ensure consistency in the baseline fluorescence for each variant, 25 µL of 0, 0.8, 4, 8, 40, 200, 400, and/or 80 mM sodium nitrate was added to the wells for a final concentration of 0 ($F_i$), 0.1, 0.5, 1, 5, 25, 50, and 100 mM ($F_f$) sodium nitrate. The emission intensity at 515 nm was used to calculate the fluorescence quenching response ($\Delta F/F = (F_f - F_i) / F_i \times 100\%$). Data for two biological replicates with two technical replicates were collected for each variant (Figures S9, S11, S14, S16, S18, S20, S22). In parallel, the plasmids for each variant were extracted using the QIAprep Spin Miniprep Kit for Sanger sequencing (Eurofins Scientific, Tables S4, S5). The plasmid for the variant exhibiting both the greatest improvement in fluorescence quenching response (ΔF/F) and initial baseline fluorescence in lysate was targeted for further mutagenesis.

**Obtaining Diffraction Quality Crystals.** To identify crystallization conditions for the OFF and ON states, NitrOFF protein was concentrated to ~20 mg/mL and screened using sparse matrix format. The OFF state was prepared in a buffer containing 150 mM sodium nitrate, while the ON state buffer contained 150 mM sodium chloride as described above. The crystallization conditions were then optimized systematically, yielding with cubic-shaped crystals in a mother liquor containing 0.1 M sodium acetate, and 1 M ammonium sulfate at pH 5.4 for the OFF state and 1 M ammonium phosphate, 0.1 M Tris at pH 7.0 for the ON state at room temperature. NitrOFF crystals were subsequently flash-frozen in liquid nitrogen following a brief incubation in a mother liquor solution supplemented with 30% (v/v) glycerol.

**Crystal Data Collection and Analysis.** X-ray diffraction data was collected at the Diamond Light Source (Oxfordshire, UK). X-ray diffraction data for the OFF and ON states were processed to resolutions of 3.17 Å and 3.00 Å using xia2 3dii and xia2 dials, respectively.[S10–S13] Phases were determined by molecular replacement using structures for EGFP from PDB ID: 7VCM for and NreA from PDB ID: 6IZJ as search model with the Phenix software suite.[S14–S16] The initial solution resulting from molecular replacement was subjected to iterative refinement using the Coot and Phenix refine package.[S17,S18] Flexible linker regions were manually built within Coot and further refined through iterative cycles in Phenix.refine. The quality of the final structure models was evaluated using MolProbity.[S19–S21] The final statistics for data collection and structure determination are included in Table S6.

**Mammalian plasmid construction and preparation.** The gene encoding the ratiometric pHluorin2 (RpHluorin2) with a trans-Golgi localization sequence was purchased from Addgene (Gal-T-pHluorin2, Addgene # 171719).[S22,S23] To use RpHluorin2 within the cytosol and nucleus, the gene encoding RpHluorin2 without the trans-Golgi localization sequence was subcloned into the pcDNA3.1(+) vector. The high fidelity Phusion Hot Start Flex (New England Biolabs) was used to amplify both the RpHluorin2 insert and the pcDNA3.1(+) backbone following the same protocol as the library cloning described above. The reaction components and PCR conditions used to amplify the insert and backbone are shown in Tables S7 and S8. The resulting plasmid construct was transformed into *E. cloni* 10G ELITE Competent Cells (Lucigen) as described in the *Bacterial plasmid design* section above. Insertion of the RpHluorin2 gene without the localization sequence into the pcDNA3.1(+) vector was then confirmed by Sanger Sequencing (Eurofins) (Figure S38). The gene encoding NitrOFF was commercially codon optimized for mammalian expression and cloned between the BamHI and EcoRI restriction sites in the pcDNA3.1(+) vector with a Kozak sequence preceding the start codon (GenScript, Figure S41). The method for preparation of endotoxin-free plasmids for mammalian transfection was adapted from our previous study.[S8]

**HEK293 cell culture and transfection** Human embryonic kidney (HEK) 293 cells were purchased from the American Type Culture Collection (ATCC, CRL-1573). Cells were grown in Dulbecco's Modified Eagle Medium (DMEM) (Gibco) supplemented with 10 % fetal bovine serum (FBS) and 1% Penicillin-Streptomycin (100 μg/mL) at 37 °C, 5% carbon dioxide in a T25 flask (Corning). Cells were regularly split at 70 – 80% confluency. Briefly, the cells were washed twice with 5 mL 1x Phosphate Buffered Saline (PBS, Gibco) and incubated in 3 mL trypsin-EDTA (0.05%) (Gibco) for 5 min at room temperature. To quench the trypsin reaction, 6 mL DMEM was added, and the cells were harvested via centrifugation at 200*g* (5702, Eppendorf) for 5 min. The cell pellet was resuspended in 2 mL DMEM. For cell counting via hemacytometer, 2 μL resuspended cells were added to 8 μL DMEM and 10 μL 0.4% trypan blue (Sigma-Aldrich).

For fluorescence imaging assays, 5 – 7 × $10^5$ cells were plated onto 35 mm dishes with a 14 mm micro-well and #1.5 glass-like polymer coverslip (Cellvis) and incubated at 37 °C, 5% carbon dioxide. The next day, a transfection mixture was prepared by combining 1.5 μg of plasmid with 2 μL lipofectamine, 3 μL P3000 reagent, and 250 μL OptiMEM Reduced Serum (Gibco). The mixture was lightly vortexed and incubated at room temperature for 30 min before being added to the dish in a dropwise manner. The cells were incubated at 37 °C, 5% carbon dioxide for two days prior to imaging.

**Fluorescence imaging of transfected HEK293 cells.** Fluorescence imaging methods were adapted from our previously published studies.[S2,S3,S8] On the day of the assay, the media was aspirated from the dish and the cells were washed with 2 mL of a prewarmed modified PBS buffer at pH 7.4 (8.1 mM disodium phosphate, 1.5 mM monopotassium phosphate, 2.7 mM potassium chloride, 0.7 mM calcium chloride, 1.1 mM magnesium chloride, and 10 mM glucose) containing 137 mM sodium chloride. The osmolality of all imaging buffers was measured and adjusted to ~280 mOsm (VAPRO, ELITechGroup) and the ionic strength was kept consistent. The cells were incubated in 2 mL buffer for 30 min in a stage top incubator with an automated perfusion system (Tokai Hit) maintained at 37 °C. An inverted microscope (IX83, Olympus) equipped with a light engine (Spectra X, Lumencor) and a 20X air objective with a numerical aperture of 0.7 was used to collect differential interference contrast (DIC) and fluorescence images. Camera resolution was set to 512 x 512 pixels with 2 x 2 binning. For both RpHluorin2 and NitrOFF, the EGFP/FITC/Cy2 excitation filter centered at 470 nm (40 nm bandwidth, Chroma) was used and exposure was provided at 25% LED power for 45 ms for RpHluorin2 and 250 – 350 ms for NitrOFF. Additionally, for RpHluorin2, the ET380x excitation filter centered at 380 nm (11 nm bandwidth, Chroma) was used and exposure was provided at 25% LED power for 30 ms. The EGFP emission filter centered at 525 nm (50 nm bandwidth, Chroma) was used for all excitations. The coordinates for four fields per dish were recorded with Z-drift compensation (ZDC).

For assays with RpHluorin2, images were recorded every min for 10 min to establish the baseline signal in the modified PBS buffer containing 137 mM sodium chloride. Following this, the imaging buffer was exchanged to the modified PBS buffer containing 0.5 mM sodium nitrate with 136.5 mM sodium chloride at a flow rate of 4 mL/min for 2 min while image acquisition continued every min for 20 min. The imaging buffer was re-exchanged with the modified PBS buffer containing 137 mM sodium chloride at a flow rate of 4 mL/min for 2 min and was incubated for 20 min. At the end of the RpHluorin2 exchange assay, the imaging buffer in the dish was manually exchanged on the stage with a pH clamping buffer at pH 6 consisting of the modified PBS buffer supplemented with 120 mM potassium chloride, 17 mM sodium chloride, 5 μM nigericin (Sigma-Aldrich), and 5 μM valinomycin (Sigma-Aldrich).[S22] Aliquots of nigericin and valinomycin were prepared at a stock concentration of 10 mM in dimethyl sulfoxide (DMSO) and diluted directly into the modified PBS buffers at 1:2,000 each. Following a 15 min incubation period, images of the same fields were recorded every min for 5 min (Figure S39, Movies S3, S4). This assay was repeated with the modified PBS buffers containing 0.25 mM sodium nitrate with 136.5 mM sodium chloride (Figure S40, Movies S5, S6). For one replicate, the imaging buffer was not re-exchanged with the modified PBS buffer containing 137 mM sodium chloride prior to the clamping phase (Figure S40, Movie S7).

For assays with NitrOFF, images were recorded every min for 10 min to establish the baseline signal in the modified PBS buffer containing 137 mM sodium chloride. Following this, the imaging buffer was exchanged at a flow rate of 4 mL/min for 2 min to the modified PBS buffer containing either 137 mM sodium chloride or 0.25 mM sodium nitrate with 136.75 mM sodium chloride. Image acquisition continued every min for 20 min. Exchange assays were also performed with 137 mM sodium gluconate replacing the sodium chloride in the modified PBS buffer. Images were recorded every min for 10 min to establish the baseline signal and the imaging buffer was then exchanged to modified PBS buffers containing 137 mM sodium chloride, 0.25 mM sodium chloride with 136.75 mM sodium gluconate and 0.2% DMSO, 0.25 mM sodium nitrate with 136.75 mM sodium gluconate and 0.2% DMSO, 0.25 mM sodium nitrate with 136.75 mM sodium gluconate and 0.1 mM 5-nitro-2-(3-phenylpropylamino)benzoic acid (NPPB, Tocris Bioscience), or 0.25 mM sodium nitrate with 136.75 mM sodium gluconate and 0.2 mM NPPB (Figures S42, S43, S45 – S49, Movies S8 – S13, S17 – S31). Aliquots of NPPB were prepared at a stock concentration of 100 mM in DMSO and diluted directly into the modified PBS buffers at 1:1,000 or 1:500 for final concentrations of 0.1 and 0.2 mM NPPB, respectively. As described above, the intracellular pH was monitored with RpHluorin2 in parallel assays where the imaging buffer was not re-exchanged with the modified PBS buffer containing 137 mM sodium chloride prior to the clamping phase (Figures S44, S50 – S53, Movies S14 – S16, S32 – S43).

**Fluorescence imaging analysis.** The imaging data was analyzed with the Fiji is Just ImageJ (Fiji v2.0) software as described in our previous studies.[S2,S3,S8] The fluorescence time lapse images were concatenated into a single stack and aligned using the StackReg Translation function. The background was then subtracted (size 75), and a mask was created from the maximum intensity Z-projection for the 480 nm excitation channel for NitrOFF and the 380 nm excitation channel for RpHluorin2 using the auto default threshold. The mask was applied to each fluorescence stack and ROIs were selected using the Fiji Analyze Particles function. ROIs with saturated intensity, containing debris, moving out of bounds, or undergoing mitosis were manually excluded. The median fluorescence intensity for each ROI at each time point was measured using the Multi Measure function in Fiji. For NitrOFF expressing cells, the median fluorescence intensity of each ROI (F) was normalized with respect to their corresponding initial median fluorescence intensity ($F_i$). For RpHluorin2 expressing cells, the ratio of the median fluorescence intensity ($F_{Ex470}/F_{Ex380}$) for each ROI (F) was normalized to the corresponding initial median fluorescence intensity ratio ($F_i$).

Outlier data points were identified using the Interquartile Range (IQR) method and omitted from analysis as previously described.[S8] Briefly, we determined the IQR by subtracting the 25$^{th}$ (Q1) from the 75$^{th}$ (Q3) percentiles. Data points that fell outside of the upper and lower limits, as determined by the equations below, were identified as outliers:

$$\text{Upper limit} = \text{Q3} + (\text{IQR} \times 1.5)$$

$$\text{Lower limit} = \text{Q1} - (\text{IQR} \times 1.5)$$

For both NitrOFF and RpHluorin2, the average of median emission response ($F/F_i$) with standard deviation from three biological replicates with four fields each is reported (Figures S39, S40, S42 – S53, Movies S3 – S43). Note: the average of the median emission response ($F/F_i$) that is reported in the Main Text is the average median emission response at the final time point of the exchange (F, $t$ = 29 min) with respect to the average median emission response at the final time point of the baseline ($F_i$, $t$ = 10 min).

### Figures and Tables

**Table S1.** Design and properties of currently available fluorescent protein-based biosensors for nitrate.

| Biosensor | Output | Fluorescent Protein | Nitrate Binding Domain | $K_d$ | Application | Reference |
|---|---|---|---|---|---|---|
| **YFP-H148Q/I152L** | Intensiometric | YFP[a] | [b] | 10 mM | 3T3 fibroblasts | S24 |
| **sCiNiS** | Intensiometric | mCitrine | NLP7 from *Arabidopsis thaliana* | 52 μM | *A. thaliana* | S25 |
| **ClopHensor** | Ratiometric | $E^2$GFP, DsRed | [b] | 5 mM | *A. thaliana* | S26 |
| **NiTrac1** | FRET[a] | mCerulean, Aphrodite | CHL1 from *A. thaliana* | 75.1 μM | *Xenopus* oocytes | S27 |
| **sNOOOpy** | FRET | CFP, Venus | NasS/NasT from *Bradyhizobium japonicum* | 39.5 μM | HeLa | S28 |
| **FLIP-NT** | FRET | CFP, YFP | NrtA from *Synechocystis sp. 6803* | 5 μM | *Escherichia coli*, *Saccharomyces cerevisiae* | S29 |
| **NiMet3.0** | FRET | edeCFP, edAFP[a] | NasR from *Klebsiella oxytoca* | 90 μM | *A. thaliana* | S30 |

[a]Abbreviations: CFP, cyan fluorescent protein; edAFP, enhanced dimerization variant of Aphrodite fluorescent protein; edeCFP, enhanced dimerization variant of enhance cyan fluorescent protein; YFP, yellow fluorescent protein. [b]For the YFP-H148Q/I152L and ClopHensor biosensors, the nitrate binding pocket is intrinsic to the fluorescent protein near the chromophore.

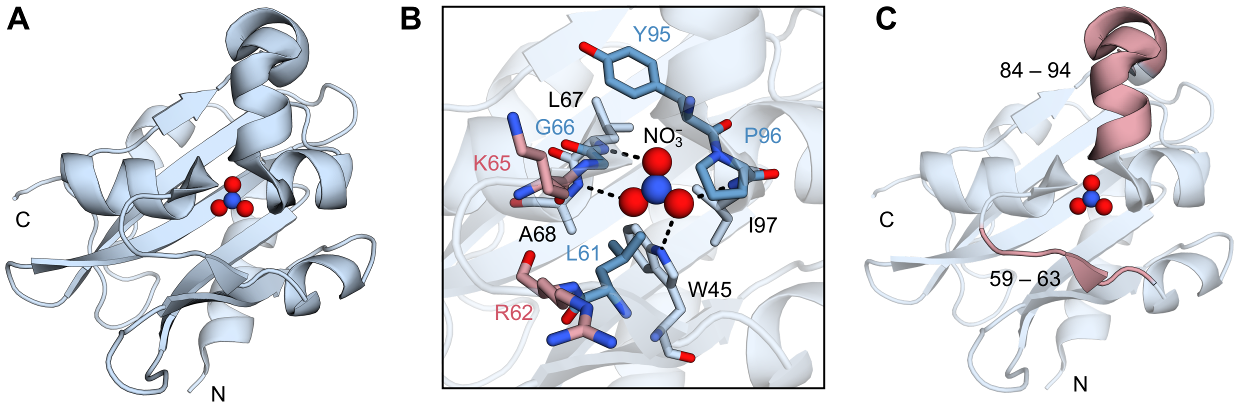


**Figure S1.** Structural and molecular dynamics (MD) characterization of NreA. (A) X-ray crystal structure of nitrate-bound NreA from *Staphylococcus carnosus* (PDB ID: 4IUK). The N and C-termini of NreA are denoted as N and C.[S31] (B) Residues that coordinate with the nitrate anion in the oxyanion hole of NreA are shown as light blue sticks. Hydrophobic residues that shield the nitrate anion from bulk water as shown as blue sticks. Residues that were revealed through MD simulations to act as anchors to draw anions into the oxyanion hole are shown as pink sticks.[S32] All residues are labeled with their amino acid identity and position according to the sequence of NreA (UniProt ID: B9DL91). (C) Conformationally flexible regions of NreA when bound to anions as revealed by root mean square fluctuations (RMSF) analysis are numbered and shown in pink.[S32]

CAT ATG GTT AGC AAG GGC GAG GAG CTG TTT ACC GGC GTG GTT CCG ATT CTG GTT GAG CTG
His Met Val Ser Lys Gly Glu Glu Leu Phe Thr Gly Val Val Pro Ile Leu Val Glu Leu 18

GAT GGT GAT GTT AAT GGT CAT AAG TTT AGC GTT AGC GGC GAG GGC GAA GGT GAC GCG ACC
Asp Gly Asp Val Asn Gly His Lys Phe Ser Val Ser Gly Glu Gly Glu Gly Asp Ala Thr 38

TAC GGC AAG CTG ACC CTG AAA TTC ATC TGC ACC ACC GGT AAA CTG CCG GTG CCG TGG CCG
Tyr Gly Lys Leu Thr Leu Lys Phe Ile Cys Thr Thr Gly Lys Leu Pro Val Pro Trp Pro 58

ACC CTG GTT ACC ACC CTG ACC TAC GGC GTG CAG TGC TTT AGC CGT TAT CCG GAC CAC ATG
Thr Leu Val Thr Thr Leu Thr Tyr Gly Val Gln Cys Phe Ser Arg Tyr Pro Asp His Met 60

AAG CAA CAC GAT TTC TTT AAA AGC GCG ATG CCG GAG GGC TAC ATC CAG GAA CGT ACC ATT
Lys Gln His Asp Phe Phe Lys Ser Ala Met Pro Glu Gly Tyr Ile Gln Glu Arg Thr Ile 98

TTC TTT AAG GAC GAT GGT AAC TAC AAG ACC CGT GCG GAA GTG AAG TTC GAA GGC GAT ACC
Phe Phe Lys Asp Asp Gly Asn Tyr Lys Thr Arg Ala Glu Val Lys Phe Glu Gly Asp Thr 118

CTG GTT AAC CGT ATC GAG CTG AAG GGT ATT GAC TTT AAA GAA GAT GGC AAC ATC CTG GGT
Leu Val Asn Arg Ile Glu Leu Lys Gly Ile Asp Phe Lys Glu Asp Gly Asn Ile Leu Gly 138

CAC AAG CTT GAG TAC AAC CTG AAC AGC GTG ATC GCG AGC GAC TAC TTC GAT TAT CAG GAC
His Lys Leu Glu Tyr Asn Leu Asn Ser Val Ile Ala Ser Asp Tyr Phe Asp Tyr Gln Asp 158

GCG CTG GAT GAG ATC CGT GAG ACC GAA AAG TTC GAC TTT GCG GCG ATT GCG CTG CCG GAA
Ala Leu Asp Glu Ile Arg Glu Thr Glu Lys Phe Asp Phe Ala Ala Ile Ala Leu Pro Glu 178

GAT GGT CTG CAC AGC GCG GTT ATT AAG TGG AAA TAC GCG AGC GGC AAC ATC AAC TAC CGT
Asp Gly Leu His Ser Ala Val Ile Lys Trp Lys Tyr Ala Ser Gly Asn Ile Asn Tyr Arg 198

TAT CGT ATG ATT GTG CTG CGT CCG GGC AAG GGT CTG GCG GGT CTG GTT ATC CGT ACC GGC
Tyr Arg Met Ile Val Leu Arg Pro Gly Lys Gly Leu Ala Gly Leu Val Ile Arg Thr Gly 218

AGC CGT AAA ATT GTG GAG GAC GTT GAT GCG GAA CTG AGC CAG AAC GAC AAG CTG GGT TAT
Ser Arg Lys Ile Val Glu Asp Val Asp Ala Glu Leu Ser Gln Asn Asp Lys Leu Gly Tyr 238

CCG ATT GTG CTG AGC GAG GCG CTG ACC GCG ATG GTT GCG ATT CCG CTG TGG AAA AAC AAC
Pro Ile Val Leu Ser Glu Ala Leu Thr Ala Met Val Ala Ile Pro Leu Trp Lys Asn Asn 258

CGT GTG TAT GGT GCG CTG CTG CTG GGT CAA CGT GAA GGT CGT CCG CTG CCG GAA GGT AGC
Arg Val Tyr Gly Ala Leu Leu Leu Gly Gln Arg Glu Gly Arg Pro Leu Pro Glu Gly Ser 278

ACC ACC TTC CGT ATC AAC CAG CGT CTG GGC AGC TTT ACC GAT GAA ATT AAC AAA CAA GGC
Thr Thr Phe Arg Ile Asn Gln Arg Leu Gly Ser Phe Thr Asp Glu Ile Asn Lys Gln Gly 298

AAC GTG TAC ATC AAG GCG GAC AAG CAG AAA AAC GGT ATT AAG GCG AAC TTC AAA ATC CGT
Asn Val Tyr Ile Lys Ala Asp Lys Gln Lys Asn Gly Ile Lys Ala Asn Phe Lys Ile Arg 318

CAC AAC ATT GAA GAT GGT GGC GTT CAA CTG GCG TAC CAC TAT CAG CAA AAC ACC CCG ATT
His Asn Ile Glu Asp Gly Gly Val Gln Leu Ala Tyr His Tyr Gln Gln Asn Thr Pro Ile 338

GGT GAT GGT CCG GTG CTG CTG CCG GAT AAC CAC TAT CTG AGC GTT CAA AGC AAG CTG AGC
Gly Asp Gly Pro Val Leu Leu Pro Asp Asn His Tyr Leu Ser Val Gln Ser Lys Leu Ser 358

AAA GAC CCG AAC GAG AAG CGT GAT CAC ATG GTG CTG CTG GAA TTT GTT ACC GCG GCG GGC
Lys Asp Pro Asn Glu Lys Arg Asp His Met Val Leu Leu Glu Phe Val Thr Ala Ala Gly 378

ATT ACC CTG GGC ATG GAC GAA CTG TAT AAG GCG GCC GCA CTC GAG CAC CAC CAC CAC CAC
Ile Thr Leu Gly Met Asp Glu Leu Tyr Lys Ala Ala Ala Leu Glu His His His His His 398

CAC TGA
His End 400

**Figure S2.** Nucleotide (top row) and amino acid (bottom row) sequence of the NreA-EGFP construct with mEGFP (FPbase ID: QKFJN) shown in green font and NreA from *Staphylococcus carnosus* (Uniprot ID: B9DL91) shown in black font. The NdeI and NotI restriction enzyme sites from the pET-21a(+) vector (gray), polyhistidine tag (blue), and stop codon (pink) are shown.

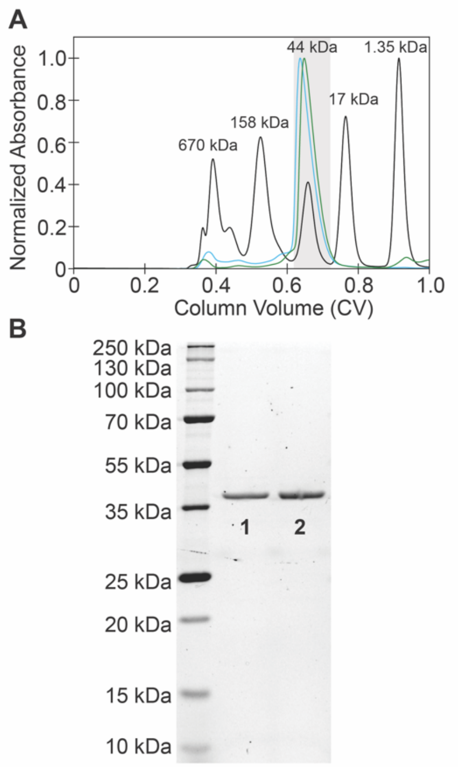


**Figure S3.** (A) Size exclusion chromatogram (SEC, normalized absorbance at 280 nm) for the first (blue) and second (green) biological replicate of the NreA-EGFP. The gel filtration standard is shown in black and made up of the following: thyroglobulin (670 kDa), $\gamma$-globulin (158 kDa), ovalbumin (44 kDa), myoglobulin (17 kDa), and vitamin B12 (1.35 kDa). All SECs were collected in 20 mM Tris buffer at pH 7.5 with 150 mM sodium chloride. For each protein run, the fraction collected for further analysis is shown in the gray area. (B) Representative SDS-PAGE of the protein batches tested in lanes 1 and 2. The theoretical molecular weight of NreA-EGFP is ~44.9 kDa.

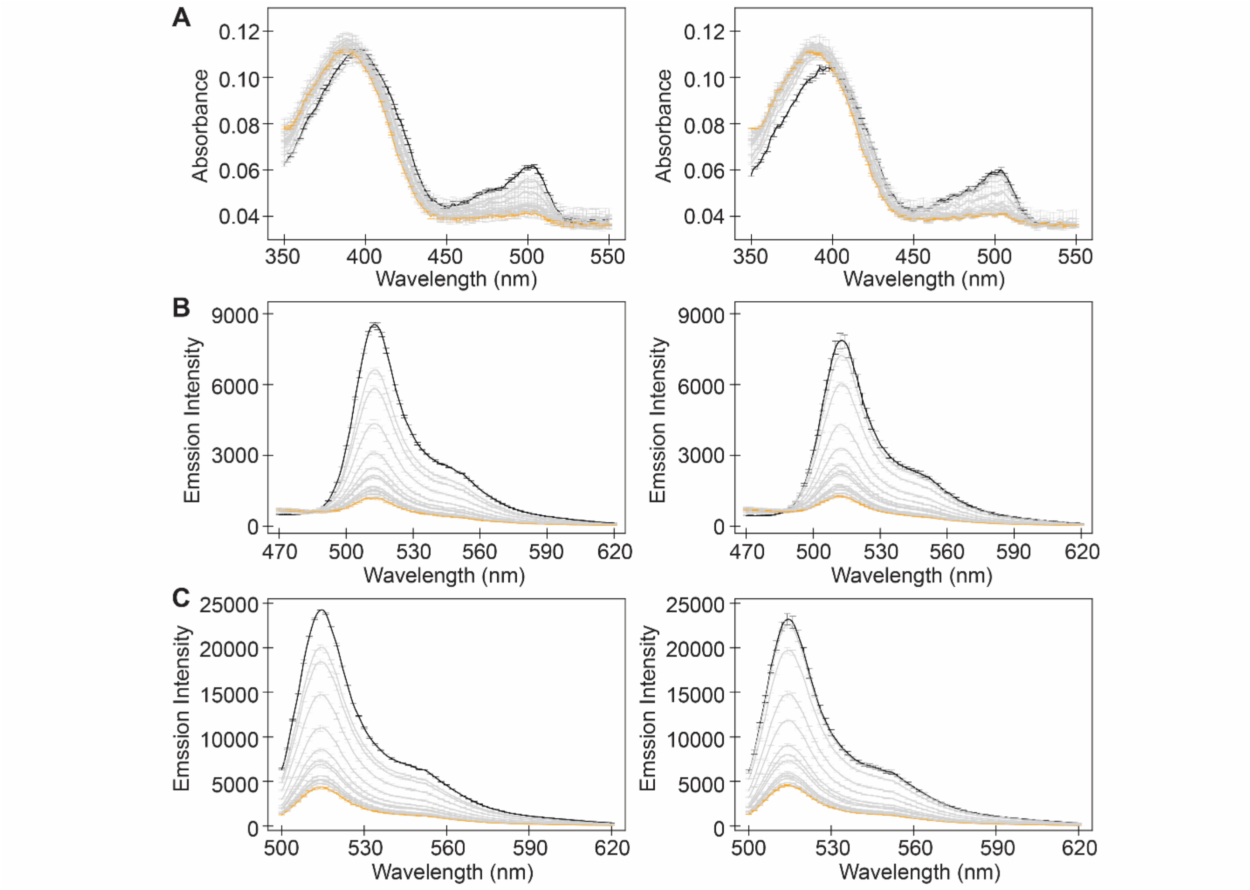


**Figure S4.** Spectroscopic characterization of NreA-EGFP with nitrate. Absorbance and emission spectra of ~4 µM NreA-EGFP excited at (A) 400 nm and (B) 480 nm in the presence of 0 mM (black), 25, 50, 100, 150, 200, 250, 300, 350, 400, 500, 600, 700 (gray) or 800 mM (orange) sodium nitrate in 25 mM sodium phosphate buffer at pH 7 with 1 mM sodium chloride. The average of three technical replicates with standard deviation is shown for one of two protein batches in each column.

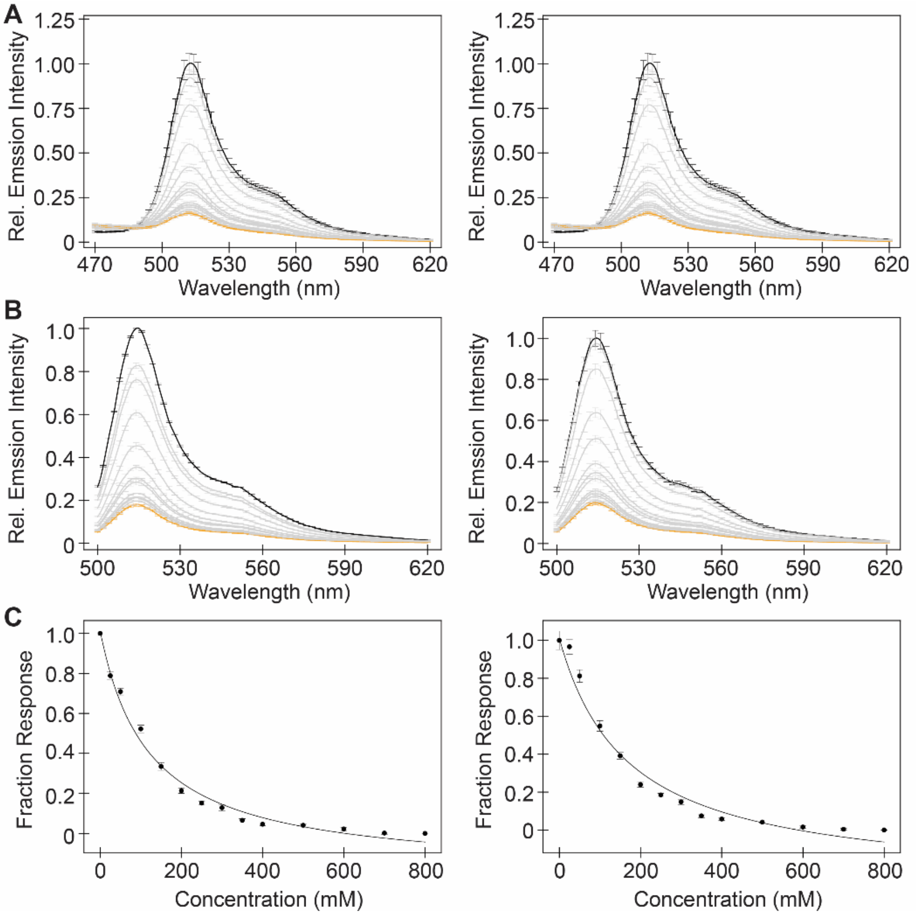


**Figure S5.** Spectroscopic characterization of NreA-EGFP with nitrate. Emission spectra of ~4 µM NreA-EGFP excited at (A) 400 nm and (B) 480 nm in the presence of 0 mM (black), 25, 50, 100, 150, 200, 250, 300, 350, 400, 500, 600, 700 (gray) or 800 mM (orange) sodium nitrate in 25 mM sodium phosphate buffer at pH 7 with 1 mM sodium chloride. (C) The emission response at 514 nm from (B) was fitted to determine the apparent dissociation constant ($K_d$) of 120.5 ± 12.1 mM and 168.4 ± 26.5 mM. The average of three technical replicates with standard deviation is shown for one of two protein batches in each column.

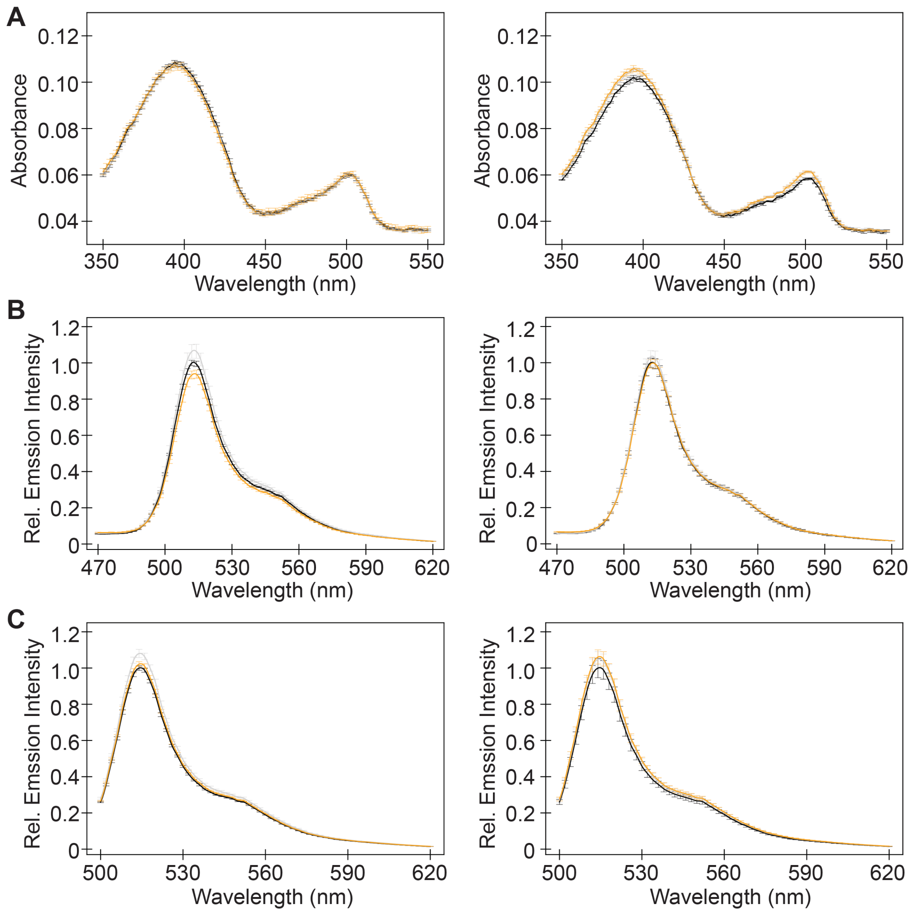


**Figure S6.** Spectroscopic characterization of NreA-EGFP with chloride. (A) Absorbance and emission spectra of ~4 μM NreA-EGFP excited at (B) 400 nm and (C) 480 nm in the presence of 0 (black), 10 (gray), and 100 mM (orange) sodium chloride in 25 mM sodium phosphate buffer at pH 7 with 1 mM sodium chloride. The average of three technical replicates with standard deviation is shown for one of two protein batches in each column.

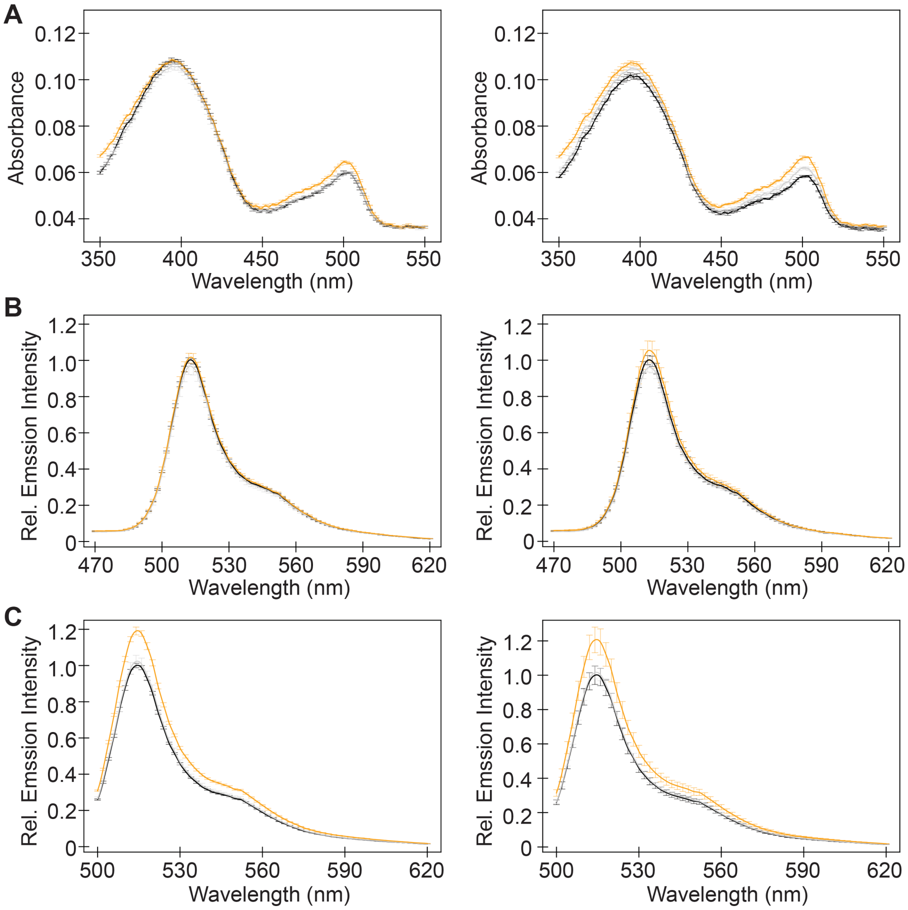


**Figure S7.** Spectroscopic characterization of NreA-EGFP with gluconate. (A) Absorbance and emission spectra of ~4 μM NreA-EGFP excited at (B) 400 nm and (C) 480 nm in the presence of 0 (black), 10 (gray), and 100 mM (orange) sodium gluconate in 25 mM sodium phosphate buffer at pH 7 with 1 mM sodium chloride. The average of three technical replicates with standard deviation is shown for one of two protein batches in each column.

**Table S2.** Primers used to generate the error prone libraries. Mutagenesis was carried out over the NreA domain only (libraries 1–4, black font) or over the entire NreA-EGFP construct (libraries 5–7, red font).

| Description | Primer Sequence (5' to 3') |
|---|---|
| NreA domain insert forward | CCGTATCGAGCTGAAGGGTATTGACTTTAAAGAA |
| NreA domain insert reverse | GCTTGTCCGCCTTGATGTACACGTT |
| NreA domain backbone forward | AACGTGTACATCAAGGCGGACAAGC |
| NreA domain backbone reverse | TTCTTTAAAGTCAATACCCTTCAGCTCGATACGG |
| NreA-EGFP insert forward | AAATAATTTTGTTTAACTTTAAGAAGGAGATATACATATG |
| NreA-EGFP insert reverse | GCCGGATCTCAGTGGTGGTGGTGGTGGTGCTCGAG |
| NreA-EGFP backbone forward | CTCGAGCACCACCACCACCACCACTGAGATCCGGC |
| NreA-EGFP backbone reverse | CATATGTATATCTCCTTCTTAAAGTTAAACAAAATTATTT |

**Table S3.** Polymerase chain reaction conditions to generate the error prone libraries. Modifications for libraries constructed using the NreA-EGFP primers are shown in red font.

| Insert Reaction Conditions | | | | | | |
|---|---|---|---|---|---|---|
| **Component \| Final [$MnCl_2$]** | **50 µM** | **100 µM** | **200 µM** | **300 µM** | **400 µM** | **500 µM** |
| DreamTaq Green 2X Master Mix | 25 µL | 25 µL | 25 µL | 25 µL | 25 µL | 25 µL |
| 10 ng/µL DNA Template | 2 µL | 2 µL | 2 µL | 2 µL | 2 µL | 2 µL |
| 10 µM Insert Forward Primer | 1 µL | 1 µL | 1 µL | 1 µL | 1 µL | 1 µL |
| 10 µM Insert Reverse Primer | 1 µL | 1 µL | 1 µL | 1 µL | 1 µL | 1 µL |
| 2 mM Manganese chloride | 1.25 µL | 2.5 µL | 5 µL | 7.5 µL | 10 µL | 12.5 µL |
| Autoclaved Water | 19.75 µL | 18.5 µL | 16 µL | 13.5 µL | 11 µL | 8.5 µL |
| **Total** | 50 µL | 50 µL | 50 µL | 50 µL | 50 µL | 50 µL |

| Insert Thermocycler Settings | | | |
|---|---|---|---|
| **Step** | **Temperature** | **Time** | **Number of Cycles** |
| Template Denaturation | 95 °C | 1 min (3 min) | 1 |
| | 95 °C | 30 s | 30 |
| Annealing | 55 °C | 30 s | |
| Extension | 68 °C (72 °C) | 30 s (75 s) | |
| Final Extension | 68 °C (72 °C) | 5 min (10 min) | 1 |
| Storage | 10 °C | ∞ | 1 |

| Backbone Reaction Conditions | |
|---|---|
| **Component** | **Volume** |
| Phusion 2X Master Mix | 25 µL |
| 10 ng/µL DNA Template | 1 µL |
| 10 µM Backbone Forward Primer | 1 µL |
| 10 µM Backbone Reverse Primer | 1 µL |
| Autoclaved Water | 22 µL |
| **Total** | 50 µL |

| Backbone Thermocycler Settings | | | |
|---|---|---|---|
| **Step** | **Temperature** | **Time** | **Number of Cycles** |
| Template Denaturation | 98 °C | 30 s | 1 |
| | 98 °C | 30 s | 30 |
| Annealing | 55 °C | 30 s | |
| Extension | 72 °C | 4 min (3.5 min) | |
| Final Extension | 72 °C | 10 min | 1 |
| Storage | 10 °C | ∞ | 1 |

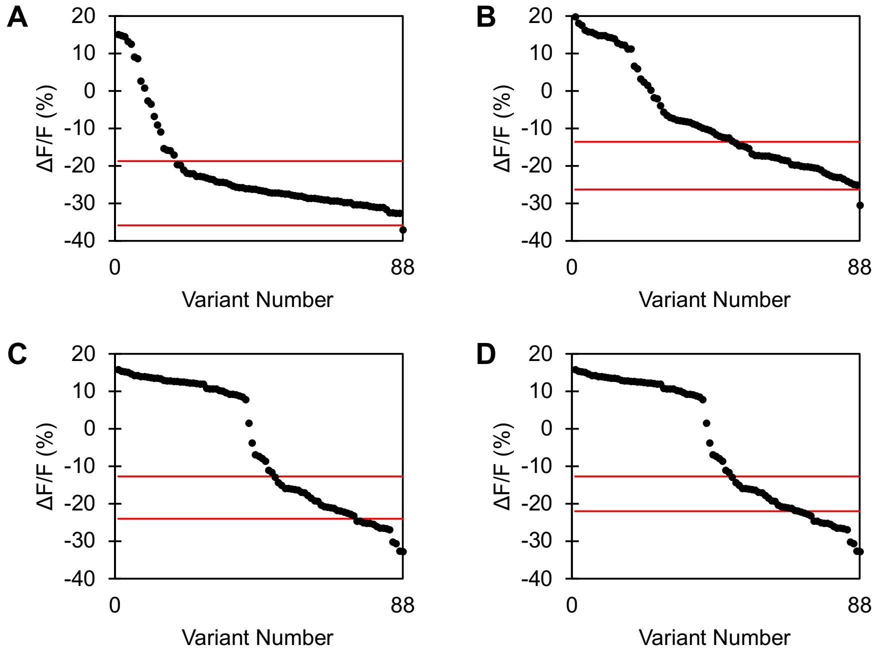


**Figure S8.** Retention of function (ROF) plots from the first directed evolution library generated by using (A) 200, (B) 300, (C) 400, and (D) 500 μM manganese chloride during the error prone polymerase chain reaction (EP-PCR). Eighty-eight variants per manganese chloride concentration were screened in the absence ($F_i$) and presence ($F_f$) of 100 mM sodium nitrate in 50 mM phosphate buffer at pH 7.4. Variants that retained the function of the parent had a fluorescence response ($\Delta F/F = (F_f - F_i) / F_i * 100\%$) within three standard deviations of the average fluorescence response for six biological replicates of the NreA-EGFP parent as indicated by the red lines. Based on these ROF plots, the libraries generated by using 200 and 300 μM manganese chloride were combined and referred to as *Library 1* for further expression and screening.

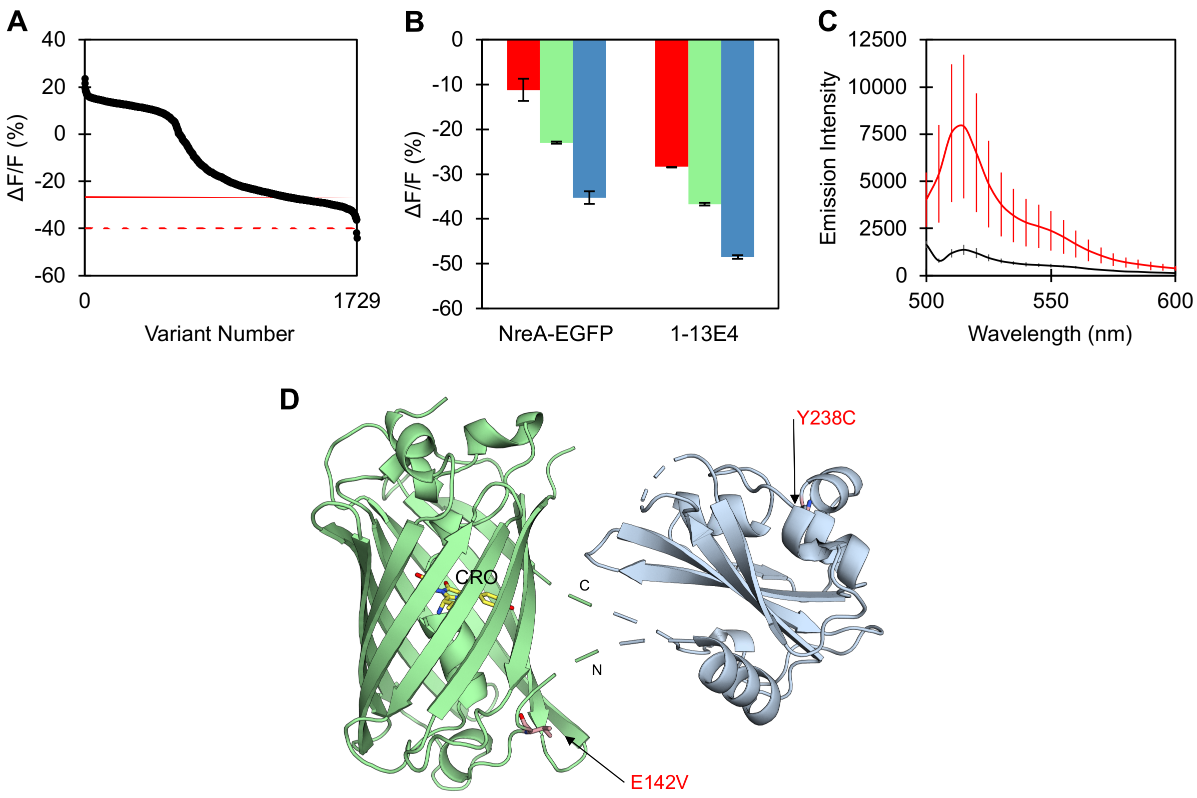


**Figure S9.** Screening of the first directed evolution library. (A) Summary plot of the first NreA domain mutagenesis library (*Library 1*) for the 1,760 variants (black circles) screened in the absence ($F_i$) and presence ($F_f$) of 100 mM sodium nitrate. The average fluorescence response ($\Delta F/F = (F_f - F_i) / F_i * 100\%$) of 100 biological replicates of the NreA-EGFP parent ($\Delta F/F = -26.7\% \pm 4.4\%$) is shown by the blue dashed line. Three standard deviations from the parent average ($\Delta F/F = -39.9\%$) was used as a threshold to identify improved variants. The variant that was selected for rescreening and used as the template for the next EP-PCR library is indicated by the red circle. (B) Summary of the rescreen of the parent and top variant. The average fluorescence response with standard deviation to 25 (red), 50 (green), and 100 (blue) mM sodium nitrate are shown for two biological replicates with two technical replicates each. (C) The average baseline emission spectra with standard deviation of the NreA-EGFP parent (black) and top performing variant (red). Excitation was provided at 485 nm. All screening was done in *E. coli* lysate with 50 mM sodium phosphate buffer at pH 7.4. (D) AlphaFold model of the NreA-EGFP construct showing the position of all mutations (shown as sticks) in the 1-13E4 variant. The positions of the mutations are highlighted as pink sticks with labels and the N- and C-terminal linker regions of the NreA domain are labeled as N and C, respectively. The EGFP chromophore (CRO) is shown as yellow sticks.

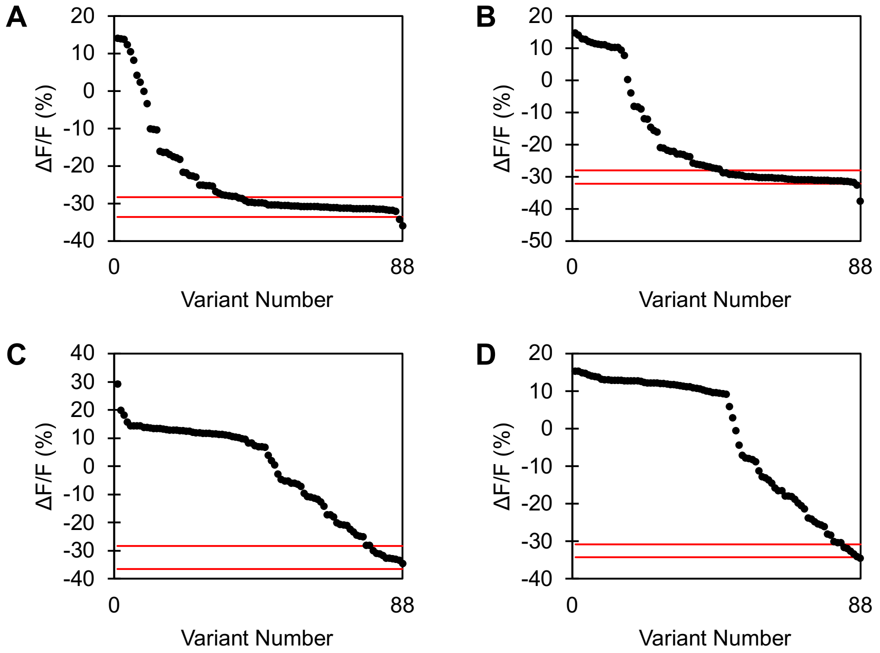


**Figure S10.** ROF plots from the second directed evolution library generated by using (A) 200, (B) 300, (C) 400, and (D) 500 µM manganese chloride during EP-PCR. Eighty-eight variants per manganese chloride concentration were screened in the absence ($F_i$) and presence ($F_f$) of 50 mM sodium nitrate in 50 mM phosphate buffer at pH 7.4. Variants that retained the function of the parent had a fluorescence response ($\Delta F/F = (F_f - F_i) / F_i * 100\%$) within three standard deviations of the average fluorescence response for six biological replicates of the 1-13E4 parent as indicated by the red lines. Based on these ROF plots, the libraries generated by using 200 and 300 µM manganese chloride were combined and referred to as *Library 2* for further expression and screening.

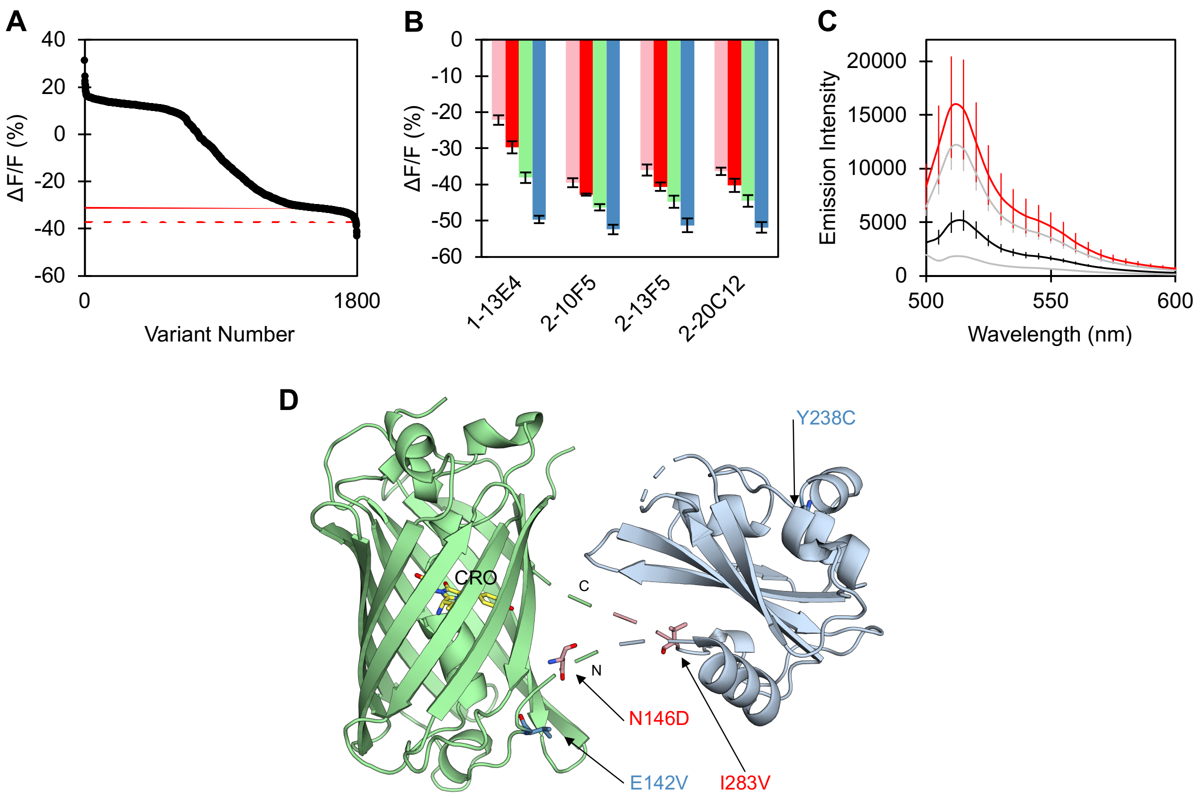


**Figure S11.** Screening of the second directed evolution library. (A) Summary plot of the second NreA domain mutagenesis library (*Library 2*) for the 1,760 variants (black circles) screened in the absence ($F_i$) and presence ($F_f$) of 50 mM sodium nitrate. The average fluorescence response ($\Delta F/F = (F_f - F_i) / F_i * 100\%$) of 120 biological replicates of the 1-13E4 parent ($\Delta F/F = -31.1\% \pm 2.1\%$) is shown by the blue dashed line. Three standard deviations from the parent average ($\Delta F/F = -37.4\%$) was used as a threshold to identify improved variants. The variant that was selected for rescreening and used as the template for the next EP-PCR library is indicated by the red circle. (B) Summary of the rescreen of the parent and top variants. The average fluorescence response with standard deviation to 5 (pink), 25 (red), 50 (green), and 100 (blue) mM sodium nitrate are shown for two biological replicates with two technical replicates each. (C) The average baseline emission spectra with standard deviation of the 1-13E4 parent (black) and top performing variant (red). The remaining top variants from (B) are shown in gray. Excitation was provided at 485 nm. All screening was done in *E. coli* lysate with 50 mM sodium phosphate buffer at pH 7.4. (D) AlphaFold model of the NreA-EGFP construct showing the position of all mutations (shown as sticks) in the 2-20C12 variant. The positions of the mutations are highlighted as pink sticks for new mutations and blue sticks for previous mutations with labels and the N- and C-terminal linker regions of the NreA domain are labeled as N and C, respectively. The EGFP chromophore (CRO) is shown as yellow sticks.

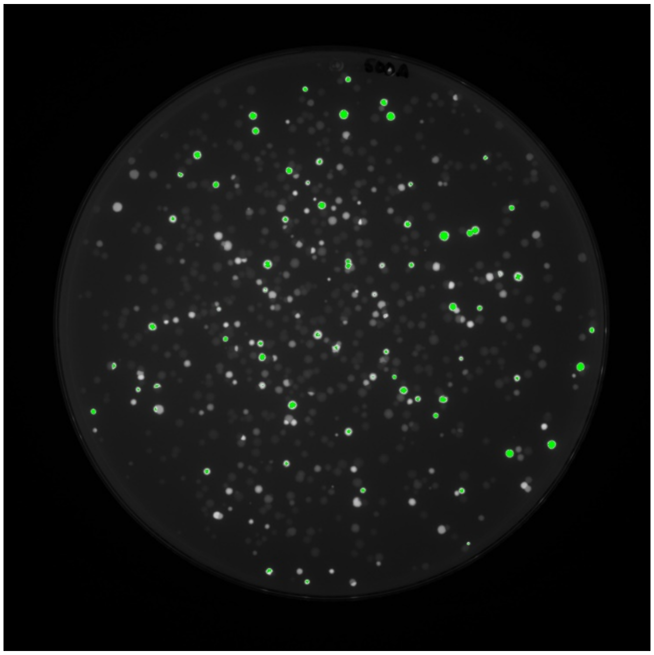

**Figure S12.** *E. coli* colonies expressing fluorescent library variants as imaged by the Azure400. The colonies exhibiting a fluorescence intensity greater than the average of the parent, represented by the green colonies, were selected for library expression and screening.

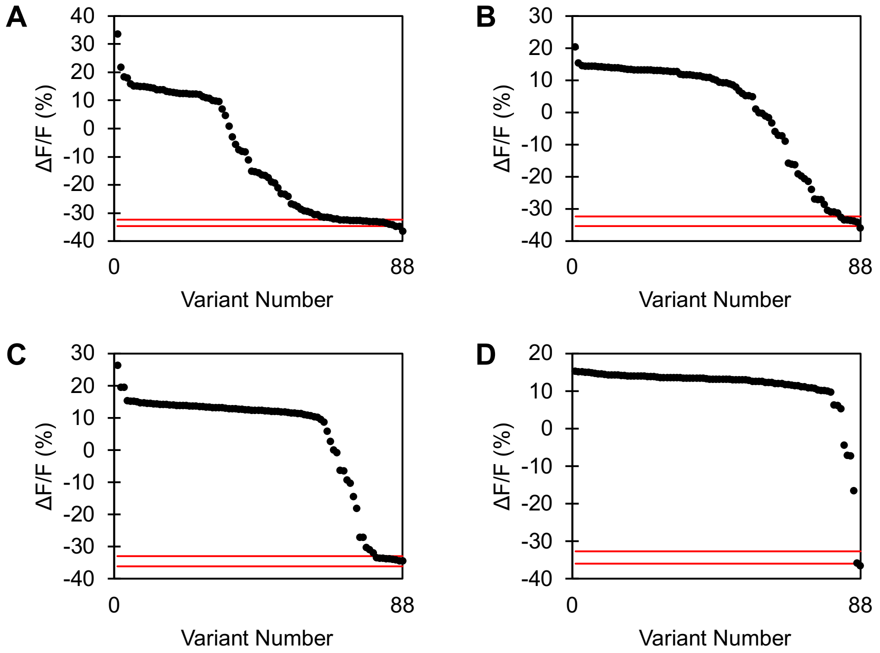


**Figure S13.** ROF plots from the third directed evolution library generated by using (A) 200, (B) 300, (C) 400, and (D) 500 μM manganese chloride during EP-PCR. Eighty-eight variants per manganese chloride concentration were screened in the absence ($F_i$) and presence ($F_f$) of 1 mM sodium nitrate in 50 mM phosphate buffer at pH 7.4. Variants that retained the function of the parent had a fluorescence response ($\Delta F/F = (F_f - F_i) / F_i * 100\%$) within three standard deviations of the average fluorescence response for six biological replicates of the 2-20C12 parent as indicated by the red lines. Based on these ROF plots, the libraries generated by using 300, 400, and 500 μM manganese chloride were combined and referred to as *Library 3* for further expression and screening.

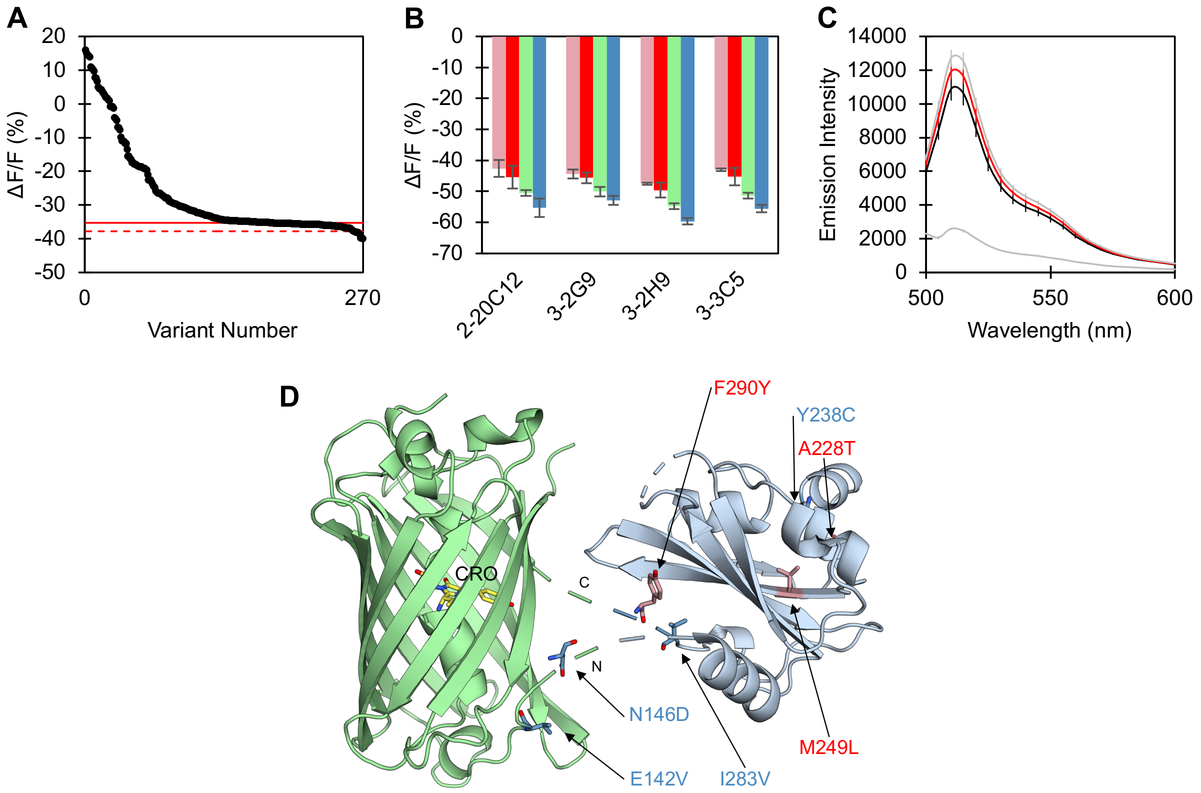


**Figure S14.** Screening of the third directed evolution library. (A) Summary plot of the third NreA domain mutagenesis library (*Library 3*) for the 264 variants (black circles) screened in the absence ($F_i$) and presence ($F_f$) of 1 mM sodium nitrate. The average fluorescence response ($\Delta F/F = (F_f - F_i) / F_i * 100\%$) of 24 biological replicates of the 2-20C12 parent ($\Delta F/F$ = -35.3% ± 0.8%) is shown by the blue dashed line. Three standard deviations from the parent average ($\Delta F/F$ = -37.7%) was used as a threshold to identify improved variants. The variant that was selected for rescreening and used as the template for the next EP-PCR library is indicated by the red circle. (B) Summary of the rescreen of the parent and top variants. The average fluorescence response with standard deviation to 5 (pink), 25 (red), 50 (green), and 100 (blue) mM sodium nitrate are shown for two biological replicates with two technical replicates each. (C) The average baseline emission spectra with standard deviation of the 2-20C12 parent (black) and top performing variant (red). The remaining top variants from (B) are shown in gray. Excitation was provided at 485 nm. All screening was done in *E. coli* lysate with 50 mM sodium phosphate buffer at pH 7.4. (D) AlphaFold model of the NreA-EGFP construct showing the position of all mutations (shown as sticks) in the 3-2H9 variant. The positions of the mutations are highlighted as pink sticks for new mutations and blue sticks for previous mutations with labels and the N- and C-terminal linker regions of the NreA domain are labeled as N and C, respectively. The EGFP chromophore (CRO) is shown as yellow sticks.

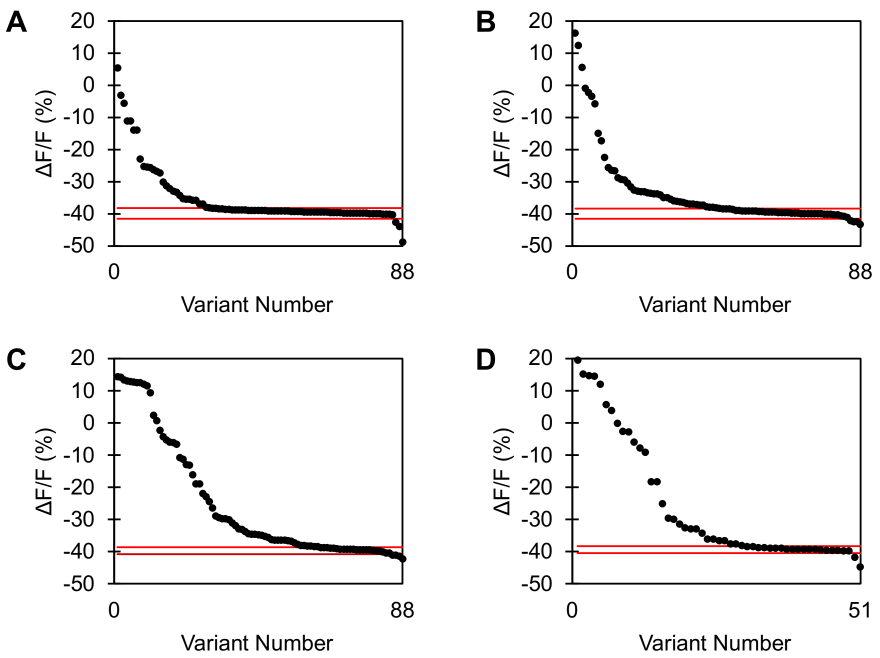


**Figure S15.** ROF plots from the fourth directed evolution library generated by using (A) 200, (B) 300, (C) 400, and (D) 500 µM manganese chloride during EP-PCR. Eighty-eight variants per manganese chloride concentration were screened in the absence ($F_i$) and presence ($F_f$) of 1 mM sodium nitrate in 50 mM phosphate buffer at pH 7.4. Variants that retained the function of the parent had a fluorescence response ($\Delta F/F = (F_f - F_i) / F_i * 100\%$) within three standard deviations of the average fluorescence response for six biological replicates of the 3-2H9 parent as indicated by the red lines. Based on these ROF plots, the libraries generated by using 300, 400, and 500 µM manganese chloride were combined and referred to as *Library 4* for further expression and screening.

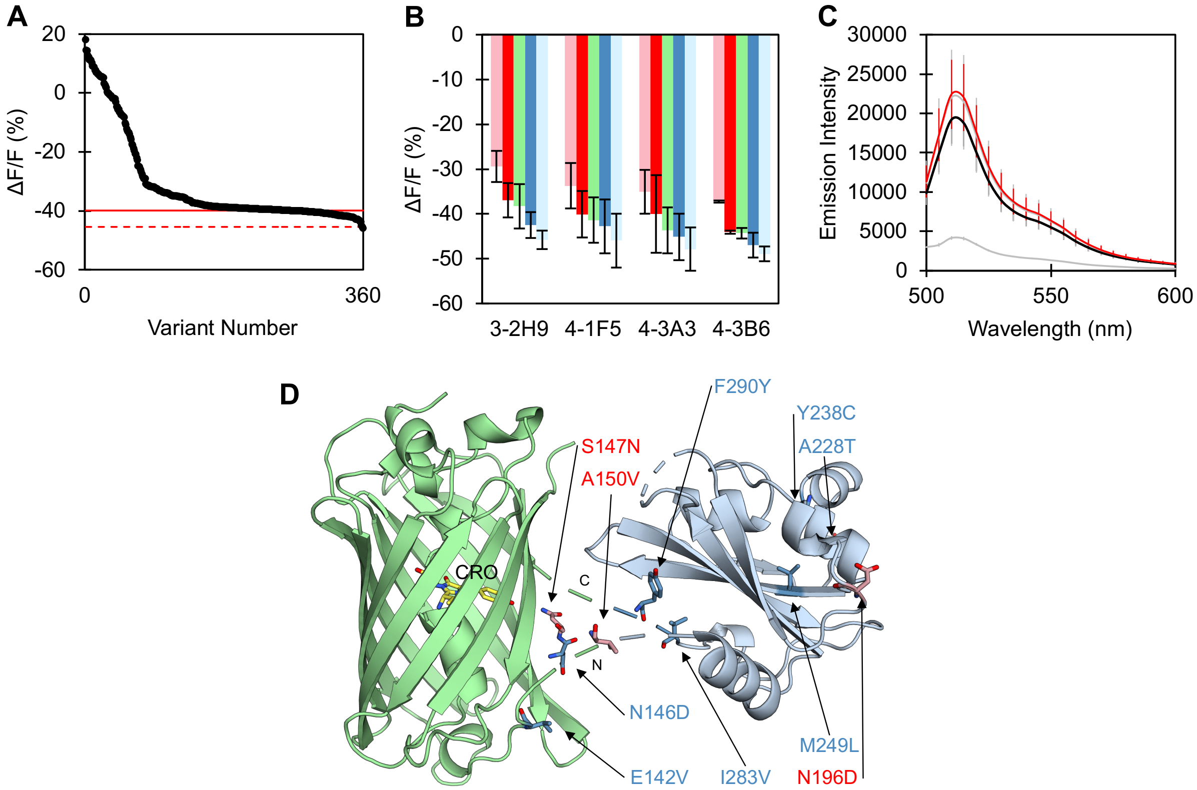


**Figure S16.** Screening of the fourth directed evolution library. (A) Summary plot of the fourth NreA domain mutagenesis library (*Library 4*) for the 352 variants (black circles) screened in the absence ($F_i$) and presence ($F_f$) of 1 mM sodium nitrate. The average fluorescence response ($\Delta F/F = (F_f - F_i) / F_i * 100\%$) of 32 biological replicates of the 3-2H9 parent (ΔF/F = -39.9% ± 1.8%) is shown by the blue dashed line. Three standard deviations from the parent average (ΔF/F = -45.3%) was used as a threshold to identify improved variants. The variant that was selected for rescreening and used as the template for the next EP-PCR library is indicated by the red circle. (B) Summary of the rescreen of the parent and top variants. The average fluorescence response with standard deviation to 0.1 (pink), 0.5 (red), 1 (green), 5 (blue), and 25 (light blue) mM sodium nitrate are shown for two biological replicates with two technical replicates each. (C) The average baseline emission spectra with standard deviation of the 3-2H9 parent (black) and top performing variant (red). The remaining top variants from (B) are shown in gray. Excitation was provided at 485 nm. All screening was done in *E. coli* lysate with 50 mM sodium phosphate buffer at pH 7.4. (D) AlphaFold model of the NreA-EGFP construct showing the position of all mutations (shown as sticks) in the 4-3A3 variant. The positions of the mutations are highlighted as pink sticks for new mutations and blue sticks for previous mutations with labels and the N- and C-terminal linker regions of the NreA domain are labeled as N and C, respectively. The EGFP chromophore (CRO) is shown as yellow sticks.

**Table S4.** Summary of top variants identified and rescreened from the mutagenesis of the NreA domain. The fluorescence response ($\Delta F/F = (F_f - F_i) / F_i * 100\%$) of the parent (italics) and top variants during library screening and rescreening in 50 mM phosphate buffer at pH 7.4 are shown. The top variant that was selected as the template for the next library is indicated by red font.

| | **Library 1** | | |
|---|---|---|---|
| **Variant** | **Library Screening ΔF/F for 100 mM $NO_3^-$** | **Rescreening ΔF/F for 50 mM $NO_3^-$** | **Mutations** |
| *NreA-EGFP* | -26.7 ± 4.4% | -23.0 ± 0.2% | - |
| **1-13E4** | -44.0% | -36.6 ± 0.3% | E142V, Y238C |
| | **Library 2** | | |
| **Variant** | **Library Screening ΔF/F for 50 mM $NO_3^-$** | **Rescreening ΔF/F for 1 mM $NO_3^-$** | **Mutations** |
| *1-13E4* | -31.1 ± 2.1% | -20.0 ± 0.1% | - |
| 2-10F5 | -40.8% | -34.3± 0.3% | N146D, A150V, V185A, N284S |
| 2-13F5 | -42.8% | -36.2 ± 0.1% | N146D, I252V |
| **2-20C12** | -41.6% | -35.4 ± 0.1% | N146D, I283V |
| | **Library 3** | | |
| **Variant** | **Library Screening ΔF/F for 1 mM $NO_3^-$** | **Rescreening ΔF/F for 1 mM $NO_3^-$** | **Mutations** |
| *2-20C12* | -35.3 ± 0.8% | -40.0 ± 3.2% | - |
| 3-2G9 | -39.8% | -41.6 ± 3.0% | E132G, N258S |
| **3-2H9** | -39.7% | -43.7 ± 1.6% | A228T, M249L, F290Y |
| 3-3C5 | -39.8% | -38.6 ± 2.4% | L181P |
| | **Library 4** | | |
| **Variant** | **Library Screening ΔF/F 1 mM $NO_3^-$** | **Rescreening ΔF/F for 1 mM $NO_3^-$** | **Mutations** |
| *3-2H9* | -39.9 ± 1.8% | -38.3 ± 5.0% | - |
| 4-1A8 | -45.5% | -45.1 ± 7.9% | A159V, A263V |
| 4-1F5 | -45.0% | -41.4 ± 5.1% | L181P |
| **4-3A3** | -43.7% | -43.7 ± 5.2% | S147N, A150V, N196D |
| 4-3B6 | -45.8% | -44.3 ± 1.2% | D161E, T280S, S289R |

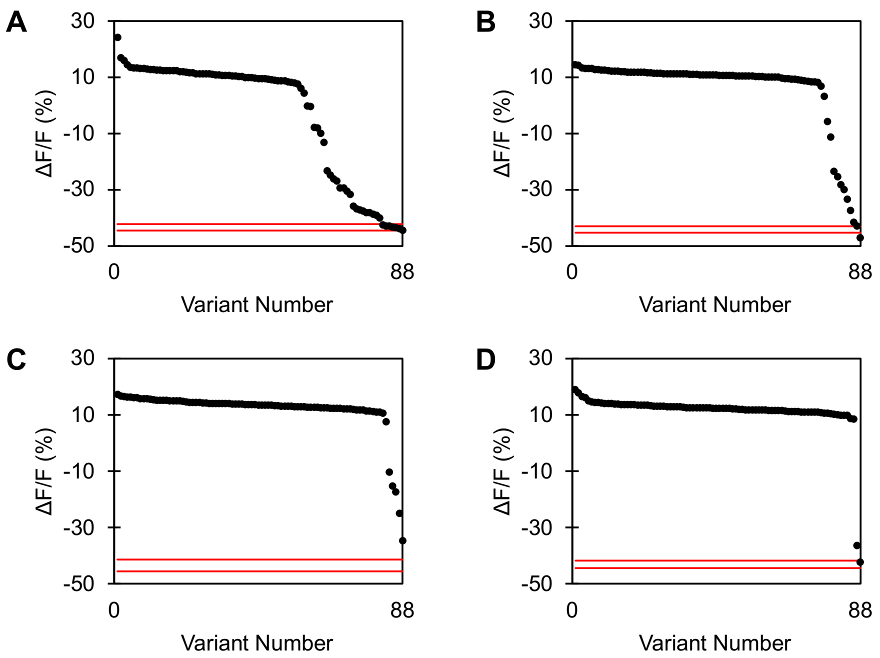


**Figure S17.** ROF plots from the fifth directed evolution library generated by using (A) 200, (B) 300, (C) 400, and (D) 500 μM manganese chloride during EP-PCR. Eighty-eight variants per manganese chloride concentration were screened in the absence ($F_i$) and presence ($F_f$) of 1 mM sodium nitrate in 50 mM phosphate buffer at pH 7.4. Variants that retained the function of the parent had a fluorescence response ($\Delta F/F = (F_f - F_i) / F_i * 100\%$) within three standard deviations of the average fluorescence response for six biological replicates of the 4-3A3 parent as indicated by the red lines. Based on these ROF plots, the libraries generated by using 200 and 300 μM manganese chloride were combined and referred to as *Library 5* for further expression and screening.

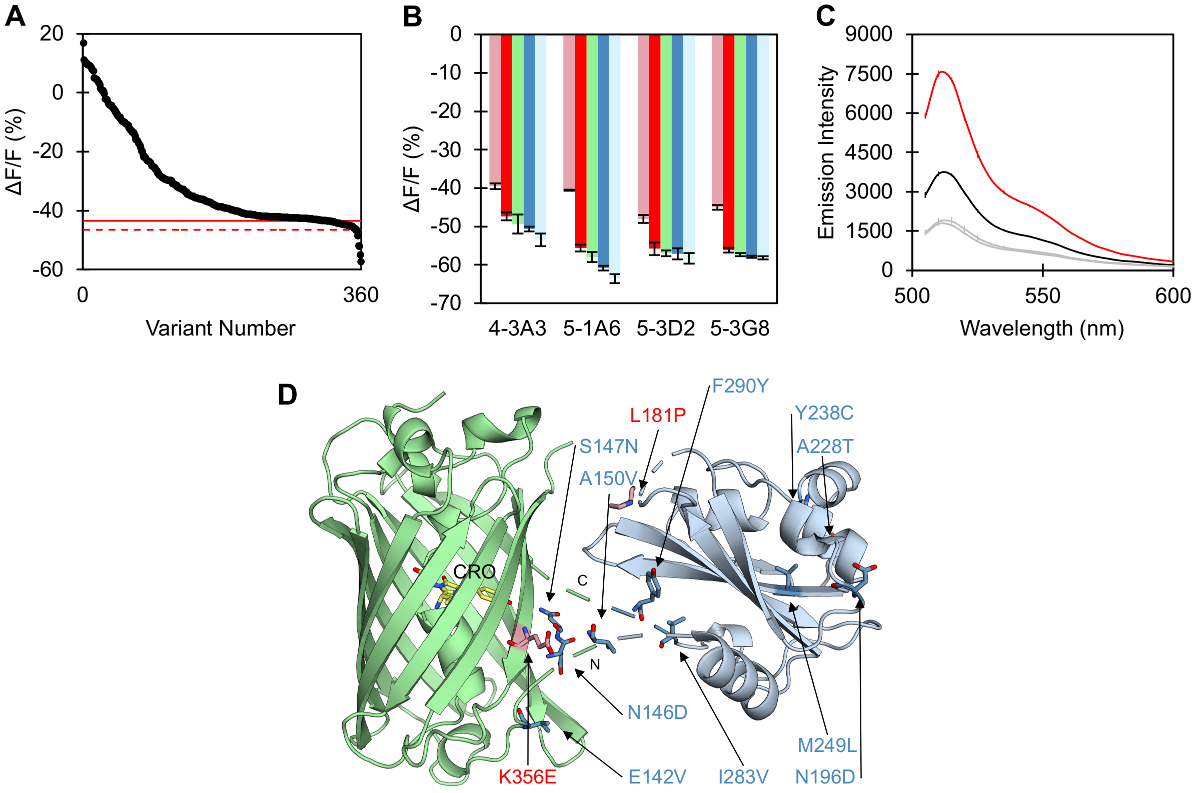


**Figure S18.** Screening of the fifth directed evolution library. (A) Summary plot of the first NreA-EGFP mutagenesis library (*Library 5*) for the 352 variants (black circles) screened in the absence ($F_i$) and presence ($F_f$) of 1 mM sodium nitrate. The average fluorescence response ($\Delta F/F = (F_f - F_i) / F_i * 100\%$) of 32 biological replicates of the 4-3A3 parent ($\Delta F/F$ = -43.5% ± 1.0%) is shown by the blue dashed line. Three standard deviations from the parent average ($\Delta F/F$ = -46.5%) was used as a threshold to identify improved variants. The variant that was selected for rescreening and used as the template for the next EP-PCR library is indicated by the red circle. (B) Summary of the rescreen of the parent and top variants. The average fluorescence response with standard deviation to 0.1 (pink), 0.5 (red), 1 (green), 5 (blue), and 25 (light blue) mM sodium nitrate are shown for two biological replicates with two technical replicates each. (C) The average baseline emission spectra with standard deviation of the 4-3A3 parent (black) and top performing variant (red). The remaining top variants from (B) are shown in gray. Excitation was provided at 485 nm. All screening was done in *E. coli* lysate with 50 mM sodium phosphate buffer at pH 7.4. (D) AlphaFold model of the NreA-EGFP construct showing the position of all mutations (shown as sticks) in the 5-1A6 variant. The positions of the mutations are highlighted as pink sticks for new mutations and blue sticks for previous mutations with labels and the N- and C-terminal linker regions of the NreA domain are labeled as N and C, respectively. The EGFP chromophore (CRO) is shown as yellow sticks.

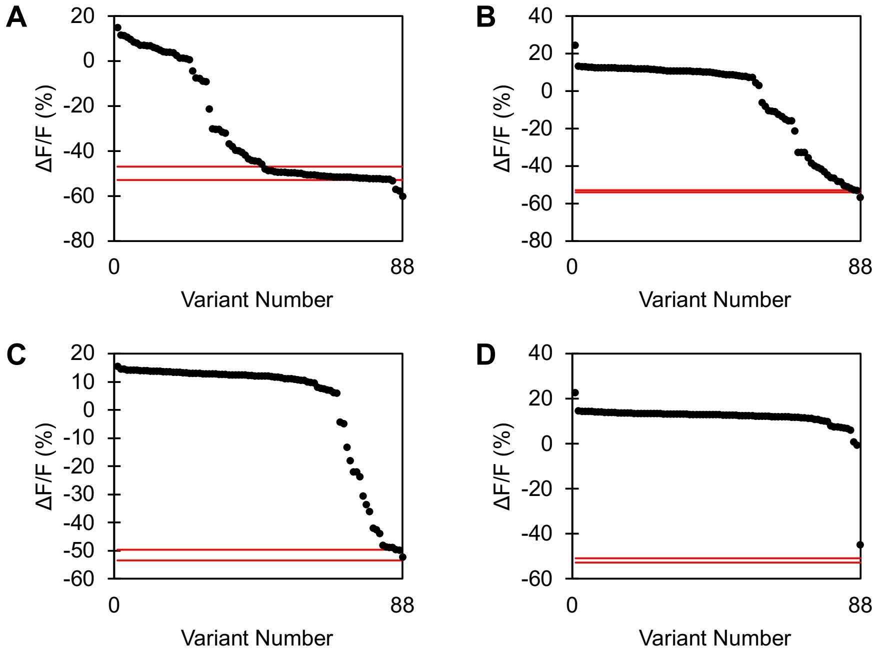


**Figure S19.** ROF plots from the sixth directed evolution library generated by using (A) 100, (B) 200, (C) 300, and (D) 400 µM manganese chloride during EP-PCR. Eighty-eight variants per manganese chloride concentration were screened in the absence ($F_i$) and presence ($F_f$) of 1 mM sodium nitrate in 50 mM phosphate buffer at pH 7.4. Variants that retained the function of the parent had a fluorescence response ($\Delta F/F = (F_f - F_i) / F_i * 100\%$) within three standard deviations of the average fluorescence response for six biological replicates of the 5-1A6 parent as indicated by the red lines. Based on these ROF plots, the libraries generated by using 100 and 200 µM manganese chloride were combined and referred to as *Library 6* for further expression and screening.

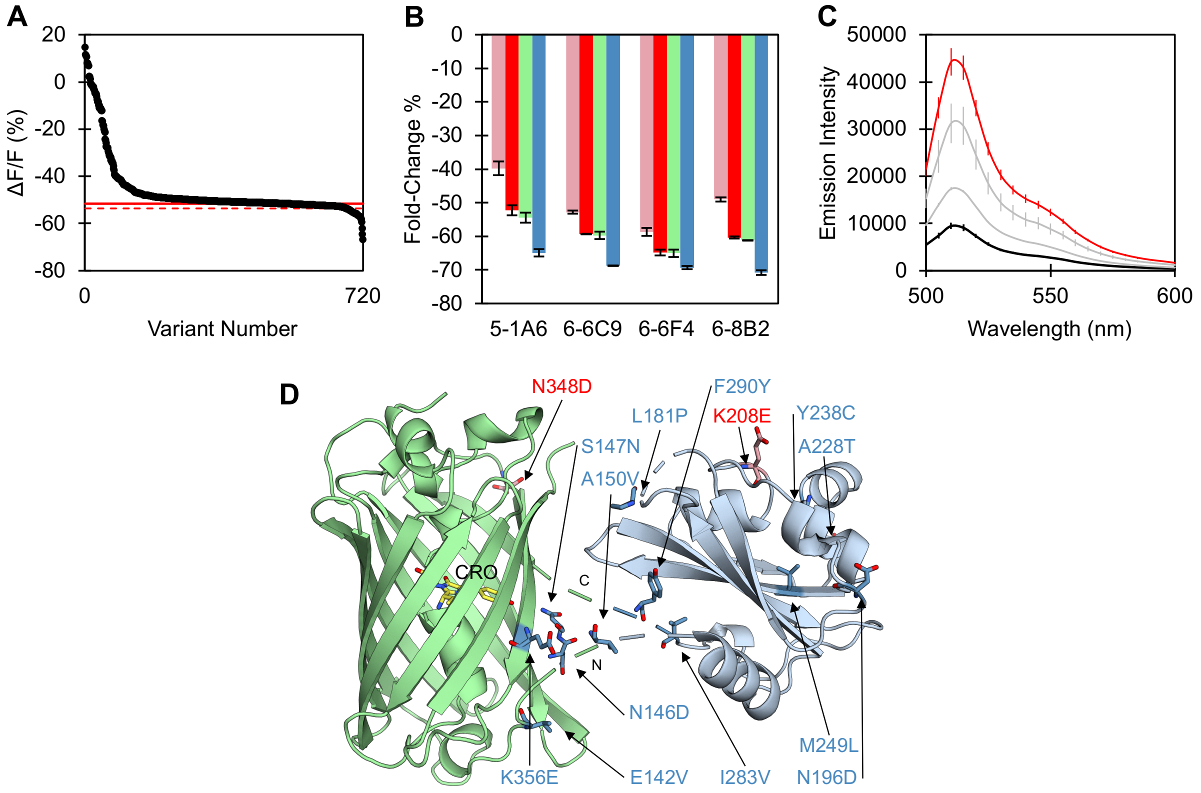


**Figure S20.** Screening of the sixth directed evolution library. (A) Summary plot of the second NreA-EGFP mutagenesis library (*Library 6*) for the 704 variants (black circles) screened in the absence ($F_i$) and presence ($F_f$) of 1 mM sodium nitrate. The average fluorescence response ($\Delta F/F = (F_f - F_i) / F_i * 100\%$) of 64 biological replicates of the 5-1A6 parent ($\Delta F/F$ = -51.6% ± 0.7%) is shown by the blue dashed line. Three standard deviations from the parent average ($\Delta F/F$ = -53.7%) was used as a threshold to identify improved variants. The variant that was selected for rescreening and used as the template for the next EP-PCR library is indicated by the red circle. (B) Summary of the rescreen of the parent and top variants. The average fluorescence response with standard deviation to 0.1 (pink), 0.5 (red), 1 (green), and 50 (blue) mM sodium nitrate are shown for two biological replicates with two technical replicates each. (C) The average baseline emission spectra with standard deviation of the 5-1A6 parent (black) and top performing variant (red). The remaining top variants from (B) are shown in gray. Excitation was provided at 485 nm. All screening was done in *E. coli* lysate with 50 mM sodium phosphate buffer at pH 7.4. (D) AlphaFold model of the NreA-EGFP construct showing the position of all mutations (shown as sticks) in the 6-6C9 variant. The positions of the mutations are highlighted as pink sticks for new mutations and blue sticks for previous mutations with labels and the N- and C-terminal linker regions of the NreA domain are labeled as N and C, respectively. The EGFP chromophore (CRO) is shown as yellow sticks.

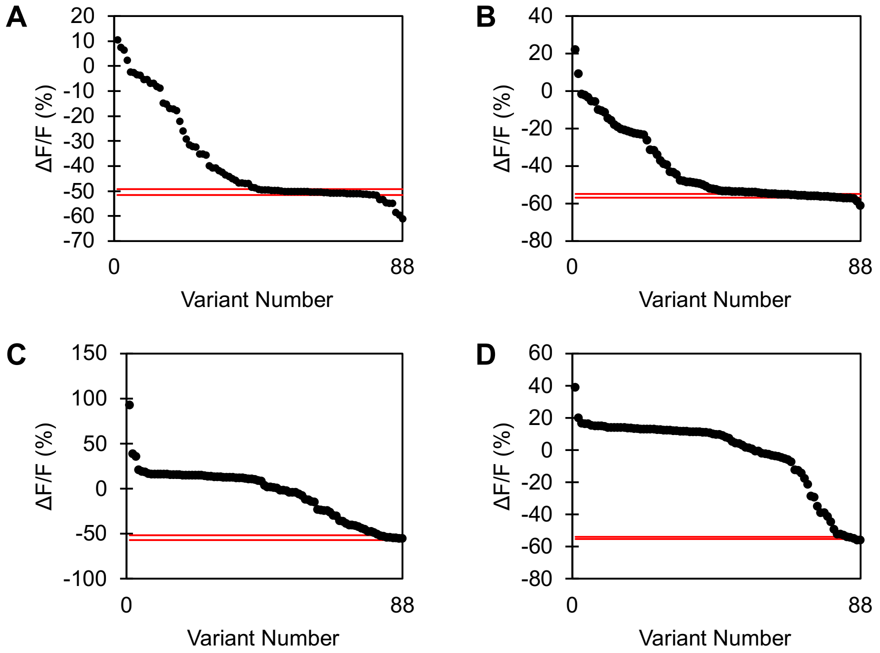


**Figure S21.** ROF plots from the seventh directed evolution library generated by using (A) 50, (B) 100, (C) 200, and (D) 300 μM manganese chloride during EP-PCR. Eighty-eight variants per manganese chloride concentration were screened in the absence ($F_i$) and presence ($F_f$) of 1 mM sodium nitrate in 50 mM phosphate buffer at pH 7.4. Variants that retained the function of the parent had a fluorescence response ($\Delta F/F = (F_f - F_i) / F_i * 100\%$) within three standard deviations of the average fluorescence response for six biological replicates of the 6-6C9 parent as indicated by the red lines. Based on these ROF plots, the libraries generated by using 50 and 100 μM manganese chloride were combined and referred to as *Library 7* for further expression and screening.

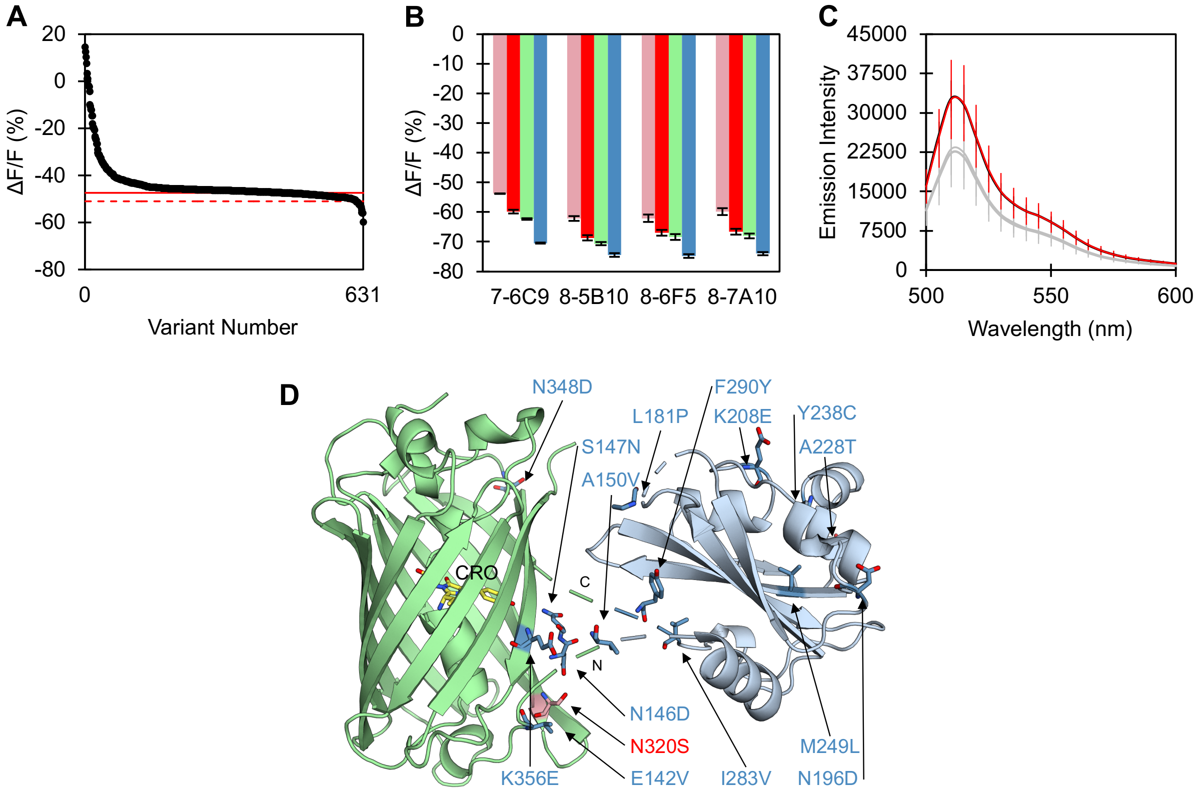


**Figure S22.** Screening of the seventh directed evolution library. (A) Summary plot of the third NreA-EGFP mutagenesis library (*Library 7*) for the 616 variants (black circles) screened in the absence ($F_i$) and presence ($F_f$) of 0.1 mM sodium nitrate. The average fluorescence response ($\Delta F/F = (F_f - F_i) / F_i * 100\%$) of 56 biological replicates of the 6-6C9 parent ($\Delta F/F$ = -47.5% ± 1.2%) is shown by the red line. Three standard deviations from the parent average ($\Delta F/F$ = -51.1%, red dashed line) was used as a threshold to identify improved variants. The variant that was selected for rescreening and used as the template for the next EP-PCR library is indicated by the red circle. (B) Summary of the rescreen of the parent and top variants. The average fluorescence response with standard deviation to 0.1 (pink), 0.5 (red), 1 (green), and 50 (blue) mM sodium nitrate is shown for two biological replicates with two technical replicates each. (C) The average baseline emission spectra with standard deviation of the 6-6C9 parent (black) and top performing variant (red). The remaining top variants from (B) are shown in gray. Excitation was provided at 485 nm. All screening was done in *E. coli* lysate with 50 mM sodium phosphate buffer at pH 7.4. (D) AlphaFold model of the NreA-EGFP construct showing the position of all mutations (shown as sticks) in the final 7-7A10 variant named NitrOFF. The positions of the mutations are highlighted as pink sticks for new mutations and blue sticks for previous mutations with labels and the N- and C-terminal linker regions of the NreA domain are labeled as N and C, respectively. The EGFP chromophore (CRO) is shown as yellow sticks.

**Table S5.** Summary of top variants identified and rescreened from the mutagenesis of the entire NreA-EGFP construct. The fluorescence response ($\Delta F/F = (F_f - F_i) / F_i * 100\%$) of the template (italics) and top variants during library screening and rescreening in 50 mM phosphate buffer at pH 7.4 are shown. The top variant that was selected as the template for the next library, or as the final variant in the case of Library 7, is indicated by red text.

| Library 5 | | | |
|---|---|---|---|
| **Variant** | **Library Screening $\Delta F/F$ for 1 mM $NO_3^-$** | **Rescreening $\Delta F/F$ for 1 mM $NO_3^-$** | **Mutations** |
| *4-3A3* | -43.5 ± 1.0% | -49.4 ± 2.5% | - |
| **5-1A6** | -52.1% | -57.9 ± 1.3% | L181P, K356E |
| 5-3D2 | -55.0% | -57.0 ± 0.8% | N258S, F281L, S289N |
| 5-3D5 | -52.0% | -43.1 ± 2.9% | L181P, K303E, S352G, V374A |
| 5-3G8 | -57.4% | -57.4 ± 0.4% | Q285E, G291D, R318C, D347N, K356E |
| **Library 6** | | | |
| **Variant** | **Library Screening $\Delta F/F$ for 1 mM $NO_3^-$** | **Rescreening $\Delta F/F$ for 0.1 mM $NO_3^-$** | **Mutations** |
| *5-1A6* | -51.6 ± 0.7% | -39.8 ± 2.0% | - |
| **6-6C9** | -57.6% | -52.8 ± 0.5% | K208E, N348D |
| 6-6F4 | -64.6% | -58.7 ± 1.2% | E5D, I14V, T43A, K208E, N299T, R318E |
| 6-8B2 | -58.2% | -50.0 ± 0.6% | E5A, V142A, F372Y |
| **Library 7** | | | |
| **Variant** | **Library Screening $\Delta F/F$ for 0.1 mM $NO_3^-$** | **Rescreening $\Delta F/F$ for 0.1 mM $NO_3^-$** | **Mutations** |
| *6-6C9* | -47.5 ± 1.2% | -53.8 ± 0.1% | - |
| 7-2G11 | -56.0% | -62.1 ± 2.4% | [a] |
| 7-4B6 | -60.0% | -62.1 ± 0.3% | [a] |
| 7-5B10 | -56.2% | -62.2 ± 0.8% | [a] |
| 7-6F5 | -56.1% | -62.1 ± 1.3% | [a] |
| **7-7A10 (NitrOFF)** | -54.8% | -60.0 ± 1.2% | N320S |

[a] Sequences were not determined for these variants.

```
CAT ATG GTT AGC AAG GGC GAG GAG CTG TTT ACC GGC GTG GTT CCG ATT CTG GTC GAG CTG
His Met Val Ser Lys Gly Glu Glu Leu Phe Thr Gly Val Val Pro Ile Leu Val Glu Leu   20

GAT GGT GAT GTT AAT GGT CAT AAG TTT AGC GTT AGC GGC GAG GGC GAA GGT GAC GCG ACC
Asp Gly Asp Val Asn Gly His Lys Phe Ser Val Ser Gly Glu Gly Glu Gly Asp Ala Thr   40

TAC GGC AAG CTG ACC CTG AAA TTC ATC TGC ACC ACC GGT AAA CTG CCG GTG CCG TGG CCG
Tyr Gly Lys Leu Thr Leu Lys Phe Ile Cys Thr Thr Gly Lys Leu Pro Val Pro Trp Pro   60

ACC CTG GTT ACC ACC CTG ACC TAC GGC GTG CAG TGC TTT AGC CGT TAT CCG GAC CAC ATG
Thr Leu Val Thr Thr Leu Thr Tyr Gly Val Gln Cys Phe Ser Arg Tyr Pro Asp His Met   80

AAG CAA CAC GAT TTC TTT AAA AGC GCG ATG CCG GAG GGC TAC ATC CAG GAA CGT ACC ATT
Lys Gln His Asp Phe Phe Lys Ser Ala Met Pro Glu Gly Tyr Ile Gln Glu Arg Thr Ile   100

TTC TTT AAG GAC GAT GGT AAC TAC AAG ACC CGT GCG GAA GTG AAG TTC GAA GGT GAT ACC
Phe Phe Lys Asp Asp Gly Asn Tyr Lys Thr Arg Ala Glu Val Lys Phe Glu Gly Asp Thr   120

CTG GTT AAC CGT ATC GAG CTG AAG GGT ATT GAC TTT AAA GAA GAT GGC AAC ATC CTG GGT
Leu Val Asn Arg Ile Glu Leu Lys Gly Ile Asp Phe Lys Glu Asp Gly Asn Ile Leu Gly   140

CAC AAG CTT GTG TAC AAC CTG GAC AAC GTG ATC GTG AGC GAC TAC TTC GAT TAT CAG GAC
His Lys Leu Val Tyr Asn Leu Asp Asn Val Ile Val Ser Asp Tyr Phe Asp Tyr Gln Asp   160

GCG CTG GAT GAG ATC CGT GAG ACT GAA AAG TTC GAC TTT GCG GCG ATT GCG CTG CCG GAA
Ala Leu Asp Glu Ile Arg Glu Thr Glu Lys Phe Asp Phe Ala Ala Ile Ala Leu Pro Glu   180

GAT GGT CCG CAC AGC GCG GTT ATT AAG TGG AAA TAC GCG AGC GGC AAC ATC GAC TAC CGT
Asp Gly Pro His Ser Ala Val Ile Lys Trp Lys Tyr Ala Ser Gly Asn Ile Asp Tyr Arg   200

TAT CGT ATG ATT GTG CTG CGT CCG GGC GAG GGA CTG GCG GGT CTG GTT ATC CGT ACC GGC
Tyr Arg Met Ile Val Leu Arg Pro Gly Glu Gly Leu Ala Gly Leu Val Ile Arg Thr Gly   220

AGC CGT AAA ATT GTG GAG GAC GTT GAT ACG GAA CTG AGC CAG AAC GAC AAG CTG GGT TGT
Ser Arg Lys Ile Val Glu Asp Val Asp Thr Glu Leu Ser Gln Asn Asp Lys Leu Gly Cys   240

CCG ATT GTG CTG AGC GAG GCG CTG ACC GCG TTG GTT GCG ATT CCG CTG TGG AAA AAC AAC
Pro Ile Val Leu Ser Glu Ala Leu Thr Ala Leu Val Ala Ile Pro Leu Trp Lys Asn Asn   260

CGT GTG TAC GGT GCG CTG CTG CTG GGT CAA CGT GAA GGT CGC CCG CTG CCA GAA GGT AGC
Arg Val Tyr Gly Ala Leu Leu Leu Gly Gln Arg Glu Gly Arg Pro Leu Pro Glu Gly Ser   280

ACC ACC TTC CGT GTC AAT CAG CGT CTG GGC AGC TAT ACC GAT GAA ATT AAC AAA CAA GGC
Thr Thr Phe Arg Val Asn Gln Arg Leu Gly Ser Tyr Thr Asp Glu Ile Asn Lys Gln Gly   300

AAC GTG TAC ATC AAG GCG GAC AAG CAG AAA AAC GGT ATT AAG GCG AAC TTC AAA ATC CGT
Asn Val Tyr Ile Lys Ala Asp Lys Gln Lys Asn Gly Ile Lys Ala Asn Phe Lys Ile Arg   320

CAC AGC ATT GAA GAT GGT GGC GTT CAA CTG GCG TAC CAC TAT CAG CAA AAC ACC CCG ATT
His Ser Ile Glu Asp Gly Gly Val Gln Leu Ala Tyr His Tyr Gln Gln Asn Thr Pro Ile   340

GGT GAT GGT CCG GTG CTG CTG CCG GAT GAC CAC TAT CTG AGC GTT CAA AGC GAG CTG AGC
Gly Asp Gly Pro Val Leu Leu Pro Asp Asp His Tyr Leu Ser Val Gln Ser Glu Leu Ser   360

AAA GAC CCG AAC GAG AAG CGT GAT CAC ATG GTG TTG CTG GAA TTT GTT ACC GCG GCG GGC
Lys Asp Pro Asn Glu Lys Arg Asp His Met Val Leu Leu Glu Phe Val Thr Ala Ala Gly   380

ATT ACC CTG GGC ATG GAC GAA CTG TAT AAG GCG GCC GCA CTC GAG CAC CAC CAC CAC CAC
Ile Thr Leu Gly Met Asp Glu Leu Tyr Lys Ala Ala Ala Leu Glu His His His His His   400

CAC TGA
His End   402
```

**Figure S23.** Nucleotide (top row) and amino acid (bottom row) sequence of the NitrOFF construct with the mEGFP and NreA domains shown in green and black font, respectively. Mutations accumulated during the directed evolution (red), NdeI and NotI restriction enzyme sites from the pET-21a(+) vector (gray), polyhistidine tag (blue) and stop codon (pink) are shown.

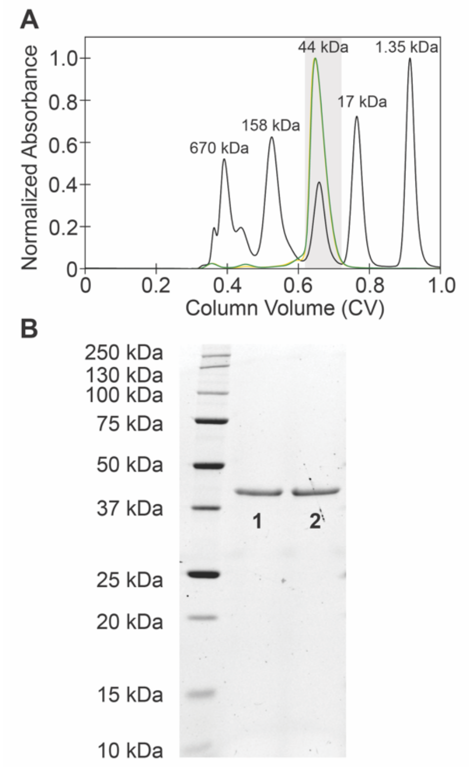


**Figure S24.** (A) Size exclusion chromatogram (SEC, normalized absorbance at 280 nm) for the first (orange) and second (green) biological replicate of the NitrOFF. The gel filtration standard is shown in black and made up of the following: thyroglobulin (670 kDa), $\gamma$-globulin (158 kDa), ovalbumin (44 kDa), myoglobulin (17 kDa), and vitamin B12 (1.35 kDa). All SECs were collected in 20 mM Tris buffer at pH 7.5 with 150 mM sodium chloride. For each protein run, the fraction collected for further analysis is shown in gray area. (B) Representative SDS-PAGE of the protein batches tested are shown in lanes 1 and 2. The theoretical molecular weight of NitrOFF is ~44.9 kDa.

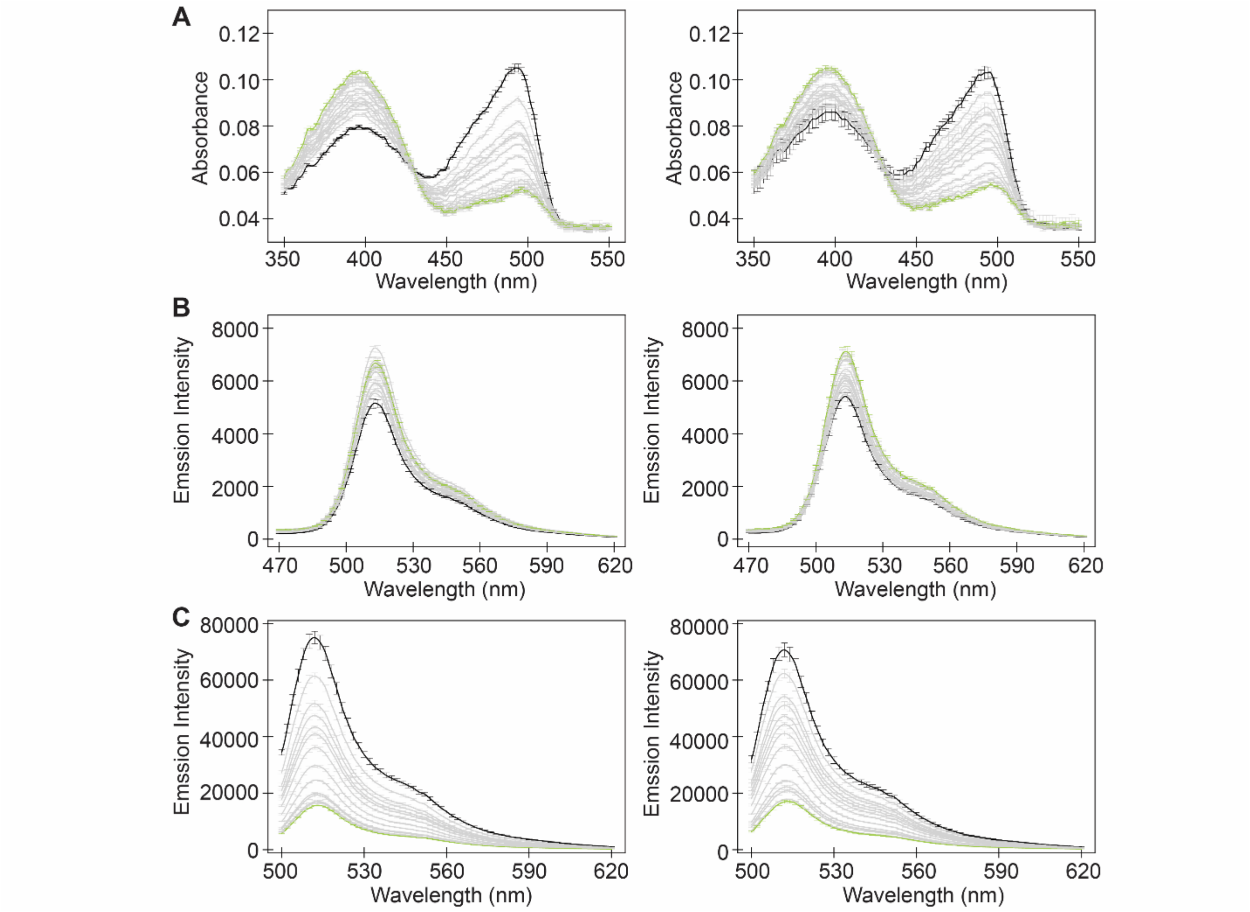


**Figure S25.** Spectroscopic characterization of NreA-EGFP with nitrate. Absorbance and emission spectra of ~4 µM NreA-EGFP excited at (A) 400 nm and (B) 480 nm in the presence 0 mM (black), 2, 4, 6, 8, 10, 15, 25, 50, 75, 100, 200, 250 (gray) or 500 µM (green) sodium nitrate in 25 mM sodium phosphate buffer at pH 7 with 1 mM sodium chloride. The average of three technical replicates with standard deviation is shown for one of two protein batches in each column.

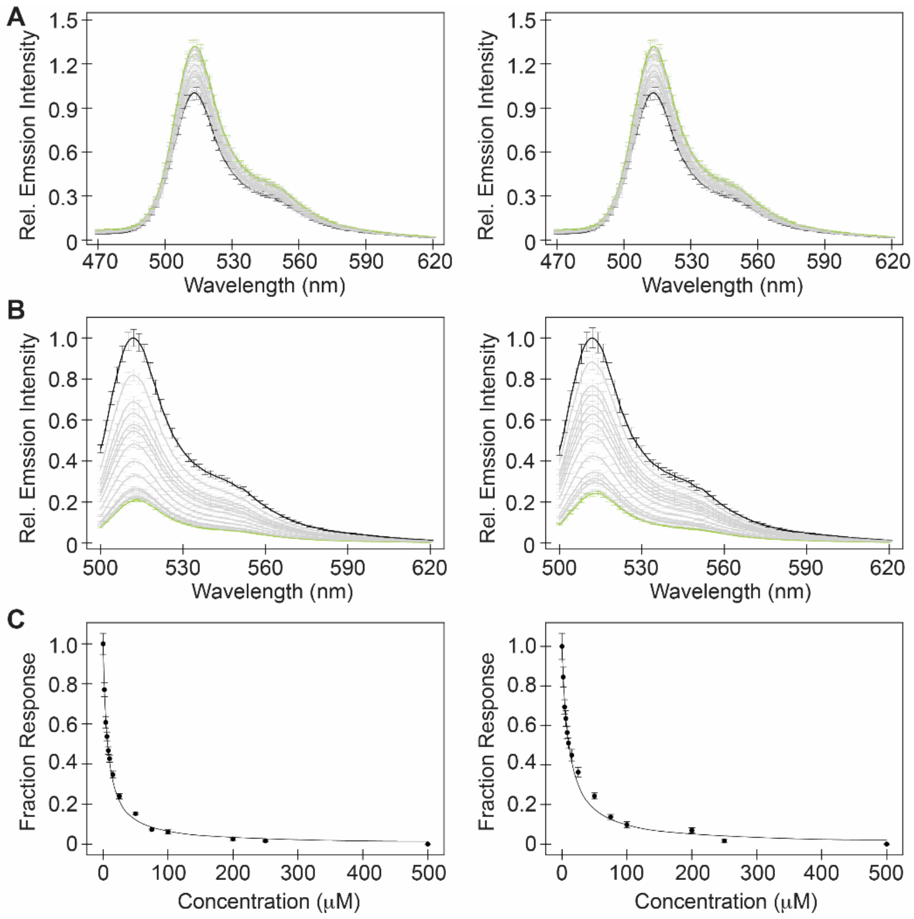


**Figure S26.** Spectroscopic characterization of NitrOFF with nitrate. Emission spectra of ~4 μM NitrOFF excited at (A) 400 nm and (B) 480 nm in the presence of 0 mM (black), 2, 4, 6, 8, 10, 15, 25, 50, 75, 100, 200, 250 (gray) or 500 μM (green) sodium nitrate in 25 mM sodium phosphate buffer at pH 7 with 1 mM sodium chloride. (C) The emission response at 514 nm from (B) was fitted to determine the apparent dissociation constant ($K_d$) of 6.9 ± 0.2 μM and 10.8 ± 0.6 μM. The average of three technical replicates with standard deviation is shown for one of two protein batches in each column.

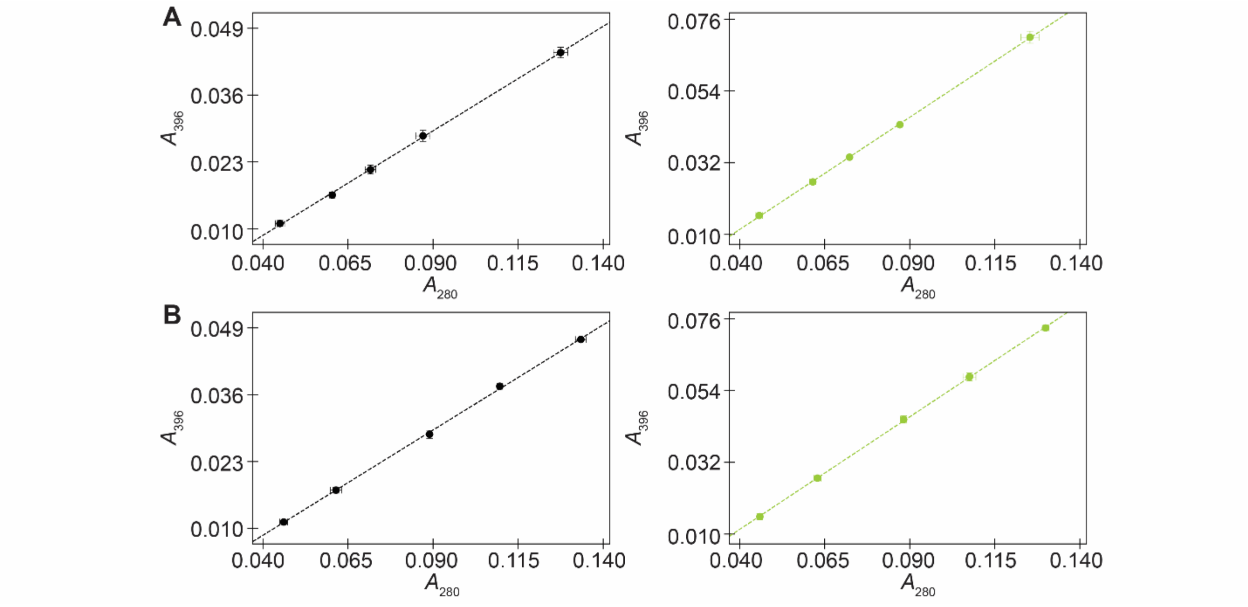


**Figure S27.** Extinction coefficient curve for NitrOFF in the absence (left) and presence (right) of 0.5 mM sodium nitrate in 25 mM sodium phosphate buffer at pH 7 with 1 mM sodium chloride. The absorbance intensity at 396 nm ($A_{396}$) is plotted versus the absorbance intensity at 280 nm ($A_{280}$) ($R^2$ > 0.99). Data from the (A) first and (B) second protein batch is plotted as the average of three measurements with standard deviation.

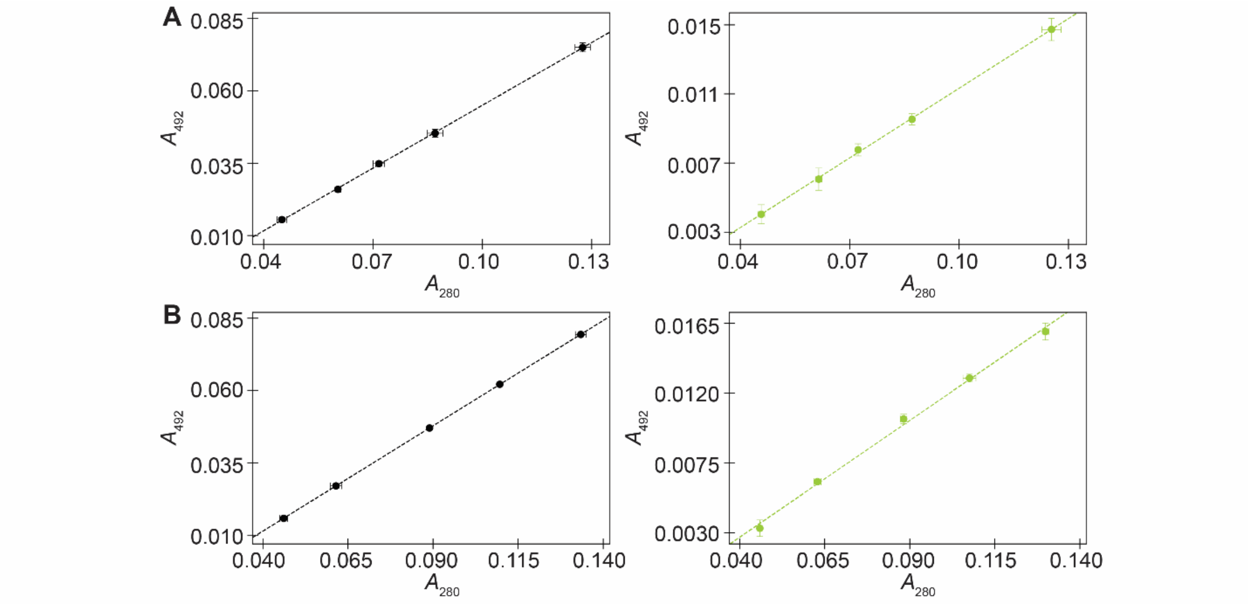


**Figure S28.** Extinction coefficient curve for NitrOFF in the absence (left) and presence (right) of 0.5 mM sodium nitrate in 25 mM sodium phosphate buffer at pH 7 with 1 mM sodium chloride. The absorbance intensity at 492 nm ($A_{492}$) is plotted versus the absorbance intensity at 280 nm ($A_{280}$) ($R^2 > 0.99$). Data from the (A) first and (B) second protein batch is plotted as the average of three measurements with standard deviation.

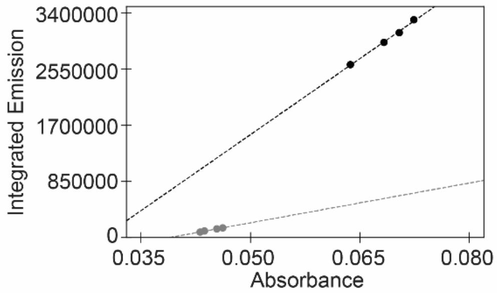


**Figure S29**. Fluorescence quantum yield standard curves for fluorescein (black, in 0.1 M NaOH) and Coumarin 135 (gray, in 50% ethanol). The integrated emission is plotted versus the corrected absorbance intensity at 488 and 425 nm, respectively ($R^2 > 0.99$). Data for each concentration was plotted as the average of three measurements with standard deviation and all experiments were conducted at room temperature (23–25 °C).

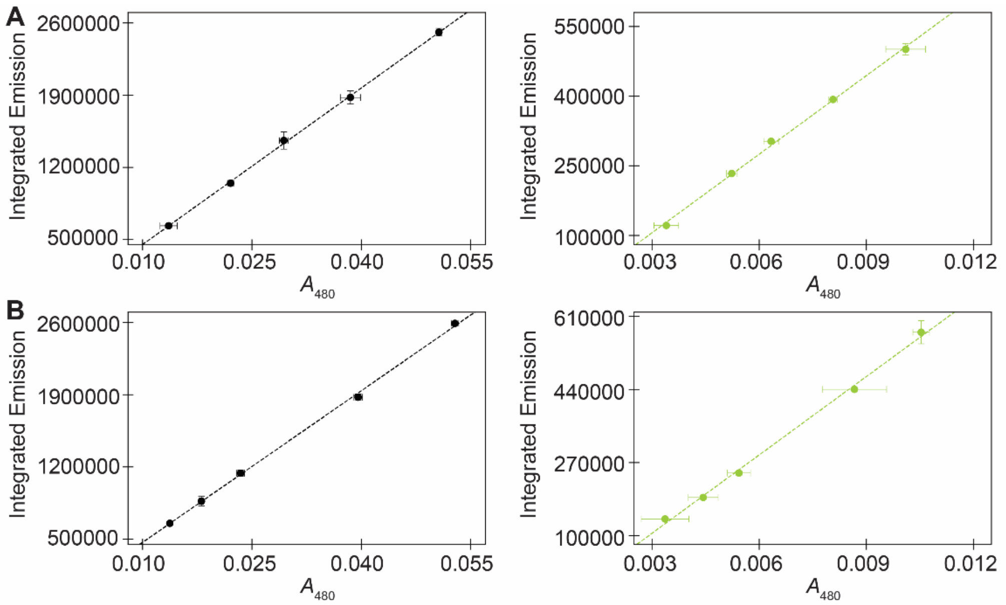


**Figure S30.** Fluorescence quantum yield curves for NitrOFF in the absence (left) and presence (right) of 0.5 mM sodium nitrate in 25 mM sodium phosphate buffer at pH 7 with 1 mM sodium chloride. The integrated emission of each sample is plotted versus the corresponding absorbance intensity at 480 nm ($A_{488}$) ($R^2$ > 0.99). Data from the (A) first and (B) second protein batch is plotted as the average of three measurements with standard deviation.

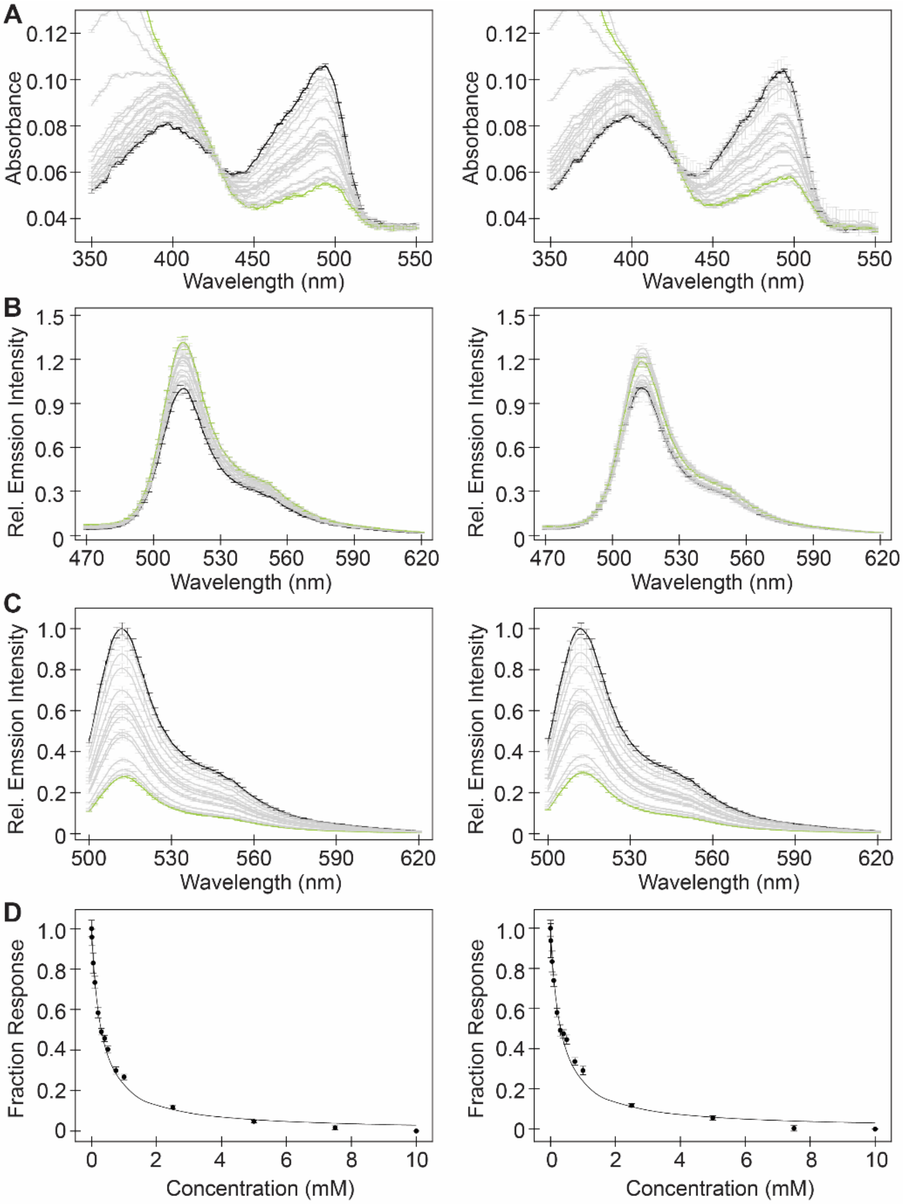


**Figure S31.** Spectroscopic characterization of NitrOFF with nitrite. (A) Absorbance and emission spectra of ~4 µM NitrOFF excited at (B) 400 nm and (C) 480 nm in the presence of 0 mM (black), 0.01, 0.05, 0.1, 0.2, 0.3, 0.4, 0.5, 0.75, 1, 2.5 5, 7.5 (gray) or 10 mM (green) sodium nitrite in 25 mM sodium phosphate buffer at pH 7 with 1 mM sodium chloride. (D) The emission response at 514 nm from *(C)* was fitted to determine the apparent dissociation constant ($K_d$) of 300.8 ± 9.4 µM and 317.5 ± 16.7 µM. The average of three technical replicates with standard deviation is shown for one of two protein batches in each column.

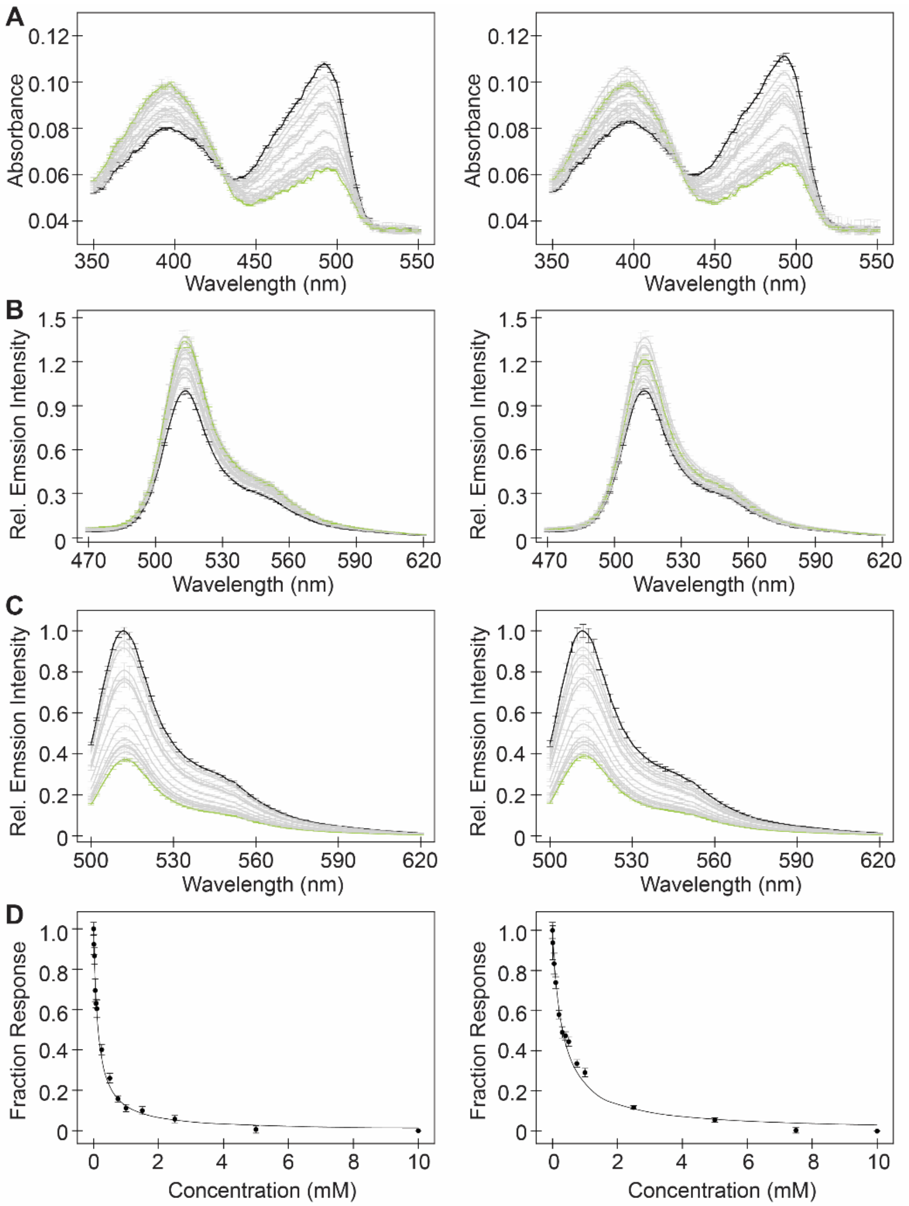


**Figure S32.** Spectroscopic characterization of NitrOFF with iodide. Emission spectra of ~4 µM NitrOFF excited at (A) 400 nm and (B) 480 nm in the presence of 0 mM (black), 0.01, 0.025, 0.05, 0.075, 0.1, 0.25, 0.5, 0.75, 1, 1.5, 2.5, 5 (gray) or 10 mM (green) sodium iodide in 25 mM sodium phosphate buffer at pH 7 with 1 mM sodium chloride. (C) The emission response at 514 nm from *(C)* was fitted to determine the apparent dissociation constant ($K_d$) of 140.8 ± 8.5 µM and 131.4 ± 8.1 µM. The average of three technical replicates with standard deviation is shown for one of two protein batches in each column.

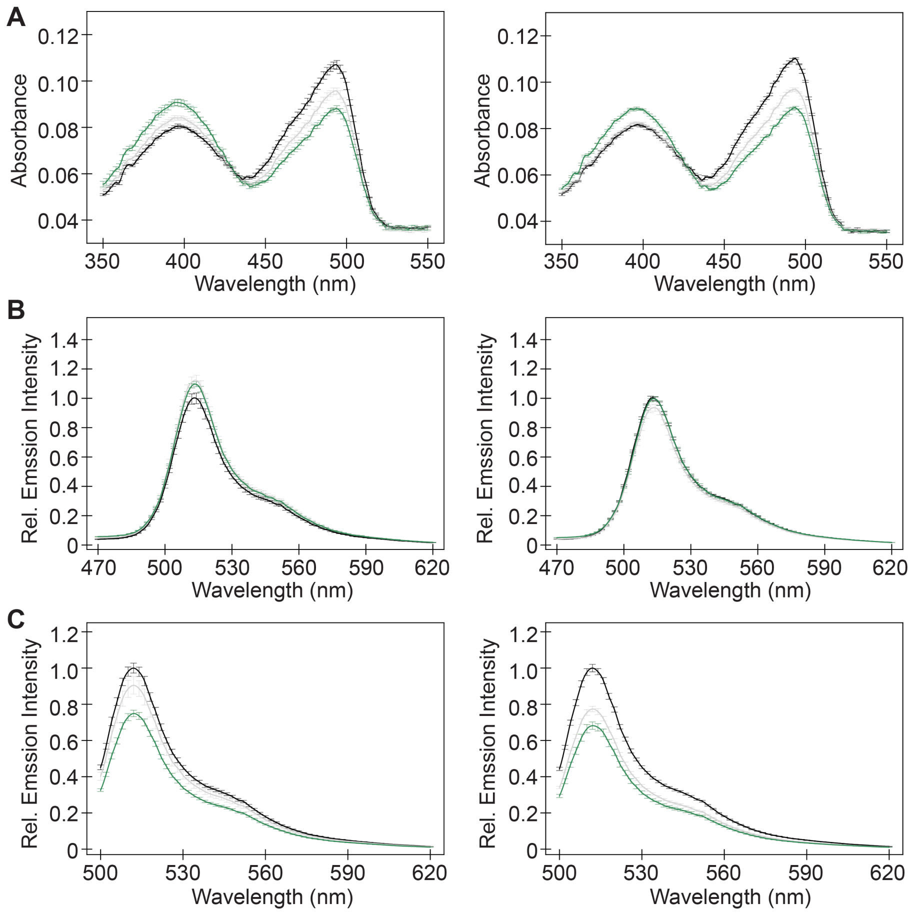


**Figure S33.** Spectroscopic characterization of NitrOFF with chloride. (A) Absorbance and emission spectra of ~4 μM NitrOFF excited at (B) 400 nm and (C) 480 nm in the presence of 0 (black), 10 (gray), and 100 mM (orange) sodium chloride in 25 mM sodium phosphate buffer at pH 7 with 1 mM sodium chloride. The average of three technical replicates with standard deviation is shown for one of two protein batches in each column.

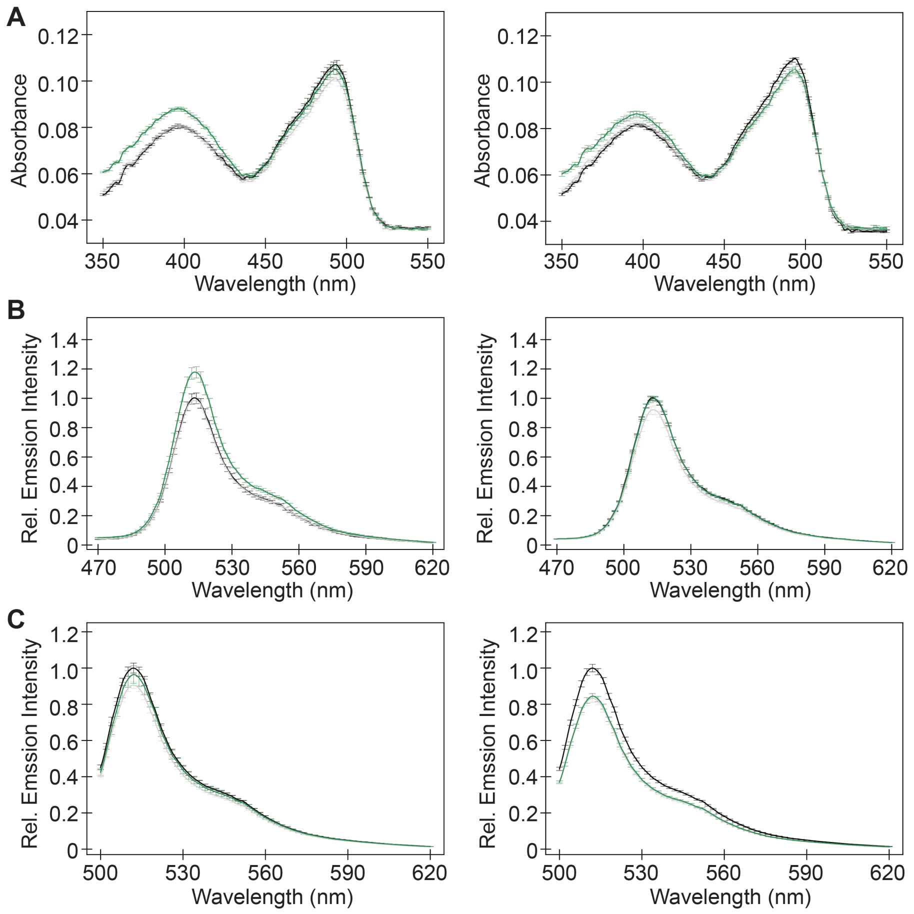


**Figure S34.** Spectroscopic characterization of NitrOFF with gluconate. (A) Absorbance and emission spectra of ~4 μM NitrOFF excited at (B) 400 nm and (C) 480 nm in the presence of 0 (black), 10 (gray), and 100 mM (orange) sodium gluconate in 25 mM sodium phosphate buffer at pH 7 with 1 mM sodium chloride. The average of three technical replicates with standard deviation is shown for one of two protein batches in each column.

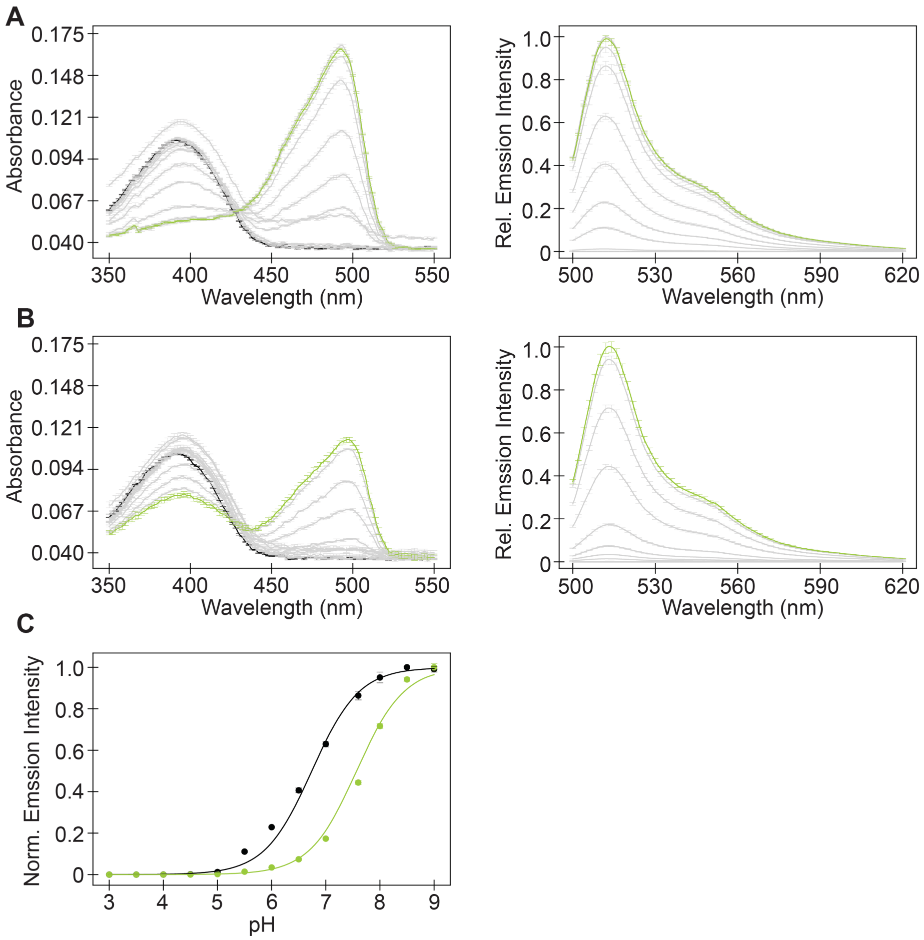


**Figure S35.** Absorbance (left panels) and emission spectra (right panels) of ~4 µM NitrOFF excited at 480 nm in the presence of (A) 0 mM and (B) 0.5 mM nitrate from pH 3 to 9. (C) The normalized emission response at 512 nm from *(A)* and *(B)* is fitted to determine the p$K_a$ of NitrOFF with 0 mM (black) and 0.5 mM (green) nitrate. Data from the first protein batch was measured in triplicate and is plotted as the average with standard deviation.

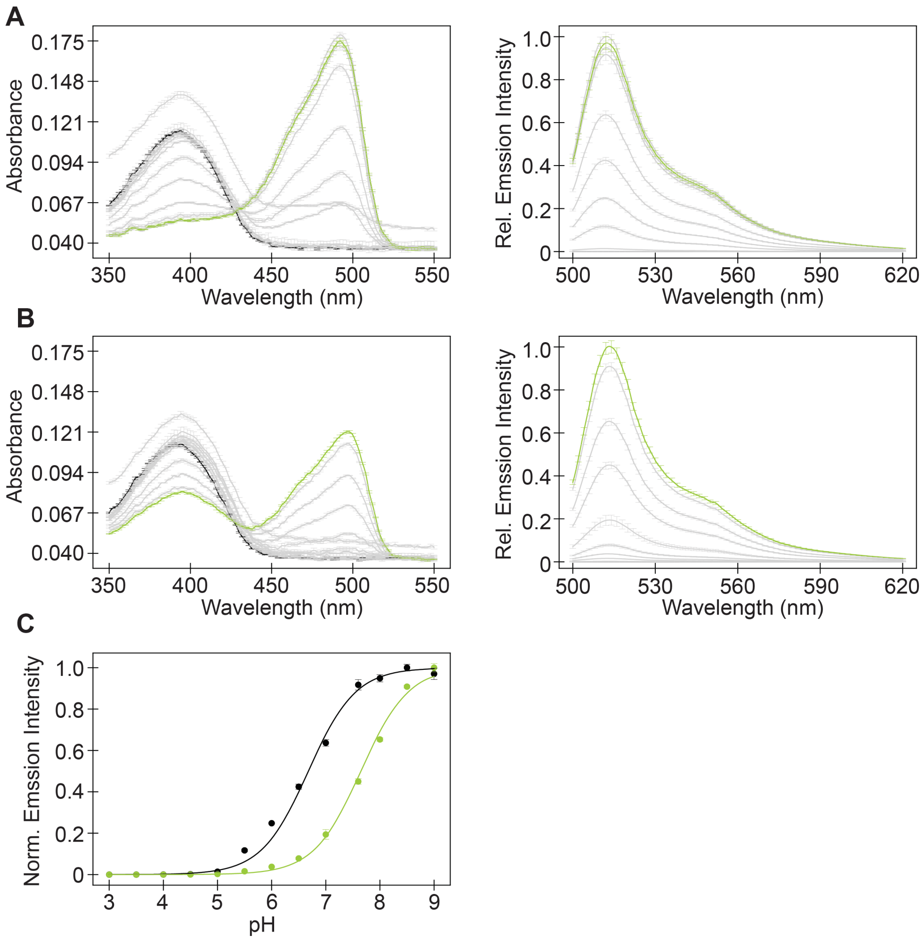


**Figure S36.** Absorbance (left panels) and emission spectra (right panels) of ~4 μM NitOFF excited at 480 nm in the presence of (A) 0 mM and (B) 0.5 mM nitrate from pH 3 to 9. (C) The normalized emission response at 512 nm from *(A)* and *(B)* is fitted to determine the p$K_a$ of NitrOFF with 0 mM (black) and 0.5 mM (green) nitrate. Data from the second protein batch was measured in triplicate and is plotted as the average with standard deviation.

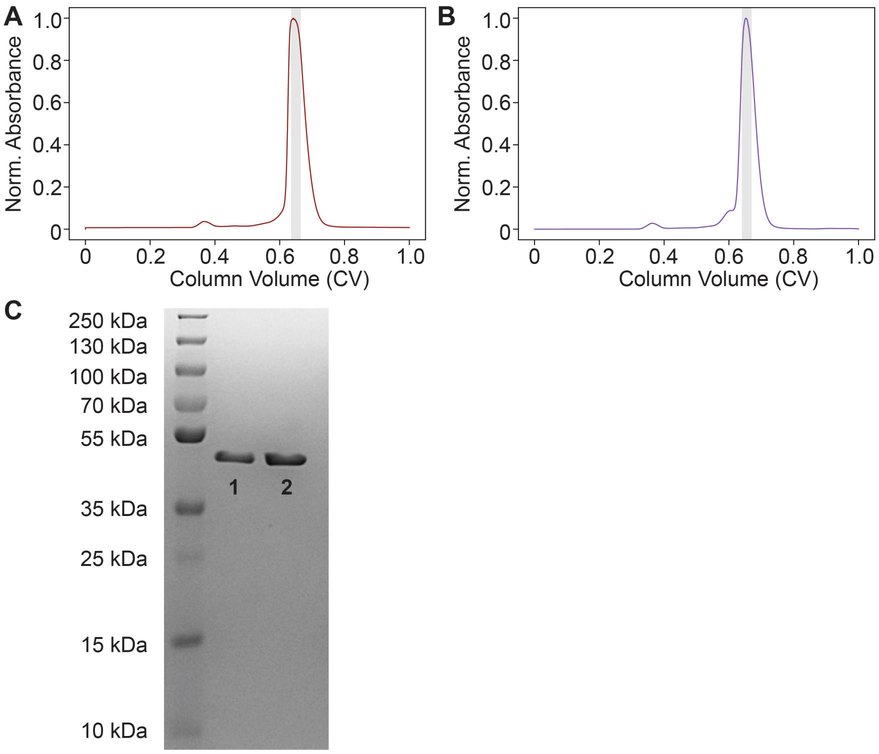


**Figure S37.** Representative size exclusion chromatogram (SEC, normalized absorbance at 280 nm) of (A) apo and (B) bound NitrOFF for protein crystallography. The apo state samples were eluted in 20 mM Tris buffer at pH 7.5 with 150 mM sodium chloride and the bound state samples were eluted in in 20 mM Tris buffer at pH 7.5 with 150 mM $NaNO_3$. For each SEC, the fraction collected for further analysis is shown in the gray area. (C) Representative SDS-PAGE of apo and bound NitrOFF in lanes 1 and 2, respectively. The theoretical molecular weight of NitrOFF is ~44.9 kDa.

**Table S6.** Data collection and refinement statistics for NitrOFF.

| | **NitrOFF (OFF)** | **NitrOFF (ON)** |
|---|---|---|
| **Data Collection** | | |
| Space group | R3H | C2221 |
| Wavelength (Å) | 1.00 | 0.99 |
| **Cell dimensions** | | |
| a,b,c (Å) | 275.6, 275.6, 438.8 | 132.9, 188.3, 188.3 |
| α, β, γ (°) | 90, 90, 120 | 90, 90, 90 |
| Resolution (Å) | 50.0-3.17 (3.22-3.17) | 56.77-3.00 (3.05-3.00) |
| $R_{merge}$/ $R_{pim}$ | 0.170 (0.473)/ 0.102(0.275) | 0.222 (1.023)/ 0.125 (0.582) |
| CC ½ $^{Y}$ | 0.935 (0.686) | 0.977 (0.374) |
| I / σ | 5.3 (1.8) | 7.2 (1.3) |
| Completeness (%) | 99.4 (99.7) | 93.2 (85.9) |
| Redundancy | 1.8 (1.9) | 4.0 (3.9) |
| **Refinement** | | |
| Resolution (Å) | 49.83-3.17 (3.25-3.17) | 56.77-3.00 (3.05-3.00) |
| No. reflections | 209452 (20697) | 48832 (4027) |
| $R_{work}$ | 0.2325 (0.2889) | 0.2368 (0.3108) |
| $R_{free}$ $^{\pm}$ | 0.2746 (0.3609) | 0.2791 (0.3700) |
| **No. atoms** | 54817 | 11729 |
| Protein | 53820 | 11506 |
| Ligand/ion | 758 | 99 |
| Water | 0 | 0 |
| **B-factors (Å$^2$)** | | |
| Protein | 36.1 | 49.61 |
| Ligand/ion | 40.3 | 49.72 |
| Water | N/A | N/A |
| **R.m.s. deviations** | | |
| Bond lengths (Å) | 0.009 | 0.011 |
| Bond angles (°) | 1.53 | 1.48 |
| **Ramachandran plot (%)** | | |
| Favored | 96.36 | 96.48 |
| Allowed | 3.51 | 2.82 |
| Outliers | 0.13 | 0.7 |
| MolProbity score^ | 1.89 (99$^{th}$ percentile) | 1.85 (99$^{th}$ percentile) |

*Values for the corresponding parameters in the outermost shell in parenthesis. $^{Y}CC_{1/2}$ is the Pearson correlation coefficient for a random half of the data, the two numbers represent the lowest and highest resolution shell, respectively. $^{\pm}R_{free}$ is the $R_{work}$ calculated for about 10% of the reflections randomly selected and omitted from refinement. ^MolProbity score is calculated by combining clashscore with rotamer and Ramachandran percentage and scaled based on X-ray resolution. The percentage is calculated with 100th percentile as the best and 0th percentile as the worst among structures of comparable resolution.

**Table S7.** Primers used to subclone the RpHluorin2 gene (insert) into the pcDNA3.1(+) vector (backbone).

| Description | Primer Sequence (5' to 3') |
|---|---|
| Insert Forward | GCTTGCCACCATGAGCAAGGGCGAGG |
| Insert Reverse | GCAGAATTCTTACTTGTACAGC |
| Backbone Forward | GCTGTACAAGTAAGAATTCTGC |
| Backbone Reverse | CCTCGCCCTTGCTCATGGTGGCAAGC |

**Table S8.** Polymerase chain reaction conditions to subclone the RpHluorin2 gene into the pcDNA3.1(+) vector.

**Insert Reaction Conditions**

| Component | Volume |
|---|---|
| Phusion 2X Master Mix | 25 µL |
| 10 ng/µL DNA Template | 1 µL |
| 10 µM Insert Forward Primer | 1 µL |
| 10 µM Insert Reverse Primer | 1 µL |
| Autoclaved Water | 22 µL |
| **Total** | 50 µL |

**Insert Thermocycler Settings**

| Step | Temperature | Time | Number of Cycles |
|---|---|---|---|
| Template Denaturation | 98 °C | 3 min | 1 |
| | 98 °C | 30 s | 30 |
| Annealing | 55 °C | 30 s | |
| Extension | 72 °C | 35 s | |
| Final Extension | 72 °C | 5 min | 1 |
| Storage | 10 °C | ∞ | 1 |

**Backbone Reaction Conditions**

| Component | Volume |
|---|---|
| Phusion 2X Master Mix | 25 µL |
| 10 ng/µL DNA Template | 1 µL |
| 10 µM Backbone Forward Primer | 1 µL |
| 10 µM Backbone Reverse Primer | 1 µL |
| Autoclaved Water | 22 µL |
| **Total** | 50 µL |

**Backbone Thermocycler Settings**

| Step | Temperature | Time | Number of Cycles |
|---|---|---|---|
| Template Denaturation | 98 °C | 3 min | 1 |
| | 98 °C | 30 s | 30 |
| Annealing | 55 °C | 30 s | |
| Extension | 72 °C | 4 min | |
| Final Extension | 72 °C | 10 min | 1 |
| Storage | 10 °C | ∞ | 1 |

```
GCC ACC ATG AGC AAG GGC GAG GAG CTG TTC ACC GGC GTG GTG CCC ATC CTG GTG GAG CTG
Ala Thr Met Ser Lys Gly Glu Glu Leu Phe Thr Gly Val Val Pro Ile Leu Val Glu Leu  20

GAC GGC GAC GTG AAC GGC CAC AAG TTC AGC GTG AGC GGC GAG GGC GAG GGC GAC GCC ACC
Asp Gly Asp Val Asn Gly His Lys Phe Ser Val Ser Gly Glu Gly Glu Gly Asp Ala Thr  40

TAC GGC AAG CTG ACC CTG AAG TTC ATC TGC ACC ACC GGC AAG CTG CCC GTG CCC TGG CCC
Tyr Gly Lys Leu Thr Leu Lys Phe Ile Cys Thr Thr Gly Lys Leu Pro Val Pro Trp Pro  60

ACC CTG GTG ACC ACC CTG AGC TAC GGC GTG CAG TGC TTC AGC CGC TAC CCC GAC CAC ATG
Thr Leu Val Thr Thr Leu Ser Tyr Gly Val Gln Cys Phe Ser Arg Tyr Pro Asp His Met  80

AAG CAG CAC GAC TTC TTC AAG AGC GCC ATG CCC GAG GGC TAC GTG CAG GAG CGC ACC ATC
Lys Gln His Asp Phe Phe Lys Ser Ala Met Pro Glu Gly Tyr Val Gln Glu Arg Thr Ile  100

TTC TTC AAG GAC GAC GGC AAC TAC AAG ACC CGC GCC GAG GTG AAG TTC GAG GGC GAC ACC
Phe Phe Lys Asp Asp Gly Asn Tyr Lys Thr Arg Ala Glu Val Lys Phe Glu Gly Asp Thr  120

CTG GTG AAC CGC ATC GAG CTG AAG GGC ATC GAC TTC AAG GAG GAC GGC AAC ATC CTG GGC
Leu Val Asn Arg Ile Glu Leu Lys Gly Ile Asp Phe Lys Glu Asp Gly Asn Ile Leu Gly  140

CAC AAG CTG GAG TAC AAC TAC AAC GAG CAC CTG GTC TAC ATC ATG GCC GAC AAG CAG AAG
His Lys Leu Glu Tyr Asn Tyr Asn Glu His Leu Val Tyr Ile Met Ala Asp Lys Gln Lys  160

AAC GGC ACC AAG GCC ATC TTC CAG GTG CAC CAC AAC ATC GAG GAC GGC GGC GTG CAG CTG
Asn Gly Thr Lys Ala Ile Phe Gln Val His His Asn Ile Glu Asp Gly Gly Val Gln Leu  180

GCC GAC CAC TAC CAG CAG AAC ACC CCC ATC GGC GAC GGC CCC GTG CTG CTG CCC GAC AAC
Ala Asp His Tyr Gln Gln Asn Thr Pro Ile Gly Asp Gly Pro Val Leu Leu Pro Asp Asn  200

CAC TAC CTG CAC ACC CAG AGC GCC CTG AGC AAG GAC CCC AAC GAG AAG CGC GAC CAC ATG
His Tyr Leu His Thr Gln Ser Ala Leu Ser Lys Asp Pro Asn Glu Lys Arg Asp His Met  220

GTG CTG CTG GAG TTC GTG ACC GCC GCC GGC ATC ACC CAC GGC ATG GAC GAG CTG TAC AAG
Val Leu Leu Glu Phe Val Thr Ala Ala Gly Ile Thr His Gly Met Asp Glu Leu Tyr Lys  240

TAA GAA TTC
End Glu Phe  243
```

**Figure S38.** Nucleotide (top row) and amino acid (bottom row) sequence of the RpHluorin2 construct (black) used for transfecting HEK293 cells. The nucleotide sequence for RpHluorin2 was cloned into the pcDNA3.1(+) vector between the BamHI and EcoRI restriction sites (gray) with a start codon (pink) after BamHI and a stop codon (red) preceding the EcoRI restriction sites.

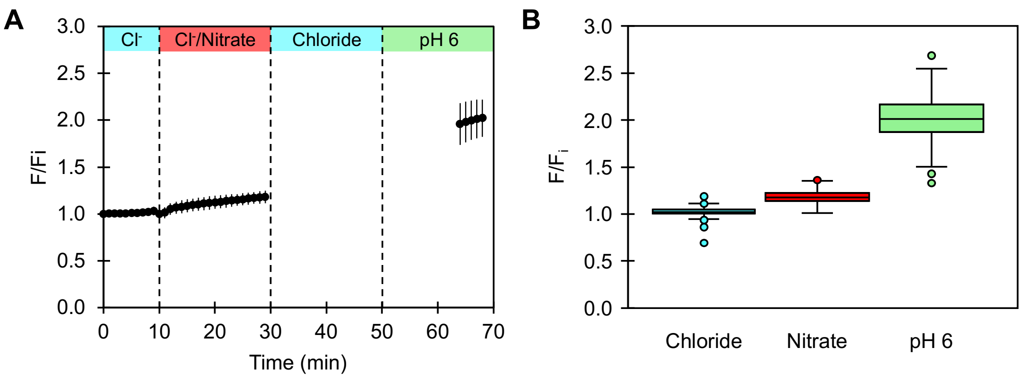


**Figure S39.** (A) The average median fluorescence response ($F/F_i$) with standard deviation of RpHluorin2 from 2 biological replicates with 685 regions of interest for the exchange of 137 mM chloride with 136.5 chloride/0.5 mM nitrate and a pH 6 clamp (Movies S3 – S4). The start time of the perfusion to exchange the buffer is indicated by the vertical dashed line. Prior to the pH clamp phase, all experiments were carried out at pH 7.4. All experiments were carried out at 37 °C. (B) Box plot showing the distribution of the median fluorescence response for all ROIs at the final frame of the baseline ($t$ = 10 min, $F/F_i$ = 1.02 ± 0.05), exchange ($t$ = 29 min, $F/F_i$ = 1.18 ± 0.07), and pH 6 clamp ($t$ = 68 min, $F/F_i$ = 2.02 ± 0.20) conditions shown in *(A)*. The first and third quartiles of each condition are represented by the boxes, maximum and minimum values are represented by the whiskers, and outlier data points are represented by the circles. Abbreviation: $Cl^-$, chloride.

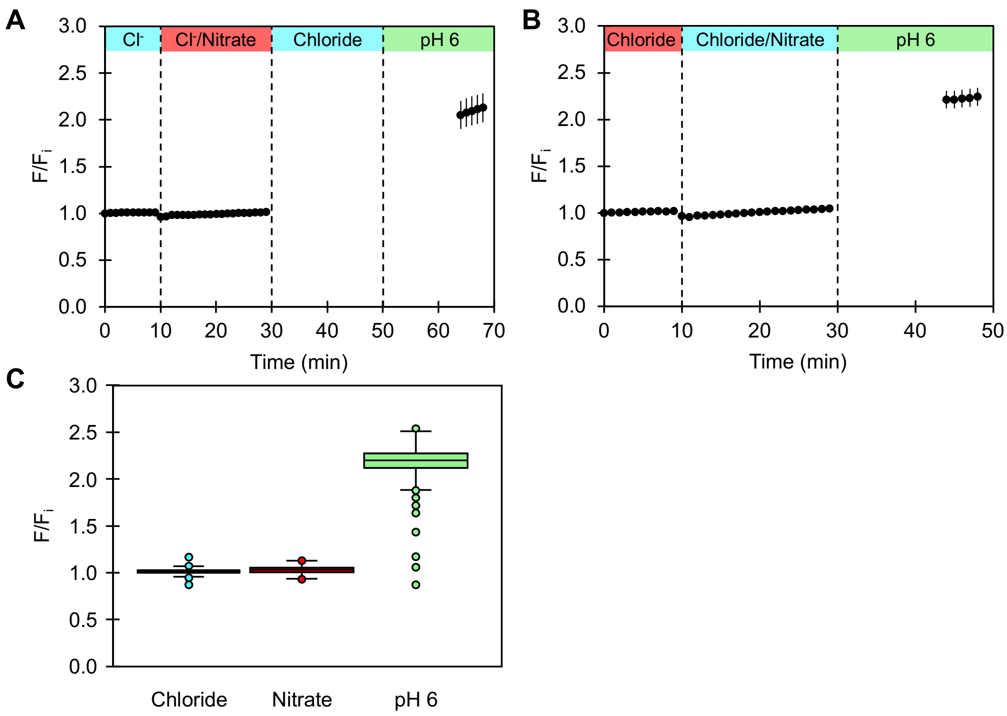


**Figure S40.** The average median fluorescence response ($F/F_i$) with standard deviation of RpHluorin2 from (A) 2 biological replicates with 772 regions of interest (ROI) (Movies S5, S6) and (B) 1 biological replicate with 763 ROIs (Movie S7) for the exchange of 137 mM chloride with 136.75 chloride/0.25 mM nitrate and a pH 6 clamp. The start time of the perfusion to exchange the buffer is indicated by the vertical dashed line. Prior to the pH clamp phase, all experiments were carried out at pH 7.4. All experiments were carried out at 37 °C. (C) Box plot showing the distribution of the median fluorescence response for all ROIs at the final frame of the baseline ($t$ = 10 min, $F/F_i$ = 1.02 ± 0.03), exchange ($t$ = 29 min, $F/F_i$ = 1.03 ± 0.04), and pH 6 clamp ($t$ = 48 min and $t$ = 68 min, $F/F_i$ = 2.19 ± 0.14) conditions shown in *(A)* and *(B)*. The first and third quartiles of each condition are represented by the boxes, maximum and minimum values are represented by the whiskers, and outlier data points are represented by the circles. Abbreviation: $Cl^-$, chloride.

```
GGA TCC GCC ACC ATG GTG TCC AAG GGC GAG GAA CTG TTC ACC GGC GTG GTG CCC ATC CTG
Gly Ser Ala Thr Met Val Ser Lys Gly Glu Glu Leu Phe Thr Gly Val Val Pro Ile Leu   15

GTG GAA CTG GAT GGC GAC GTG AAT GGC CAC AAG TTC AGC GTT TCT GGC GAG GGA GAA GGG
Val Glu Leu Asp Gly Asp Val Asn Gly His Lys Phe Ser Val Ser Gly Glu Gly Glu Gly   35

GAC GCC ACC TAC GGC AAA CTG ACC CTG AAG TTC ATC TGC ACC ACC GGC AAG CTG CCA GTG
Asp Ala Thr Tyr Gly Lys Leu Thr Leu Lys Phe Ile Cys Thr Thr Gly Lys Leu Pro Val   55

CCA TGG CCT ACC CTG GTC ACT ACA CTG ACC TAC GGC GTG CAG TGC TTC AGC CGC TAC CCT
Pro Trp Pro Thr Leu Val Thr Thr Leu Thr Tyr Gly Val Gln Cys Phe Ser Arg Tyr Pro   75

GAC CAT ATG AAG CAA CAC GAC TTC TTT AAA AGC GCT ATG CCT GAA GGC TAC ATT CAG GAG
Asp His Met Lys Gln His Asp Phe Phe Lys Ser Ala Met Pro Glu Gly Tyr Ile Gln Glu   95

AGA ACA ATC TTC TTC AAG GAC GAT GGC AAC TAC AAG ACA AGA GCC GAG GTG AAG TTT GAG
Arg Thr Ile Phe Phe Lys Asp Asp Gly Asn Tyr Lys Thr Arg Ala Glu Val Lys Phe Glu   115

GGC GAT ACA CTG GTG AAC AGA ATC GAG CTG AAA GGC ATT GAC TTT AAA GAG GAC GGA AAC
Gly Asp Thr Leu Val Asn Arg Ile Glu Leu Lys Gly Ile Asp Phe Lys Glu Asp Gly Asn   135

ATC CTG GGA CAC AAG CTG GTG TAC AAC CTG GAC AAC GTG ATC GTC AGC GAC TAC TTC GAC
Ile Leu Gly His Lys Leu Val Tyr Asn Leu Asp Asn Val Ile Val Ser Asp Tyr Phe Asp   155

TAT CAG GAC GCC CTG GAT GAA ATC AGA GAG ACA GAA AAG TTC GAC TTC GCC GCT ATC GCC
Tyr Gln Asp Ala Leu Asp Glu Ile Arg Glu Thr Glu Lys Phe Asp Phe Ala Ala Ile Ala   175

CTG CCC GAA GAT GGA CCT CAC TCC GCC GTG ATC AAA TGG AAG TAC GCC AGC GGC AAC ATC
Leu Pro Glu Asp Gly Pro His Ser Ala Val Ile Lys Trp Lys Tyr Ala Ser Gly Asn Ile   195

GAC TAC AGA TAC CGG ATG ATC GTG CTT AGA CCT GGC GAA GGC CTG GCT GGC CTG GTG ATC
Asp Tyr Arg Tyr Arg Met Ile Val Leu Arg Pro Gly Glu Gly Leu Ala Gly Leu Val Ile   215

CGG ACA GGC AGC AGA AAA ATC GTC GAG GAC GTG GAC ACC GAG CTG AGC CAG AAC GAC AAG
Arg Thr Gly Ser Arg Lys Ile Val Glu Asp Val Asp Thr Glu Leu Ser Gln Asn Asp Lys   235

CTG GGT TGT CCT ATC GTG CTG TCT GAG GCC CTG ACC GCC CTG GTG GCC ATC CCC CTG TGG
Leu Gly Cys Pro Ile Val Leu Ser Glu Ala Leu Thr Ala Leu Val Ala Ile Pro Leu Trp   255

AAA AAC AAC CGG GTG TAC GGC GCC CTG CTG CTC GGC CAG CGG GAA GGA AGA CCC CTC CCT
Lys Asn Asn Arg Val Tyr Gly Ala Leu Leu Leu Gly Gln Arg Glu Gly Arg Pro Leu Pro   275

GAG GGG TCT ACA ACC TTC CGG GTG AAC CAG AGA CTG GGC AGC TAC ACC GAC GAG ATC AAC
Glu Gly Ser Thr Thr Phe Arg Val Asn Gln Arg Leu Gly Ser Tyr Thr Asp Glu Ile Asn   295

AAG CAG GGC AAT GTG TAC ATC AAG GCT GAT AAG CAG AAG AAT GGC ATC AAG GCC AAC TTC
Lys Gln Gly Asn Val Tyr Ile Lys Ala Asp Lys Gln Lys Asn Gly Ile Lys Ala Asn Phe   315

AAG ATC AGA CAC AGC ATC GAG GAT GGC GGA GTT CAG CTG GCC TAC CAC TAT CAG CAG AAT
Lys Ile Arg His Ser Ile Glu Asp Gly Gly Val Gln Leu Ala Tyr His Tyr Gln Gln Asn   335

ACC CCT ATC GGC GAT GGC CCT GTG CTG CTG CCC GAT GAC CAC TAC CTG AGC GTG CAA AGC
Thr Pro Ile Gly Asp Gly Pro Val Leu Leu Pro Asp Asp His Tyr Leu Ser Val Gln Ser   355

GAA CTG TCT AAG GAC CCT AAC GAG AAG CGG GAC CAC ATG GTC CTG CTG GAG TTC GTG ACA
Glu Leu Ser Lys Asp Pro Asn Glu Lys Arg Asp His Met Val Leu Leu Glu Phe Val Thr   375

GCC GCG GGC ATC ACC CTG GGC ATG GAC GAG CTG TAC AAG TGA GAA TTC
Ala Ala Gly Ile Thr Leu Gly Met Asp Glu Leu Tyr Lys End Glu Phe   391
```

**Figure S41**. Nucleotide (top row) and amino acid (bottom row) sequence of the NitrOFF construct (green) used for transfecting HEK293 cells. The nucleotide sequence for NitrOFF was cloned into the pcDNA3.1(+) vector between the BamHI and EcoRI restriction sites (gray) with a Kozak sequence (pink) preceding the start codon (blue) and a stop codon (red) preceding the EcoRI restriction site.

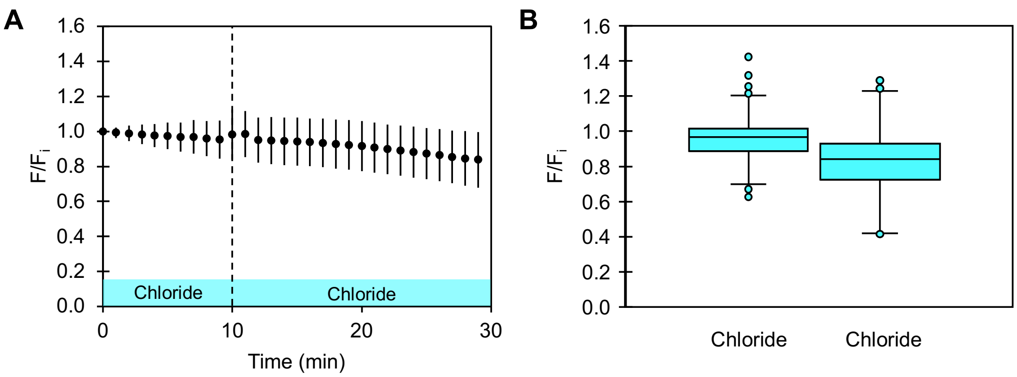


**Figure S42.** The average median fluorescence response ($F/F_i$) with standard deviation of NitrOFF from 3 biological replicates with 624 regions of interest for the exchange of 137 mM chloride with 137 chloride (Movies S8 – S10). The start time of the perfusion to exchange the buffer is indicated by the vertical dashed line. All experiments were carried out at pH 7.4, 37 °C. (B) Box plot showing the distribution of the median fluorescence response for all ROIs at the final frame of the baseline ($t$ = 10 min, $F/F_i$ = 0.95 ± 0.14) and the exchange ($t$ = 29 min, $F/F_i$ = 0.84 ± 0.16) conditions shown in *(A)*. The first and third quartiles of each condition are represented by the boxes, maximum and minimum values are represented by the whiskers, and outlier data points are represented by the circles.

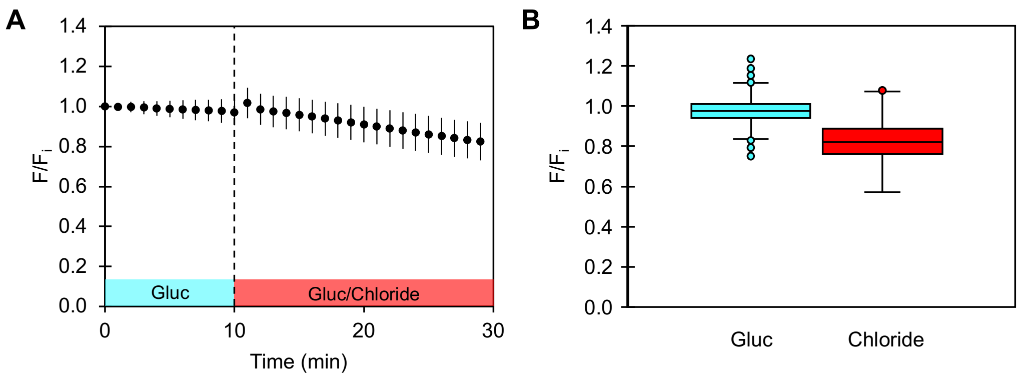


**Figure S43.** The average median fluorescence response ($F/F_i$) with standard deviation of NitrOFF from 3 biological replicates with 2384 regions of interest for the exchange of 137 mM gluconate with 137 chloride (Movies S11 – S13). The start time of the perfusion to exchange the buffer is indicated by the vertical dashed line. All experiments were carried out at pH 7.4, 37 °C. (B) Box plot showing the distribution of the median fluorescence response for all ROIs at the final frame of the baseline ($t$ = 10 min, $F/F_i$ = 0.98 ± 0.06) and the exchange ($t$ = 29 min, $F/F_i$ = 0.82 ± 0.09) conditions shown in *(A)*. The first and third quartiles of each condition are represented by the boxes, maximum and minimum values are represented by the whiskers, and outlier data points are represented by the circles. Abbreviation: Gluc, gluconate.

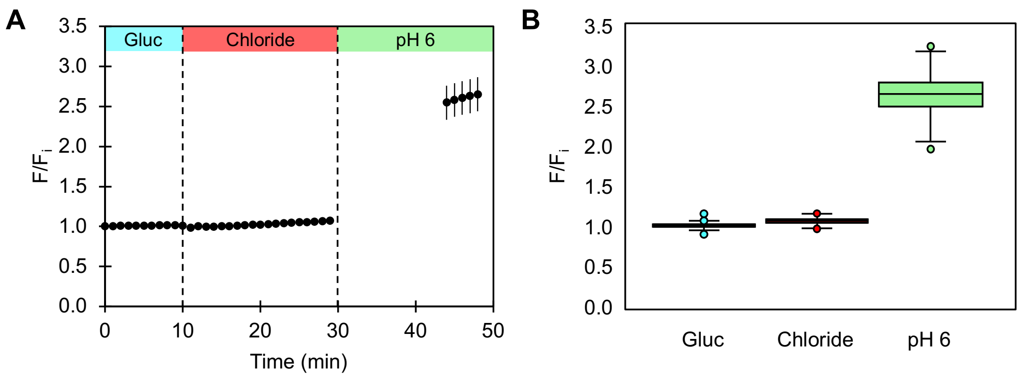


**Figure S44.** (A) The average median fluorescence response ($F/F_i$) with standard deviation of RpHluorin2 from 3 biological replicates with 1308 regions of interest (ROI) for the exchange of 137 mM gluconate with 137 chloride and a pH 6 clamp (Movies S14 – S16). The start time of the perfusion to exchange the buffer is indicated by the vertical dashed line. All experiments, prior to the pH clamp phase, were carried out at pH 7.4. All experiments were carried out at 37 °C. (B) Box plot showing the distribution of the median fluorescence response for all ROIs at the final frame of the baseline ($t$ = 10 min, $F/F_i$ = 1.01 ± 0.02), exchange ($t$ = 29 min, $F/F_i$ = 1.07 ± 0.04), and pH 6 clamp ($t$ = 48 min, $F/F_i$ = 2.65 ± 0.21) conditions shown in *(A)*. The first and third quartiles of each condition are represented by the boxes, maximum and minimum values are represented by the whiskers, and outlier data points are represented by the circles. Abbreviation: Gluc, gluconate.

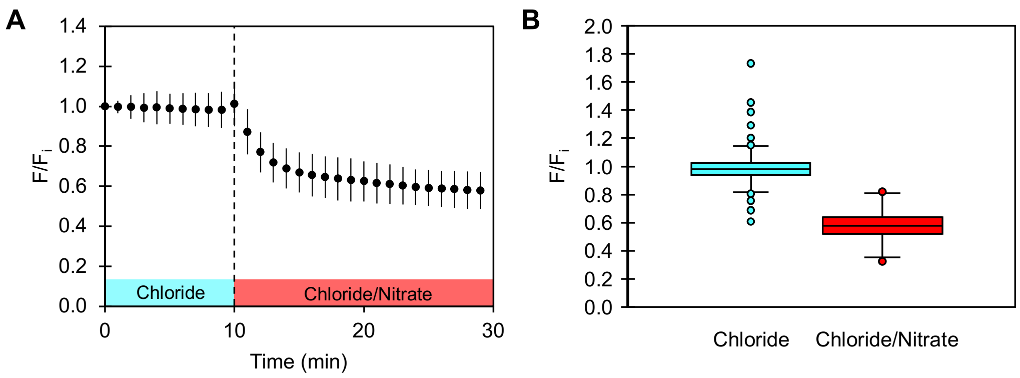


**Figure S45.** The average median fluorescence response ($F/F_i$) with standard deviation of NitrOFF from 3 biological replicates with 664 regions of interest for the exchange of 137 mM chloride with 136.75 chloride/0.25 mM nitrate (Movies S17 – S19). The start time of the perfusion to exchange the buffer is indicated by the vertical dashed line. All experiments were carried out at pH 7.4 at 37 °C. (B) Box plot showing the distribution of the median fluorescence response for all ROIs at the final frame of the baseline ($t$ = 10 min, $F/F_i$ = 1.01 ± 0.11) and the exchange ($t$ = 29 min, $F/F_i$ = 0.58 ± 0.09) conditions shown in *(A)*. The first and third quartiles of each condition are represented by the boxes, maximum and minimum values are represented by the whiskers, and outlier data points are represented by the circles.

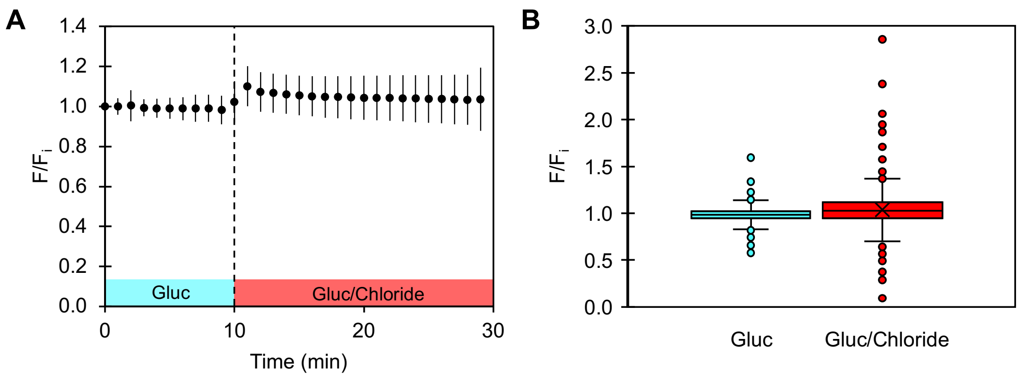


**Figure S46.** The average median fluorescence response ($F/F_i$) with standard deviation of NitrOFF from 3 biological replicates with 2129 regions of interest for the exchange of 137 mM gluconate/0.2% DMSO with 136.75 gluconate/0.25 mM chloride/0.2% DMSO (Movies S20 – S22). The start time of the perfusion to exchange the buffer is indicated by the vertical dashed line. All experiments were carried out at pH 7.4 at 37 °C. (B) Box plot showing the distribution of the median fluorescence response for all ROIs at the final frame of the baseline ($t$ = 10 min, $F/F_i$ = 0.98 ± 0.07) and the exchange ($t$ = 29 min, $F/F_i$ = 1.04 ± 0.16) conditions shown in *(A)*. The first and third quartiles of each condition are represented by the boxes, maximum and minimum values are represented by the whiskers, and outlier data points are represented by the circles. Abbreviations: DMSO, dimethyl sulfoxide; Gluc, gluconate.

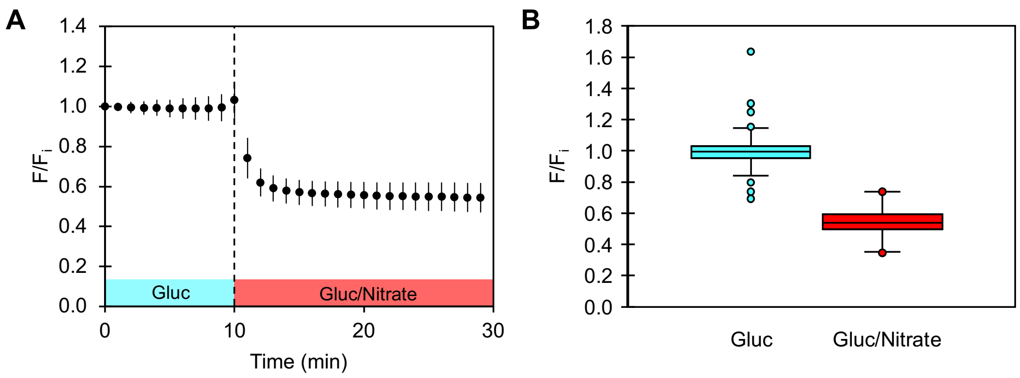


**Figure S47.** The average median fluorescence response ($F/F_i$) with standard deviation of NitrOFF from 3 biological replicates with 1414 regions of interest for the exchange of 137 mM gluconate/0.2% DMSO with 136.75 gluconate/0.25 mM nitrate/0.2% DMSO (Movies S23 – S25). The start time of the perfusion to exchange the buffer is indicated by the vertical dashed line. All experiments were carried out at pH 7.4 at 37 °C. (B) Box plot showing the distribution of the median fluorescence response for all ROIs at the final frame of the baseline ($t$ = 10 min, $F/F_i$ = 0.99 ± 0.07) and the exchange ($t$ = 29 min, $F/F_i$ = 0.54 ± 0.07) conditions shown in *(A)*. The first and third quartiles of each condition are represented by the boxes, maximum and minimum values are represented by the whiskers, and outlier data points are represented by the circles. Abbreviations: DMSO, dimethyl sulfoxide; Gluc, gluconate.

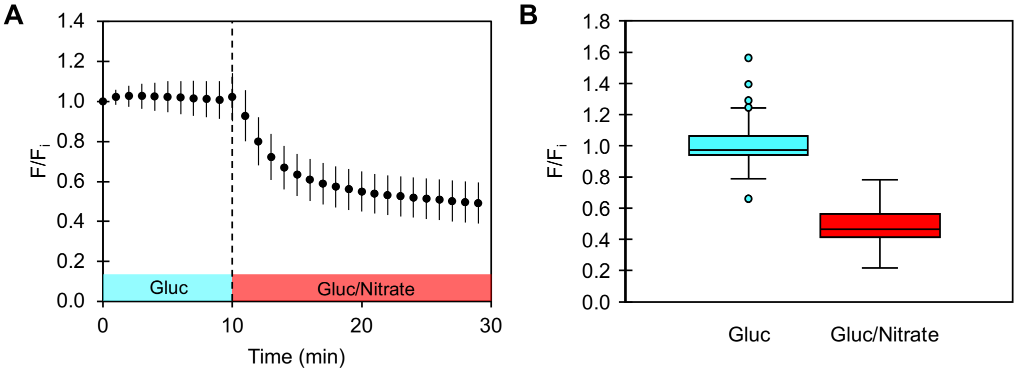


**Figure S48.** The average median fluorescence response ($F/F_i$) with standard deviation of NitrOFF from 3 biological replicates with 1823 regions of interest for the exchange of 137 mM gluconate/0.1 mM NPPB with 136.75 gluconate/0.25 mM chloride/0.1 mM NPPB (Movies S26 – S28). The start time of the perfusion to exchange the buffer is indicated by the vertical dashed line. All experiments were carried out at pH 7.4 at 37 °C. (B) Box plot showing the distribution of the median fluorescence response for all ROIs at the final frame of the baseline ($t$ = 10 min, $F/F_i$ = 1.01 ± 0.09) and the exchange ($t$ = 29 min, $F/F_i$ = 0.49 ± 0.10) conditions shown in *(A)*. The first and third quartiles of each condition are represented by the boxes, maximum and minimum values are represented by the whiskers, and outlier data points are represented by the circles. Abbreviations: Gluc, gluconate, NPPB, 5-nitro-2-(3-phenylpropylamino)benzoic acid.

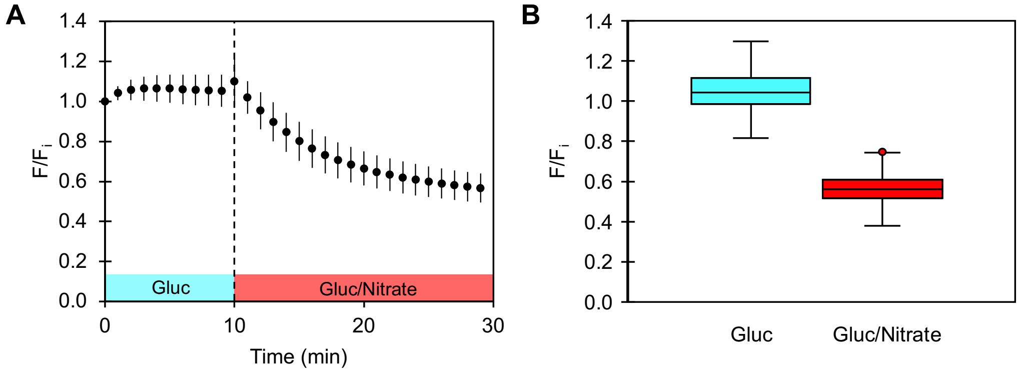


**Figure S5490.** The average median fluorescence response ($F/F_i$) with standard deviation of NitrOFF from 3 biological replicates with 1483 regions of interest for the exchange of 137 mM gluconate/0.2 mM NPPB with 136.75 gluconate/0.25 mM nitrate/0.2 mM NPPB (Movies S29 – S31). The start time of the perfusion to exchange the buffer is indicated by the vertical dashed line. All experiments were carried out at pH 7.4 at 37 °C. (B) Box plot showing the distribution of the median fluorescence response for all ROIs at the final frame of the baseline ($t$ = 10 min, $F/F_i$ = 1.05 ± 0.08) and the exchange ($t$ = 29 min, $F/F_i$ = 0.57 ± 0.08) conditions shown in *(A)*. The first and third quartiles of each condition are represented by the boxes, maximum and minimum values are represented by the whiskers, and outlier data points are represented by the circles. Abbreviations: Gluc, gluconate, NPPB, 5-nitro-2-(3-phenylpropylamino)benzoic acid.

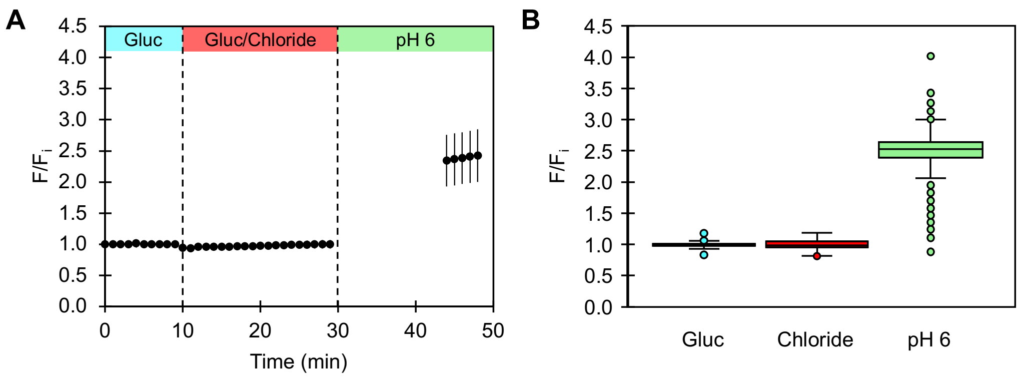


**Figure S50.** (A) The average median fluorescence response ($F/F_i$) with standard deviation of RpHluorin2 from 3 biological replicates with 1087 regions of interest for the exchange of 137 mM gluconate/0.2% DMSO with 136.75 gluconate/0.25 mM chloride/0.2% DMSO, and a pH 6 clamp (Movies S32 – S34). The start time of the perfusion to exchange the buffer is indicated by the vertical dashed line. All experiments, prior to the pH clamp phase, were carried out at pH 7.4. All experiments were carried out at 37 °C. (B) Box plot showing the distribution of the median fluorescence response for all ROIs at the final frame of the baseline ($t$ = 10 min, $F/F_i$ = 1.00 ± 0.03), exchange ($t$ = 29 min, $F/F_i$ = 1.00 ± 0.07), and pH 6 clamp ($t$ = 48 min, $F/F_i$ = 2.43 ± 0.42) conditions shown in *(A)*. The first and third quartiles of each condition are represented by the boxes, maximum and minimum values are represented by the whiskers, and outlier data points are represented by the circles. Abbreviations: DMSO, dimethyl sulfoxide; Gluc, gluconate.

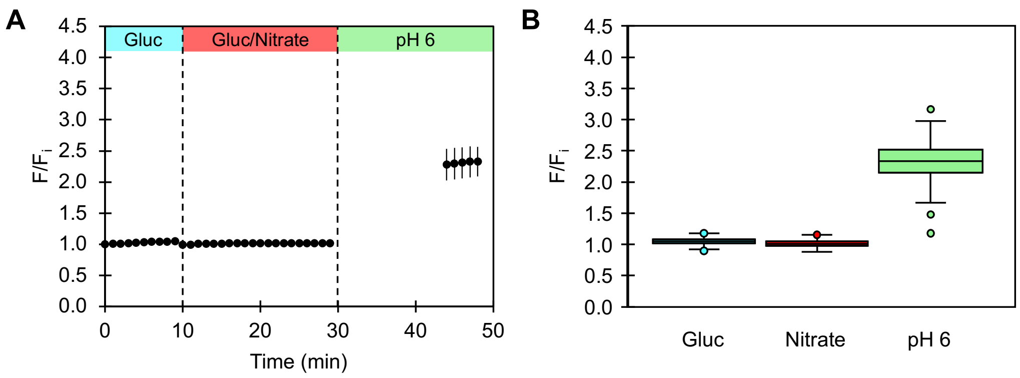


**Figure S51.** (A) The average median fluorescence response ($F/F_i$) with standard deviation of RpHluorin2 from 3 biological replicates with 1033 regions of interest (ROI) for the exchange of 137 mM gluconate/0.2% DMSO with 136.75 gluconate/0.25 mM nitrate/0.2 % DMSO, and a pH 6 clamp (Movies S35 – S37). The start time of the perfusion to exchange the buffer is indicated by the vertical dashed line. All experiments, prior to the pH clamp phase, were carried out at pH 7.4. All experiments were carried out at 37 °C. (B) Box plot showing the distribution of the median fluorescence response for all ROIs at the final frame of the baseline ($t$ = 10 min, $F/F_i$ = 1.05 ± 0.05), exchange ($t$ = 29 min, $F/F_i$ = 1.02 ± 0.05), and pH 6 clamp ($t$ = 48 min, $F/F_i$ = 2.33 ± 0.24) conditions shown in *(A)*. The first and third quartiles of each condition are represented by the boxes, maximum and minimum values are represented by the whiskers, and outlier data points are represented by the circles. Abbreviation: DMSO, dimethyl sulfoxide.

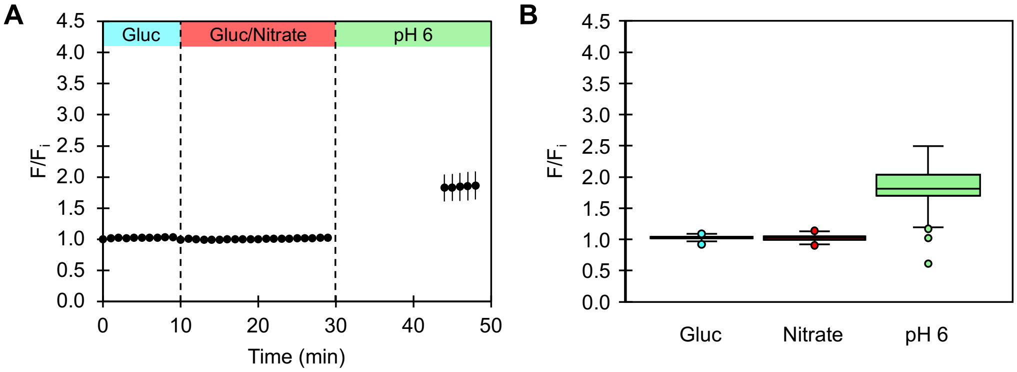


**Figure S52.** (A) The average median fluorescence response (F/F$_i$) with standard deviation of RpHluorin2 from 3 biological replicates with 910 regions of interest for the exchange of 137 mM gluconate/0.1 mM NPPB with 136.75 gluconate/0.25 mM nitrate/0.1 mM NPPB, and a pH 6 clamp (Movies S38 – S40). The start time of the perfusion to exchange the buffer is indicated by the vertical dashed line. Prior to the pH clamp phase, all experiments were carried out at pH 7.4. All experiments were carried out at 37 °C. (B) Box plot showing the distribution of the median fluorescence response for all ROIs at the final frame of each condition shown in *(A)*. The first and third quartiles of the baseline ($t$ = 10 min, F/F$_i$ = 1.03 ± 0.03), exchange ($t$ = 29 min, F/F$_i$ = 1.02 ± 0.04), and pH 6 clamp ($t$ = 48 min, F/F$_i$ = 1.86 ± 0.23) conditions are represented by the boxes, maximum and minimum values are represented by the whiskers, and outlier data points are represented by the circles. Abbreviation: NPPB, 5-nitro-2-(3-phenylpropylamino)benzoic acid.

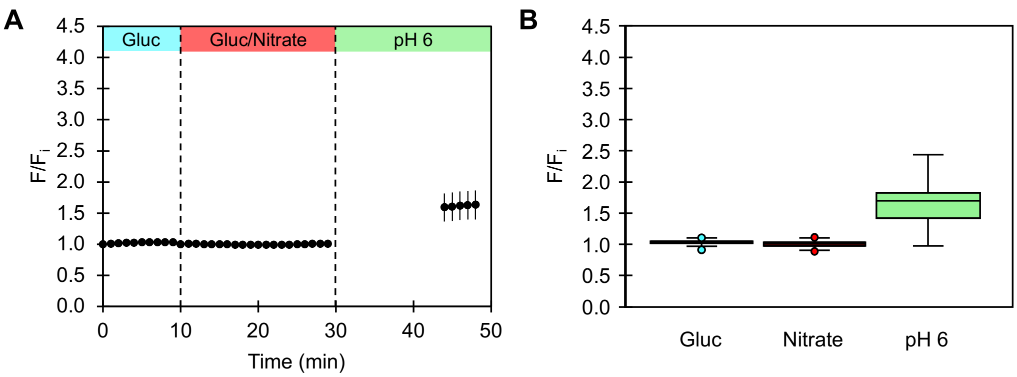


**Figure S53.** (A) The average median fluorescence response ($F/F_i$) with standard deviation of RpHluorin2 from 3 biological replicates with 819 regions of interest for the exchange of 137 mM gluconate/0.2 mM NPPB with 136.75 gluconate/0.25 mM nitrate/0.2 mM NPPB, and a pH 6 clamp (Movies S41 – S43). The start time of the perfusion to exchange the buffer is indicated by the vertical dashed line. Prior to the pH clamp phase, all experiments were carried out at pH 7.4. All experiments were carried out at 37 °C. (B) Box plot showing the distribution of the median fluorescence response for all ROIs at the final frame of each condition shown in *(A)*. The first and third quartiles of the baseline ($t$ = 10 min, $F/F_i$ = 1.03 ± 0.03), exchange ($t$ = 29 min, $F/F_i$ = 1.01 ± 0.04), and pH 6 clamp ($t$ = 48 min, $F/F_i$ = 1.63 ± 0.23) conditions are represented by the boxes, maximum and minimum values are represented by the whiskers, and outlier data points are represented by the circles. Abbreviation: NPPB, 5-nitro-2-(3-phenylpropylamino)benzoic acid.

(S14) Liebschner, D.; Afonine, P. V.; Baker, M. L.; Bunkóczi, G.; Chen, V. B.; Croll, T. I.; Hintze, B.; Hung, L.-W.; Jain, S.; McCoy, A. J.; Moriarty, N. W.; Oeffner, R. D.; Poon, B. K.; Prisant, M. G.; Read, R. J.; Richardson, J. S.; Richardson, D. C.; Sammito, M. D.; Sobolev, O. V.; Stockwell, D. H.; Terwilliger, T. C.; Urzhumtsev, A. G.; Videau, L. L.; Williams, C.

J.; Adams, P. D. Macromolecular Structure Determination Using X-Rays, Neutrons and Electrons: Recent Developments in Phenix. *Acta Cryst. D* **2019**, *75*, 861–877.

(S15) Sangare, L.; Chen, W.; Wang, C.; Chen, X.; Wu, M.; Zhang, X.; Zang, J. Structural insights into the conformational change of *Staphylococcus aureus* NreA at *C*-terminus. *Biotechnol. Lett.* **2020**, *42*, 787–795.

(S16) Wu, S.-Y.; Wen, Y.; Serre, N. B. C.; Laursen, C. C. H.; Dietz, A. G.; Taylor, B. R.; Drobizhev, M.; Molina, R. S.; Aggarwal, A.; Rancic, V.; Becker, M.; Ballanyi, K.; Podgorski, K.; Hirase, H.; Nedergaard, M.; Fendrych, M.; Lemieux, M. J.; Eberl, D. F.; Kay, A. R.; Campbell, R. E.; Shen, Y. A Sensitive and Specific Genetically-Encoded Potassium Ion Biosensor for in Vivo Applications across the Tree of Life. *PLoS Biol.* **2022**, *20*, e3001772.

(S17) Emsley, P.; Lohkamp, B.; Scott, W. G.; Cowtan, K. Features and Development of Coot. *Acta Cryst. D* **2010**, *66*, 486–501.

(S18) Afonine, P. V.; Grosse-Kunstleve, R. W.; Echols, N.; Headd, J. J.; Moriarty, N. W.; Mustyakimov, M.; Terwilliger, T. C.; Urzhumtsev, A.; Zwart, P. H.; Adams, P. D. Towards Automated Crystallographic Structure Refinement with Phenix.Refine. *Acta Cryst. D* **2012**, *68*, 352–367.

(S19) Davis, I. W.; Leaver-Fay, A.; Chen, V. B.; Block, J. N.; Kapral, G. J.; Wang, X.; Murray, L. W.; Arendall, W. B., III; Snoeyink, J.; Richardson, J. S.; Richardson, D. C. MolProbity: All-Atom Contacts and Structure Validation for Proteins and Nucleic Acids. *Nucleic Acids Res.* **2007**, *35*, W375–W383.

(S20) Chen, V. B.; Arendall, W. B.; Headd, J. J.; Keedy, D. A.; Immormino, R. M.; Kapral, G. J.; Murray, L. W.; Richardson, J. S.; Richardson, D. C. MolProbity: All-Atom Structure Validation for Macromolecular Crystallography. *Acta Cryst. D* **2010**, *66*, 12–21.

(S21) Williams, C. J.; Headd, J. J.; Moriarty, N. W.; Prisant, M. G.; Videau, L. L.; Deis, L. N.; Verma, V.; Keedy, D. A.; Hintze, B. J.; Chen, V. B.; Jain, S.; Lewis, S. M.; Arendall III, W. B.; Snoeyink, J.; Adams, P. D.; Lovell, S. C.; Richardson, J. S.; Richardson, D. C. MolProbity: More and Better Reference Data for Improved All-Atom Structure Validation. *Protein Sci.* **2018**, *27*, 293–315.

(S22) Mahon, M. J. pHluorin2: An Enhanced, Ratiometric, pH-Sensitive Green Florescent Protein. *Adv. Biosci. Biotechnol.* **2011**, *2*, 132–137.

(S23) Linders, P. T. A.; Ioannidis, M.; ter Beest, M.; van den Bogaart, G. Fluorescence Lifetime Imaging of pH along the Secretory Pathway. *ACS Chem. Biol.* **2022**, *17*, 240–251.

(S24) Galietta, L. J. V.; Haggie, P. M.; Verkman, A. s. Green Fluorescent Protein-Based Halide Indicators with Improved Chloride and Iodide Affinities. *FEBS Lett.* **2001**, *499*, 220–224.

(S25) Liu, K.-H.; Liu, M.; Lin, Z.; Wang, Z.-F.; Chen, B.; Liu, C.; Guo, A.; Konishi, M.; Yanagisawa, S.; Wagner, G.; Sheen, J. NIN-like Protein 7 Transcription Factor Is a Plant Nitrate Sensor. *Science* **2022**, *377*, 1419–1425.

(S26) Demes, E.; Besse, L.; Cubero-Font, P.; Satiat-Jeunemaitre, B.; Thomine, S.; De Angeli, A. Dynamic Measurement of Cytosolic pH and [$NO_3^-$] Uncovers the Role of the Vacuolar Transporter AtCLCa in Cytosolic pH Homeostasis. *Proc. Natl. Acad. Sci. U.S.A.* **2020**, *117*, 15343–15353.

(S27) Ho, C.-H.; Frommer, W. B. Fluorescent Sensors for Activity and Regulation of the Nitrate Transceptor CHL1/NRT1.1 and Oligopeptide Transporters. *eLife* **2014**, *3*, e01917.

(S28) Hidaka, M.; Gotoh, A.; Shimizu, T.; Minamisawa, K.; Imamura, H.; Uchida, T. Visualization of NO3−/NO2− Dynamics in Living Cells by Fluorescence Resonance Energy Transfer (FRET) Imaging Employing a Rhizobial Two-Component Regulatory System. *Journal of Biol. Chem.* **2016**, *291*, 2260–2269.

(S29) Fatima, U.; Ameen, F.; Soleja, N.; Khan, P.; Almansob, A.; Ahmad, A. A Fluorescence Resonance Energy Transfer-Based Analytical Tool for Nitrate Quantification in Living Cells. *ACS Omega* **2020**, *5*, 30306–30314.

(S30) Chen, Y.-N.; Cartwright, H. N.; Ho, C.-H. In Vivo Visualization of Nitrate Dynamics Using a Genetically Encoded Fluorescent Biosensor. *Sci. Adv.* **2022**, *8*, eabq4915.

(S31) Niemann, V.; Koch-Singenstreu, M.; Neu, A.; Nilkens, S.; Götz, F.; Unden, G.; Stehle, T. The NreA Protein Functions as a Nitrate Receptor in the Staphylococcal Nitrate Regulation System. *Journal of Mol. Biol.* **2014**, *426*, 1539–1553.E

(S32) Ji, K.; Pack, E.; Maydew, C.; Alberto, K.; Abeyrathna, S.; Villones, R.; Gull, H.; Meloni, G.; Nielsen, S.; Dodani, S. Probing the Anion Binding Promiscuity of the Soluble Nitrate Sensor NreA from Staphylococcus Carnosus. *ChemRxiv* **2024**. https://doi.org/10.26434/chemrxiv-2024-pmgn5.
